# Whole blood transcriptional responses associated with bacterial burden in pulmonary tuberculosis

**DOI:** 10.64898/2026.01.29.702471

**Authors:** Le Hoang Thanh Nhat, Thai Minh Triet, Hoang Thanh Hai, Le Hong Van, Le Nguyen Hong Thai, Trinh Thi Bich Tram, Do Dang Anh Thu, Dang Thi Minh Ha, Ho Dang Trung Nghia, Guy Thwaites, Nguyen Thuy Thuong Thuong

**Affiliations:** Oxford University Clinical Research Unit, Ho Chi Minh City, Viet Nam; Pham Ngoc Thach Hospital, Ho Chi Minh City, Viet Nam; Hospital for Tropical Diseases, Ho Chi Minh City, Viet Nam; Pham Ngoc Thach University of Medicine, Ho Chi Minh City, Viet Nam; Centre for Tropical Medicine and Global Health, Nuffield Department of Medicine, University of Oxford, Oxford, United Kingdom

**Author notes:** Correspondence should be addressed to: Nguyen Thuy Thuong Thuong Oxford University Clinical Research Unit, Hospital for Tropical Diseases 764 Vo Van Kiet, W.1, Dist.5, Ho Chi Minh City. Equally contributed (Thai Minh Triet, Hoang Thanh Hai).

**Keywords:** Pulmonary TB, bacterial burden, immune response, CNIH4, neutrophils, monocytes, TCR signaling pathway, lung damage

## Abstract

**Background:** Pulmonary TB (PTB) patients present with a wide range of pre-treatment *Mycobacterium tuberculosis* (*Mtb*) burdens, which predict poor treatment outcomes. We sought to identify immune pathways and biomarkers associated with pre-treatment *Mtb* burden.

**Methods:** We conducted whole-blood RNA sequencing in 295 Vietnamese adults with PTB, quantifying bacterial load using GeneXpert Ct values. Weighted gene co-expression network analysis (WGCNA) identified gene modules, pathways, and hub genes associated with *Mtb* burden. Deconvolution analysis assessed contributions of immune cell types. Key findings were validated in independent PTB (n=171) and TB meningitis (TBM, n=281) cohorts, and publicly available animal datasets. We used non-linear regression for variable selection to identify gene predictors of *Mtb* burden and hurdle regression to model *Mtb* loads below the detection limit.

**Findings:** Higher *Mtb* burden correlated with prolonged symptom duration, elevated neutrophil and monocyte counts, and severe lung pathology. WGCNA identified a 1,093-gene module associated with *Mtb* burden, characterized by coordinated innate-adaptive pathway interactions. Within this module, IFN-γ signaling participates in modulating the increase of innate signaling (Toll-like, Nod-like receptors, TNF) and the decrease of adaptive signaling (T- and B-cell receptor) pathways in high-burden patients. These responses were primarily driven by neutrophils and classical monocytes. CNIH4 emerged as the strongest hub-gene and a top predictor of bacterial burden, with consistent validation across independent PTB and TBM cohorts.

**Interpretation:** Our study reveals systemic innate–adaptive immune dynamics underlying bacterial burden in PTB and identifies CNIH4 as a potential biomarker for treatment monitoring as well as a therapeutic target.

**Funding:** National Institute of Health; Wellcome Trust, UK.

**Research in Context:** *Evidence before this study:* We searched PubMed from Jan 1, 2000, to Dec 10, 2025, without language restrictions, for human studies examining the relationship between Mycobacterium tuberculosis (Mtb) burden and host blood transcriptional responses in pulmonary tuberculosis. Search terms were used in combination as follows: (“Tuberculosis, Pulmonary” OR “pulmonary tuberculosis” OR PTB) AND (“Mycobacterium tuberculosis” OR mycobacter*) AND (“bacterial load” OR “bacterial burden” OR “sputum smear” OR “smear grade” OR xpert OR genexpert OR “cycle threshold” OR ct OR “time to positivity” OR TTP OR CFU OR “molecular bacterial load” OR MBLA)) AND (blood OR “whole blood” OR “peripheral blood”) AND (RNA-seq OR “RNA sequencing” OR transcriptom* OR “gene expression” OR microarray). We included studies of active pulmonary tuberculosis that measured quantitative or semi-quantitative bacterial burden and profiled host blood transcriptome-wide responses at baseline, reporting either differential expression by burden strata or associations between bacterial burden and host gene expression. We excluded studies limited to latent tuberculosis, animal or in vitro models, diagnostic or prognostic signature studies without bacterial burden measurement, studies focused on treatment response, and studies using targeted assays without transcriptome-wide profiling. This search identified 12 articles describing blood transcriptional signatures for tuberculosis diagnosis, prognosis, and treatment response. However, only one study directly examined the relationship between pre-treatment bacterial burden and whole-blood transcriptome-wide profiles. That study demonstrated differences in systemic gene expression between patients with higher and lower sputum mycobacterial load and proposed a 20-gene blood signature associated with bacterial burden. However, the analysis was limited by small sample size, lack of pathway-level and cellular interpretation or assessment of correlation between signature with bacterial load.

*Added value of this study:* Our study advances existing evidence by leveraging the wide spectrum of pre-treatment bacterial burden observed in routine clinical populations, quantified using GeneXpert Ct values, and integrating this with whole-blood RNA sequencing in large, well-characterized clinical cohorts. Through network-based transcriptomic analysis, immune cell deconvolution, and non-linear modelling, we identify a bacterial burden–associated gene network characterized by enhanced innate inflammatory signaling and relative suppression of adaptive immune pathways, predominantly driven by neutrophils and classical monocytes and modulated by IFN-γ signaling. Within this network, CNIH4 emerges as a central hub gene and a robust predictor of bacterial burden, with consistent validation across independent pulmonary tuberculosis and tuberculous meningitis cohorts.

*Implications of all the available evidence:* Taken together, the available evidence indicates that host blood transcriptional responses correlate with bacterial burden in pulmonary tuberculosis, but previous studies have provided limited insight into the underlying immune processes. Our findings strengthen the biological link between pre-treatment mycobacterial burden and systemic immune dysregulation, showing that higher bacterial burden is associated with transcriptional state marked by coordinated upregulation of innate immune responses and downregulation of adaptive immune pathways. These results support the use of host transcriptomic profiling as a biologically informative complement to sputum-based measures of bacterial burden and highlight burden-associated immune pathways, particularly CNIH4, as a potential target for treatment monitoring and host-directed therapeutic development.

## INTRODUCTION

Tuberculosis (TB), caused by *Mycobacterium tuberculosis* (*Mtb*), remains a significant global health challenge. In 2023, TB affected 10·8 million people worldwide and caused over 1·25 million deaths, including those with HIV.^1^ Despite advances in TB research, the interplay between immune responses, bacterial killing, and disease severity is still not fully understood.

One key observation is that people with pulmonary TB (PTB) exhibit substantial variability in pre-treatment bacterial burden. In sputum samples, this burden ranges from approximately 10³ to 10⁸ CFU/ml. ^2^ A high pre-treatment *Mtb* burden is a robust prognostic factor for unfavorable treatment outcomes from PTB.^3^ Such pronounced heterogeneity raises an important question: how do differences in host immune responses shape the initial *Mtb* burden? Therefore, deciphering the underlying mechanisms of host response in controlling bacterial burden, and identifying biomarkers associated with this variability are crucial. They may help to improve treatment monitoring and guide the development of host-directed therapies. Such biomarkers and interventions may be particularly important for patients with high *Mtb* burden, as they are more likely to experience unfavorable outcomes. We hypothesized that disparity in bacterial burden arises from differing host immune profiles, particularly in the interaction of innate and adaptive immune pathways before the initiation of anti-TB therapy.

Innate immune sensors detect *Mtb.* These include Toll-like receptors (TLR2, TLR4, and TLR9) and Nod-like receptors (NOD2, NLRP3, and AIM2), which trigger pro-inflammatory cytokines (*e.g.,* TNF-α and IL-1β). These cytokines recruit immune cells to the infection site to control bacterial replication,^4–7^ but their excessive production can cause tissue damage.^8–10^ Adaptive immune response plays an important role in *Mtb* bacterial clearance. Specifically, IFN-γ–producing T cells enhance antimicrobial macrophage activities to kill bacteria^11–16^ (summary in Box 1, Appendix). Neutrophils responding to IFN-γ signaling can further amplify immune responses. However, if dysregulated, they can contribute to tissue destruction, forming lung cavities^17^ promotes bacterial dissemination.^18–21^ Nevertheless, how these mechanisms collectively influence bacterial burden in patients before anti-TB treatment remains poorly understood.

Biological and clinical data, such as gene expression levels or bacterial burden, often exhibit complex interactions and may contain a substantial proportion of measurements below the detection limit. While simple linear regression models are commonly used to study relationships among these variables, they are limited in their ability to capture non-linear patterns (*e.g.*, parabola-shaped or sigmoidal relationships) or to accurately estimate associations. Therefore, more advanced statistical models are needed to improve the analysis of such data.

To investigate the immune response to bacterial burden, we obtained whole-blood RNA sequencing (RNA-seq) transcriptional profiles from a cohort of 295 PTB patients before treatment and integrated these data with clinical features. Key findings were subsequently validated in independent cohorts of 171 PTB patients and 281 TB meningitis (TBM) patients, as well as in cross-species animal models. A key analytical strength of our study was the use of non-linear regression modeling to capture complex biological relationships within gene expression profiles and with bacterial burden. In addition, we applied a hurdle regression model to assess associations between gene expression and bacterial load, addressing the challenge of measurements falling below the detection limit. This integrated approach enabled us to uncover critical immune regulatory pathways linked to *Mtb* burden. It also identified potential blood-based biomarkers for monitoring treatment efficacy and therapeutic targets for enhancing bacterial clearance.

## METHODS

### Study population

We collected clinical data and whole peripheral blood samples for transcriptomic profiling from 295 HIV negative PTB patients, forming our discovery cohort. Additionally, we performed targeted RT-qPCR on samples from 171 randomly selected HIV negative PTB patients, establishing our validation cohort. These patients were recruited as part of a large observational study investigating host and bacterial determinants of PTB treatment outcomes, which enrolled 900 participants from Pham Ngoc Thach Hospital and District TB Units in Ho Chi Minh City, Vietnam, between 2017 and 2020· We excluded HIV-coinfected patients due to the very low prevalence of HIV in this population. All participants had culture-confirmed PTB (either drug-susceptible or newly diagnosed multidrug-resistant TB) and had received less than 7 days of anti-TB treatment at enrollment without clinical evidence of extrapulmonary TB.

We also collected peripheral blood samples for transcriptional profiling from two additional cohorts: 281 TBM patients (including both microbiologically confirmed and clinically diagnosed cases) and 30 healthy controls. For the TBM cohort, we included the first 207 consecutively enrolled HIV-negative patients and randomly selected 74 HIV-positive patients from two randomized clinical trials (the LAST-ACT trial [NCT03100786] for HIV-negative and the ACT-HIV trial [NCT03092817] for HIV-positive). Healthy controls were enrolled from the Hospital for Tropical Diseases in Ho Chi Minh City as part of a prospective study on human resistance to *Mtb* infection; these adults exhibited no TB symptoms or recent TB exposure (within the past two years).

Ethics approval was obtained from the institutional review board at the Hospital for Tropical Diseases, Pham Ngoc Thach Hospital, the Ministry of Health in Vietnam, and the Oxford Tropical Research Ethics Committee, UK. All participants, or their relatives if incapacitated, provided written informed consent.

### Study design and analytical workflow

We used our primary PTB cohort (n = 295) as the discovery cohort to examine the association between whole blood transcriptional profiles and pre-treatment sputum bacterial burden. We then validated these signals in an independent PTB cohort (n = 171), where gene expression was quantified by RT-qPCR, and in the TBM cohort (n = 281, including 74 HIV-positive subjects) for cerebrospinal fluid bacterial burden. Targeted genes for RT-qPCR analysis were selected based on results from the discovery cohort. **Supplementary Figure S9** presents a schematic representation of the cohorts and the data analysis workflow. For our main analysis, we performed gene co-expression network analysis to identify transcriptional modules linked to pre-treatment sputum bacterial burden. To ensure reproducibility, we randomly split the PTB discovery cohort into a training set (n = 147) and a test set (n = 148). We constructed a gene co-expression network using the training set, identified modules associated with bacterial burden, and confirmed these associations in the test set (details in the primary analysis section). Finally, we delineated the related functional pathways and hub genes and compared them across all four cohorts (healthy control, PTB, TBM HIV-negative, and TBM HIV-positive).

### Bacterial burden quantification by GeneXpert

We used the GeneXpert MTB/RIF system (Cepheid, Sunnyvale, CA, USA), an automated real-time PCR test that reports the cycle threshold (Ct) value from five probes targeting the *rpoB* gene region, to quantify *Mycobacterium tuberculosis* (*Mtb*) bacterial burden. For participants with PTB, we collected pre-treatment sputum samples and decontaminated it with 2% NALC-NaOH, followed by resuspension in 2 mL of phosphate-buffered saline. A 500 μL aliquot of the resuspended sediment was mixed with 1·5 mL of GeneXpert sample reagent, vortexed for 30 seconds, incubated at room temperature for 15 minutes, and then transferred to a GeneXpert cartridge for measurement according to standard procedures.^22^ For participants with TBM, we collected about 3-6 mL of cerebrospinal fluid (CSF) and centrifuged it for 20 minutes at 3000×g. The supernatant was removed, and the pellet was resuspended in the remaining 500 μL of CSF. An 200 μL aliquot of the resuspended pellet (about 40% of the total volume) was then mixed with 0·5 mL of phosphate-buffered saline and 1·5 mL of GeneXpert sample reagent. This mixture was vortexed for 30 seconds, incubated at room temperature for 15 minutes, and subsequently transferred to a GeneXpert cartridge for measurement as previously described.^23,24^

The GeneXpert Dx software (version 4·0; Cepheid) provided semiquantitative *Mtb* load results as Ct values, which indicate the number of PCR cycles needed for the signal to reach a defined detection threshold. To estimate the bacterial load for each participant, the average Ct value of the five probes was calculated, excluding any delayed values potentially indicative of rifampicin resistance. In cases where *Mtb* was not detected, a Ct value of 40 was assigned. *Mtb* loads were classified as high (mean Ct value < 16), medium (16 ≤ mean Ct value < 23), low (23 ≤ mean Ct value ≤ 28), and very low (mean Ct value > 28).^25,26^

### Sample processing and RNA-seq

We collected whole blood samples from participants at enrollment and stored them in PAXgene collection tubes at −80°C. We extracted RNA and performed RNA-seq in two batches. In Batch 1 (2020), we processed samples from 207 HIV-negative TBM, 31 HIV-positive TBM, and 295 PTB cases; in Batch 2 (2022), we processed samples from 43 HIV-positive TBM and 30 healthy controls. We isolated RNA using the PAXgene Blood RNA kit (Qiagen, Valencia, CA, USA) following the manufacturer’s instructions, with an additional washing step before eluting the RNA. We digested DNA on the columns using the RNase-free DNase Set (Qiagen, Valencia, CA, USA). We assessed RNA quantity and quality with the Epoch spectrophotometer and evaluated RNA integrity using the Tapestation Eukaryotic RNA Screentape, excluding samples with an RNA integrity number below 4. We sent the samples to the Ramaciotti Centre for Genomics (Sydney, Australia) for RNA-seq. We used one microgram of total RNA as input for each sample with the TruSeq Stranded Total RNA Ribo-zero Globin kit (Illumina). We generated libraries on the Sciclone G3 NGS (Perkin Elmer, Utah, USA) and amplified the cDNA using 11 PCR cycles. Finally, we pooled 75 samples per pool and sequenced them with NovaSeq 6000 S4 reagents at 2×100 bp, generating approximately 30 million reads per sample.^27^

### RNA-seq data quality control and pre-processing

We performed quality control and alignment using an in-house pipeline that we modified from previously published RNA-seq analysis practices ^28,29^ in the Linux command line. Briefly, we analyzed the quality of the sequencing fastq files using FastQC (v0·11·5^30^) and excluded poor-quality samples from further analysis. We trimmed adapters and low-quality bases from the sequence reads using Trimmomatic (v0·36^31^) and then removed duplicated optical reads using the BBMap (v38·79) tool. We downloaded the human reference genome (GRCh38 build 99) from Ensembl and used the STAR aligner (v2·5·2a^32^) to map the reads to it, permitting up to two mismatches per 25 bp segment and up to 20 alignment hits per read. Next, we sorted and indexed the alignment results into BAM format files and generated gene expression counts from the uniquely mapped reads using FeatureCounts (v2·0·0^33^) against the human reference annotation (GRCh38 build 99), which produced raw counts for 60,676 genes.

For the RNA-seq PTB cohort, we extracted a gene expression matrix for 60,676 genes across 295 participants after performing data quality control and processing. We excluded lowly expressed genes (raw counts <10 in 20% of samples, n = 34,402), ribosomal RNA genes (n = 568), and sex-related genes (n = 2,946), leaving 22,760 genes for analysis. We normalized these genes and transformed them into a log_2_ scale using the variance stabilizing transformation (VST) algorithm in the R package DESeq2,^34^ and we used the resulting normalized expression matrix for primary analysis.

For comparisons in TBM or across the three cohorts (PTB, TBM, healthy control), we applied batch correction since we sequenced the samples in two batches, as described above. First, we normalized the gene expression matrix for 60,676 genes across 606 participants using the VST algorithm. Next, we selected the 20,000 most variable genes and removed the batch effect for these genes using the ComBat function in the R package SVA.^35^ We visualized and evaluated the effectiveness of batch correction using principal component analysis (PCA). Before batch correction, the samples clustered according to their sequencing batch, indicating the presence of a batch effect. After correction, the samples from batch 1 and batch 2 showed substantial overlap, especially in the five repeated measurements, indicating that we successfully removed the batch effects (**Supplementary Figure S10**). Finally, we extracted the normalized data for the genes of interest and conducted comparisons using this data.

### Validation of hub genes by Microfluidic multiplex RT-qPCR

We collected whole blood samples, stored them, and performed RNA extraction as described in the previous section. We then evaluated the expression of housekeeping genes (GAPDH and TMBIM6) and other hub genes by microfluidic RT-qPCR using the Biomark™ 48·48 Complete Bundle with Delta Gene™ Assays and the BioMark™ HD system (Fluidigm Corporation, South San Francisco, CA, USA) following the manufacturer’s instructions with some optimized modifications. Briefly, we reverse transcribed 2 µL of total RNA (50 ng/µL) to cDNA. Next, we performed 14 cycles of specific target amplification (STA) on the cDNA, treated the STA products with Exonuclease I (New England Biolabs, Ipswich, MA, USA), and diluted them 20-fold. We used SsoFast™ EvaGreen® Supermix with Low ROX (Bio-Rad Laboratories, Hercules, CA, USA) in the RT-qPCR reaction before applying the samples to the IFC Controllers MX and the BioMark™ HD system (Fluidigm Corporation, South San Francisco, CA, USA). Finally, we normalized the cycle threshold (CT) values of the target hub genes by subtracting the CT value of GAPDH before analysis.^27^

### Re-analysis experimental animal model data

We investigated the relationship between bacterial burden and the expression of immune-related genes (TLR4, TLR6, AIM2, NOD2, STAT1, IL-1β, PLC-γ1, ZAP70, and CNIH4) in lung tissue by downloading and reanalyzing a published dataset^36^ generated from *Mtb*-infected mice and macaques. The original study provided RNA-seq data normalized to FPKM (fragments per kilobase of transcript per million mapped reads). In that study, the investigators infected 39 mice (10 naïve, 13 controllers, and 16 progressors) with approximately 100 CFU of *Mtb* strain HN878 and exposed 24 macaques (4 naïve, 4 controllers, and 8 progressors) to *Mtb* strain CDC1551. They then harvested lung tissue from each animal, performed RNA-seq, and quantified the bacterial burden by CFU plating and counting method. They defined distinct criteria for each animal to classify into controller and progressor. In mice, controller and progressor group are defined based on “TB severity score” assessed at euthanasia (log10 bacterial burden of the lung tissue sample × log10 percentage lung inflammation, by area), using thresholds of <6·5 for controllers and >8·5 for progressors. In macaque, asymptomatic TB cases are defined as controllers, while those exhibited clinical symptoms and TB disease are defined as progresses.

### Statistical analysis

#### Descriptive analysis

We summarized baseline patient characteristics and lung damage phenotype for PTB patients—stratified by *Mtb* burden—and provided reference summaries for healthy controls. We reported binary variables as proportions and continuous variables as medians with 1^st^/3^rd^ interquartile. To evaluate the association between *Mtb* burden (log₂ Ct value) and clinical/immunological variables, we fitted univariate linear regression models with *Mtb* burden as the outcome and each characteristic as a covariate. For the two chest X-ray phenotypes, we used logistic regression to model the association between Mtb burden and cavity, and beta regression to quantify the relationship between *Mtb* burden and the proportion of lung damage

#### Clustering analysis

We performed hierarchical clustering using Ward’s minimum variance method as implemented in the R package *ComplexHeatmap*.

#### Association analysis

We visualized relationships between any pair of continuous variables using scatterplots and assessed their *linear associations* by computing Pearson correlation coefficients (*r*) with p-values derived via Fisher’s z-transformation. For correlation analysis of all 17 deconvoluted cell-subtype proportions, we employed Spearman correlation to account for their inherently skewed distributions, capturing monotonic non-linear associations while minimizing sensitivity to outliers.

To explore broader *non-linear relationships* both among gene expression profiles and between gene expression and *Mtb* burden, we employed generalized additive models (GAMs) implemented in the *mgcv* R package. These models fit penalized smoothing splines, optimized via cross-validation, to capture complex biological patterns such as U-shaped or sigmoidal associations. The penalization approach priorities simpler, approximately linear relationships unless sufficient evidence supports non-linearity, consistent with the principle of parsimony of statistical modeling. Model fit was assessed using the *adjusted coefficient of determination* 𝑅_adj_^2^ , which quantifies the proportion of variance explained after accounting for model complexity.

To account for zero-inflated distribution of sputum and CSF bacterial load data, arising from a substantial proportion of patients with undetectable *Mtb* levels in the validation cohort, we applied a hurdle regression model. This two-part modeled separately (1) the probability of bacterial detection (binary component) and (2) the conditional mean among detectable samples (continuous component). In particular, we modeled the transformed Ct value (Y = log₂(40/Ct)) as the outcome and *targeted gene (such as CNIH4)* expression as the covariate using the Bayesian R package *brms*. With this transformation, an undetectable *Mtb* burden (Ct = 40) corresponds to Y = 0, allowing us to simultaneously model the binary component (Y = 0 vs. Y > 0) and the continuous, log-transformed component. We then back-transformed the model estimates to the original Ct scale to report mean values and facilitate biological interpretation. We reported Wald-type p-values from both model components, namely the p-values 𝑝_𝑚𝑢_ for continuous (non-hurdle) and 𝑝_ℎ𝑢_ for binary (hurdle) parts. To assess the overall evidence of association between gene expression and bacterial burden refers to 𝑝_𝑜𝑣𝑒𝑟𝑎𝑙𝑙_ , we combined these two p-values using Simes’ procedure,^37^ which controls the family-wise error rate across the two correlated hypotheses, namely, that gene expression may influence either the odds of undetectable burden (𝑝_ℎ𝑢_) or the magnitude of bacterial burden among *Mtb* detected cases (𝑝_𝑚𝑢_).

### Primary analysis

We normalized RNA-seq data from whole blood of 295 individuals, which included expression profiles for 22,760 genes, and selected the most variable genes using an unsupervised filtering method based on the inverse gamma distribution implemented in the R package *CEMiTool* (v3·16). This step yielded 5,849 genes. To ensure robust findings, we randomly split the PTB cohort into a training set (n = 147) and a test set (n = 148). We constructed a gene co-expression network using the training set and performed preservation analysis in the test set.

#### Weighted gene co-expression network analysis (WGCNA)

We performed WGCNA on the 5,849 most variable genes from the PTB patients in the training set using the R package *WGCNA* to identify gene clusters (modules) in the host transcriptomic co-expression network. First, we computed pairwise Pearson correlation coefficients between all gene pairs to form a similarity matrix. We then analyzed the network topology to determine the optimal soft-thresholding power, finding that 𝛽 = 5 achieved a scale-free fit index of 𝑅² = 0 · 9 (**Supplementary Figure S1A**). Using this value, we constructed an unsigned adjacency matrix (𝐴 = 𝑎_𝑖𝑗_ , where 𝑎ᵢⱼ = |𝑐𝑜𝑟(𝑔𝑒𝑛𝑒ᵢ, 𝑔𝑒𝑛𝑒ⱼ)| ) and computed the topological overlap matrix (TOM) to capture the shared connectivity among genes. Next, we performed hierarchical clustering on the genes using the TOM-based dissimilarity matrix defined as (1 − TOM) with *average* linkage and grouped them into modules using a dynamic tree cut algorithm^38^ with a cut height of 0·975, a minimum module size of 30 genes, and a module merging threshold of 0·25. This approach identified 18 gene co-expression modules, each containing between 44 and 1,483 genes (**Supplementary Figures S1C-D**). Finally, we summarized each module’s expression profile by its eigengene—defined as the first principal component of the module’s expression matrix obtained via singular value decomposition—for use in downstream analyses.

#### Preservation analysis

We evaluated the reproducibility of the identified modules and overall network structure by performing a network module preservation analysis in the test set (n = 148) using the WGCNA function *modulePreservation*. This function randomly permuted the module labels from the training set network and computed seven key module statistics—capturing both density and connectivity measures—in the test set. We repeated this procedure 1,000 times to generate null distributions (mean and standard deviation) for each statistic and then calculated a Z-score for each statistic by subtracting the null mean from the observed value and dividing by the null standard deviation. WGCNA combined these Z-scores into overall Z_density_ and Z_connectivity_ scores; modules with both scores greater than 10 were considered well preserved and retained for further analysis.^39^

#### Module-trait association analysis and hub gene identification

We then assessed the association between each module and the primary outcome, *Mtb* burden, by fitting simple linear regression models with the module eigengene (the first principal component of the module’s expression matrix) as covariate. We corrected for false discovery rate (FDR) using the Benjamini-Hochberg procedure^40^ and considered modules associated with *Mtb* burden in both the training and test sets at an FDR < 0·1 as significant.

Within each validated *Mtb*-associated module, we identified hub genes by applying three criteria:

1. The gene must be protein coding.
2. It must exhibit high module membership—defined as the correlation between the gene’s expression and the module eigengene (i.e., top or bottom 2·5% of the module membership distribution, as allowing for negative correlation).
3. It must show an extremely strong association with *Mtb* burden—defined by the slope from a linear regression model (i.e., top or bottom 2·5% of the distribution of gene-*Mtb* burden association, as allowing for negative slope).

Of note, module membership defined as correlation between individual gene and the module’s eigengene (PC1) was quantified by computing the beta coefficient from a linear regression model where the module’s eigengene was regressed on the gene expression levels. This is mathematically equivalent to gene connectivity,^41^ we used these two concepts as exchangeable. We pooled data from the training and test sets to enhance statistical power for hub gene identification and subsequently compared the distribution of these hub genes across cohorts (healthy control, PTB RNA-seq, TBM RNA-seq HIV-negative, and TBM RNA-seq HIV-positive). Finally, we validated their associations with *Mtb* burden in independent PTB (n = 171) and TBM cohorts (n=281).

#### Module pathway annotation

We hypothesize that co-expressed genes within each module are functionally related and likely involved in shared biological processes. To characterize these functions, we first identified the biological processes from three data bases (GO-BP,^42^ KEGG,^43^ and Reactome^44^) in which the genes within each module participate—specifically, the pathways that contain genes from the module. We then used single-sample gene set enrichment analysis (ssGSEA) to calculate pathway activity scores for each sample across this same set of pathways. Next, we computed the correlation between these pathway activity scores and the module eigengene (PC1), which represents the module’s overall expression profile. Finally, we highlighted the module’s most relevant biological processes by selecting the top 30 pathways showing the strongest correlation with the module eigengene, aiming for a balance between interpretability and comprehensive coverage (**Figure 1A**). This ssGSEA-based correlation approach provides evidence at both the gene and patient levels for the pathways driving each module.

**Figure 1.**
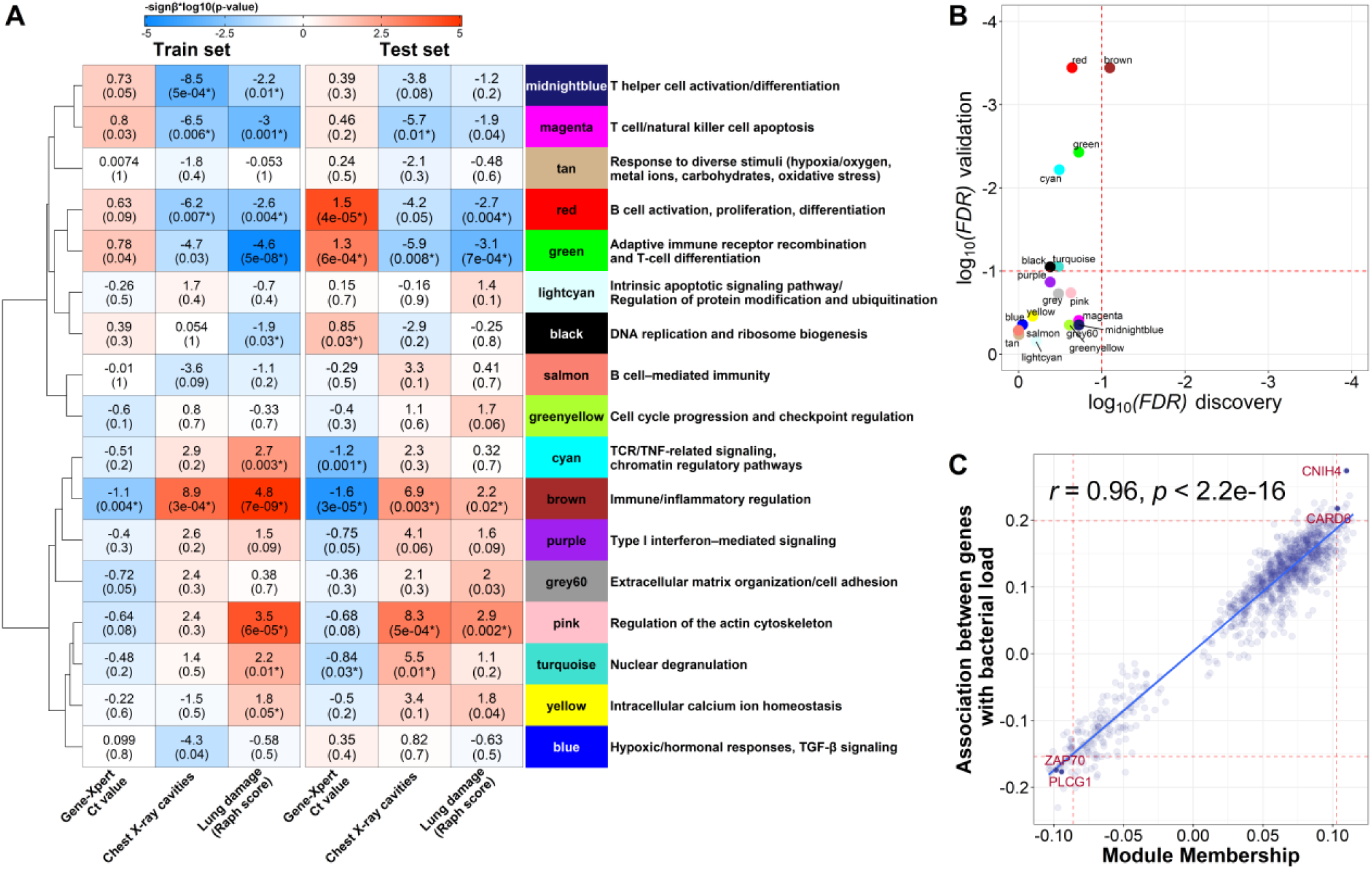
**Whole blood transcriptional modules associated with Mtb burden and lung pathology in PTB before treatment** (A) Heatmap showing the associations between WGCNA modules and Mtb burden (GeneXpert Ct value), cavities (chest X-ray), and lung damage (Raph score) in the training and test sets. Each cell displays the slope estimate (with unadjusted p-value, * indicates FDR < 0·1) derived from linear regression (for bacterial burden), logistic regression (for chest X-ray cavity), or beta regression (for lung proportion damage), with module’s eigengene (PC1) as the covariate. Blue shading indicates negative slopes, while red shading indicates positive slopes. Modules are organized based on hierarchical clustering of module eigengene correlations, as shown by the dendrogram on the left. Each module was annotated with the most highly correlated biological pathway (from GO-BP, KEGG, or Reactome) with its eigengene (PC1). See also **Figure S3**. (B) Scatterplot validating module associations with Mtb burden in training and test sets, –log₁₀(FDR) for the training set (x-axis) vs. –log₁₀(FDR) for the test set (y-axis). FDR was corrected using the Benjamini-Hochberg procedure. ^40^ Red dashed lines indicate the FDR threshold (FDR = 0·1). Among all modules, only the brown module demonstrated significant validation in the test set. ***(C)*** Scatterplot presenting the relationship between module membership (x-axis) and genes-Mtb burden association (y-axis) of 1,093 genes in the brown module. Pearson’s correlation coefficient and p-value are indicated on the plot, demonstrating the strength and significance of this relationship. Two red dashed vertical lines indicate 2·5% and 97·5% quantiles of the module membership distribution, while two red dashed horizontal lines indicate 2·5% and 97·5% quantiles of gene-Mtb burden association distribution. Genes located beyond both ranges in the same direction are considered as hub genes of the brown module.

#### Pathway analysis

To identify biological pathways linked to the clinical phenotype of *Mtb* burden, we performed an active subnetwork-oriented pathway analysis, which was implemented in the R package *pathfindR*.^45^ This analysis included an active sub-network search and enrichment analysis to connect pathways and *Mtb*-burden associated modules. First, we conducted an active subnetwork search over a curated protein–protein interaction (PPI) network from the BIOGRID database^46^, incorporating information from three databases: GO-BP; KEGG, and Reactome. This search focused on subnetworks (i.e., relevant pathways) ranging in size from 10 to 500 genes and employed a simulated annealing algorithm^47^ with 100 simulations to identify sub-networks significantly enriched for Mtb-burden–related genes. Next, we performed enrichment analysis (specifically, over-representation analysis using the GO-BP, KEGG, and Reactome databases). Enrichment statistics were computed using the hypergeometric test, and p-values were adjusted for false discovery rate using the Benjamini–Yekutieli procedure.^48^ A total of 144 pathways were enriched in the brown module with FDR < 0·05. We then assessed pathway relevance by calculating ssGSEA score for each pathway in each sample and associating these scores with the corresponding *Mtb* burden using the adjusted coefficient of determination (𝑅_adj_^2^ ) from GAM model. The pathway whose ssGSEA score showed the strongest association (highest 𝑅_adj_^2^ ) with *Mtb* burden was deemed the most relevant for further interpretation. Finally, we ranked the top 20 pathways that were most strongly associated with *Mtb* burden, grouped them into subgroups based on their similarities using hierarchical clustering analysis and pinpointed the most prominent pathways that strongly connect with the *Mtb*-associated modules (**Supplementary Figure S4B**).

### Secondary analyses

We conducted several ad-hoc and post-hoc analyses as described below

#### Deconvolution analysis

We performed a Bayesian deconvolution analysis to infer both the cellular composition and cell-type–specific expression profiles from bulk whole-blood RNA-seq, using single-cell RNA-seq (scRNA-seq) data served from healthy control as our reference priors, using the BayesPrism algorithm^49^ and its optimized re-implementation, InstaPrism,^50^ for faster computation.

We pooled the reference single-cell RNA-seq dataset from two sources: (i) human PBMCs^51^ accessed through the Azimuth web portal, comprising cells from eight HIV-vaccine trial participants sampled at days zero, three, and seven post-vaccination; and (ii) neutrophil profiles from four healthy neutrophil samples.^52^ To enhance robustness against cross-protocol gene expression differences, we applied a standard cell filtering scheme to remove low-quality cells.^53^ Data were normalized using the *sctransform* function from R package seurat. Clustering of the combined data identified 17 subtypes: neutrophil, CD14+ monocyte, CD16+ monocyte, CD4+ T memory (CD4 TM), CD4+ naive, Cytotoxic CD4+ T cell (CD4 CTL), regulatory T cell (Treg), CD8+ T memory (CD8 TM), CD8+ Naive, other T cell (otherT), B naive, B memory, plasmablast, conventional dendritic cell (cDC), plasmacytoid dendritic cell (pDC), natural killer (NK) cell, and MAIT cell (**Supplementary Figure S6A**). We then filtered for protein-coding genes, excluding mitochondrial, ribosomal, globin, sex-chromosome, and other high-noise features. Genes expressed in at least five cells were retained. From this set, 4,851 marker genes (pairwise t-test p ≤ 0·1 and log_2_FC ≥ 0·1) constituted our deconvolution signature.

Raw counts from bulk RNA-seq (healthy controls and PTB patients) were directly input into BayesPrism/InstaPrism. To minimize outlier influence, we excluded bulk genes where reads exceeded 1% of the total in >10% of samples. These cut-off values for scRNA-seq and bulk RNA-seq data were selected to ensure sufficient gene representation for each cell type and achieve optimal deconvolution performance during calibration. Bayesian inference proceeded in two phases:

4. **Initial estimation**—Gibbs sampling (chain length = 1,000; burn-in = 500; thinning = 2) with default priors (σ = 2, α = 10⁻⁸) to derive per-sample posterior cell-fraction estimates.
5. **Iterative refinement**—up to 100 iterations of conjugate-gradient updates, adjusting only the neutrophil reference to reflect its dominance in PTB (**Table 1**).

**Table 1:**
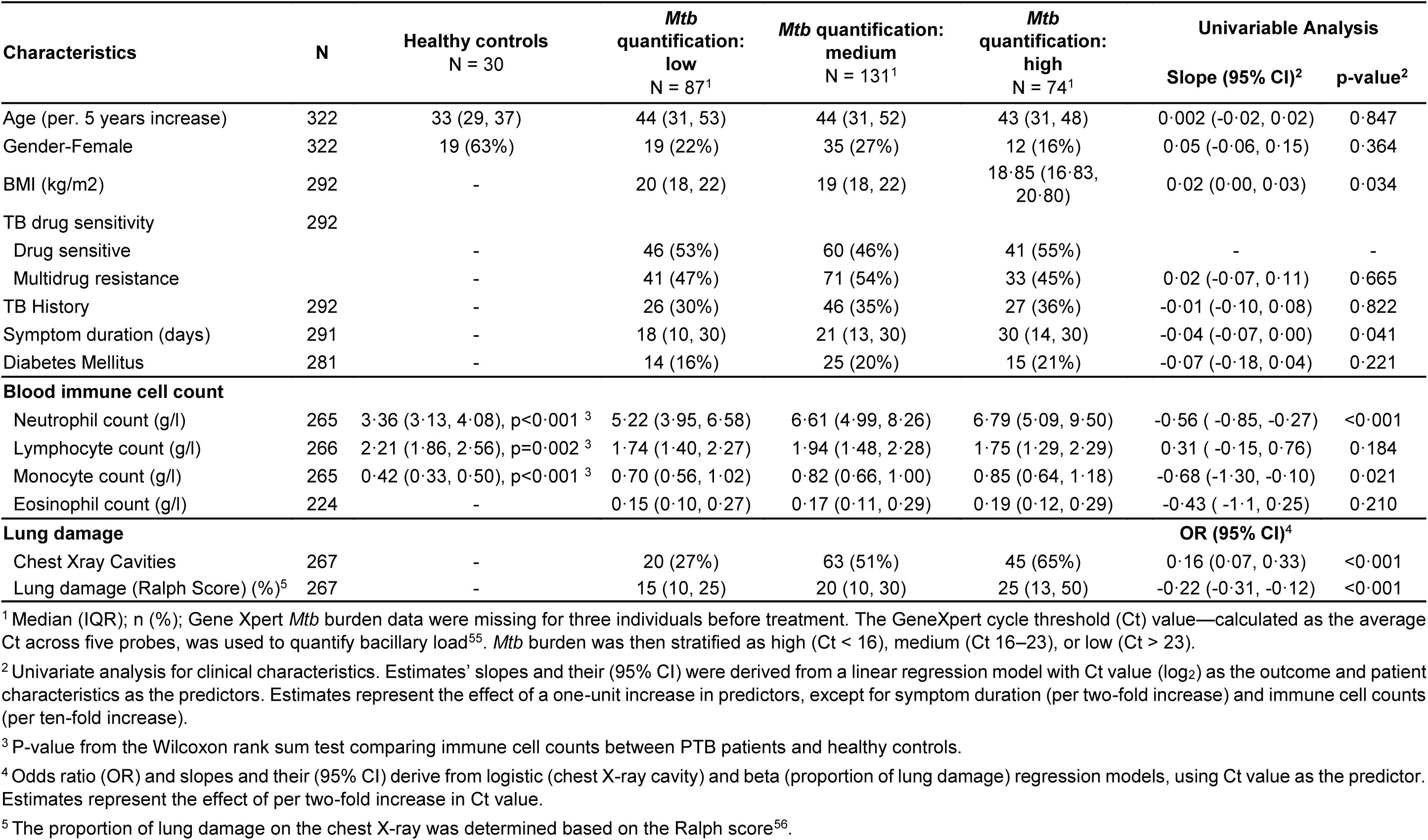
Clinical and immunological correlations with bacterial burden in PTB.

| Characteristics | N | Healthy controls<br>N = 30 | *Mtb*<br>quantification:<br>low<br>N = 87<sup>1</sup> | *Mtb* quantification:<br>medium<br>N = 131<sup>1</sup> | *Mtb*<br>quantification:<br>high<br>N = 74<sup>1</sup> | Univariable Analysis |  |
| --- | --- | --- | --- | --- | --- | --- | --- |
|  |  |  |  |  |  | Slope (95% CI)<sup>2</sup> | p-value<sup>2</sup> |
| Age (per. 5 years increase) | 322 | 33 (29, 37) | 44 (31, 53) | 44 (31, 52) | 43 (31, 48) | 0.002 (-0.02, 0.02) | 0.847 |
| Gender-Female | 322 | 19 (63%) | 19 (22%) | 35 (27%) | 12 (16%) | 0.05 (-0.06, 0.15) | 0.364 |
| BMI (kg/m<sup>2</sup>) | 292 | - | 20 (18, 22) | 19 (18, 22) | 18.85 (16.83, 20.80) | 0.02 (0.00, 0.03) | 0.034 |
| TB drug sensitivity | 292 |  |  |  |  |  |  |
| Drug sensitive |  | - | 46 (53%) | 60 (46%) | 41 (55%) | - | - |
| Multidrug resistance |  | - | 41 (47%) | 71 (54%) | 33 (45%) | 0.02 (-0.07, 0.11) | 0.665 |
| TB History | 292 | - | 26 (30%) | 46 (35%) | 27 (36%) | -0.01 (-0.10, 0.08) | 0.822 |
| Symptom duration (days) | 291 | - | 18 (10, 30) | 21 (13, 30) | 30 (14, 30) | -0.04 (-0.07, 0.00) | 0.041 |
| Diabetes Mellitus | 281 | - | 14 (16%) | 25 (20%) | 15 (21%) | -0.07 (-0.18, 0.04) | 0.221 |
| **Blood immune cell count** |  |  |  |  |  |  |  |
| Neutrophil count (g/l) | 265 | 3.36 (3.13, 4.08), p<0.001<sup>3</sup> | 5.22 (3.95, 6.58) | 6.61 (4.99, 8.26) | 6.79 (5.09, 9.50) | -0.56 (-0.85, -0.27) | <0.001 |
| Lymphocyte count (g/l) | 266 | 2.21 (1.86, 2.56), p=0.002<sup>3</sup> | 1.74 (1.40, 2.27) | 1.94 (1.48, 2.28) | 1.75 (1.29, 2.29) | 0.31 (-0.15, 0.76) | 0.184 |
| Monocyte count (g/l) | 265 | 0.42 (0.33, 0.50), p<0.001<sup>3</sup> | 0.70 (0.56, 1.02) | 0.82 (0.66, 1.00) | 0.85 (0.64, 1.18) | -0.68 (-1.30, -0.10) | 0.021 |
| Eosinophil count (g/l) | 224 | - | 0.15 (0.10, 0.27) | 0.17 (0.11, 0.29) | 0.19 (0.12, 0.29) | -0.43 (-1.1, 0.25) | 0.210 |
| **Lung damage** |  |  |  |  |  | **OR (95% CI)<sup>4</sup>** |  |
| Chest Xray Cavities | 267 | - | 20 (27%) | 63 (51%) | 45 (65%) | 0.16 (0.07, 0.33) | <0.001 |
| Lung damage (Ralph Score) (%)<sup>5</sup> | 267 | - | 15 (10, 25) | 20 (10, 30) | 25 (13, 50) | -0.22 (-0.31, -0.12) | <0.001 |

<sup>1</sup> Median (IQR); n (%); Gene Xpert *Mtb* burden data were missing for three individuals before treatment. The GeneXpert cycle threshold (Ct) value—calculated as the average Ct across five probes, was used to quantify bacillary load<sup>55</sup>. *Mtb* burden was then stratified as high (Ct < 16), medium (Ct 16–23), or low (Ct > 23).
<sup>2</sup> Univariate analysis for clinical characteristics. Estimates' slopes and their (95% CI) were derived from a linear regression model with Ct value (log<sub>2</sub>) as the outcome and patient characteristics as the predictors. Estimates represent the effect of a one-unit increase in predictors, except for symptom duration (per two-fold increase) and immune cell counts (per ten-fold increase).
<sup>3</sup> P-value from the Wilcoxon rank sum test comparing immune cell counts between PTB patients and healthy controls.
<sup>4</sup> Odds ratio (OR) and slopes and their (95% CI) derive from logistic (chest X-ray cavity) and beta (proportion of lung damage) regression models, using Ct value as the predictor. Estimates represent the effect of per two-fold increase in Ct value.
<sup>5</sup> The proportion of lung damage on the chest X-ray was determined based on the Ralph score<sup>56</sup>.

Upon convergence, the 17 subtypes were aggregated into ten major populations for downstream analyses: Neutrophil; CD14+ Monocyte; CD16+ Monocyte; CD4+ T cell (CD4+ TM, CD4+ Naive, CD4+ CTL, Treg); CD8+ T cell (CD8+ TM, CD8+ Naive); other T cell; B cell (B Naive, B memory, Plasmablast); DC (pDC, cDC); NK; and MAIT. Derived cell-type expression profiles were variance-stabilized (**DESeq2 VST**) and used for single-sample GSEA.

To validate proportions, we compared “ground-truth” myeloid (neutrophils and monocytes) and lymphocyte (all lymphoid subsets) percentages—derived from the automated blood counter—with our deconvolution estimates. We observed low root mean square errors (RMSE) and high Pearson correlations (**Supplementary Figure S6B**), confirming robust performance.

#### Variable selection analysis

Our variable selection process involved two main steps: a resampling ranking procedure and a stability variable selection analysis.

***1. Resampling Ranking Procedure:*** We began by focusing on the 10,000 most variable genes to narrow down the candidates associated with pre-treatment *Mtb* burden in sputum while accounting for sampling variability. First, we modeled the association between each gene and *Mtb* burden. Although fewer than 4% of patients had undetected bacterial levels (Ct = 40), the log₂ Ct value distribution showed a slight inflation in the tail. Given the small number of these cases, we approximated the data with a normal distribution and modeled it using linear regression, while also allowing for potential non-linear relationships by fitting a generalized additive model (GAM) with the R package *mgcv*. We used this approximation to reduce computational complexity during the subsequent resampling procedure and quantified the strength of these associations using the adjusted coefficient of determination (𝑅_adj_^2^ ).

Next, we performed 2,000 iterations of random subsampling, drawing a 2/3 subset from the test set in each iteration. In each resampled dataset, we ranked all genes by their 𝑅_adj_^2^ , generating an empirical rank distribution for each gene. We then defined “top-ranking genes” as those that consistently met the following criteria in both the training (n = 147) and test sets (n = 148):

6. A significant association with Mtb burden (p < 0·05 and 𝑅^2^ > 0 · 04).
7. Rank consistency, with the 2·5% quantile of the rank distribution below 𝑅1 = 50 and the 97·5% quantile below 𝑅2 = 500.
2. ***Stability variable selection analysis:*** We combined the top-ranking validated genes with the hub genes identified from our primary WGCNA analysis. This combined set served as input for a stability variable selection analysis aimed at pinpointing the most predictive genes for *Mtb* burden across the entire PTB discovery cohort. We implemented a gradient boosting–linear regression approach within the stability selection framework proposed by Meinshausen and Bühlmann^54^ (as implemented in the R package *stabs*). This method controls the Per-Family Error Rate (PFER) by limiting the expected number of falsely selected variables E(V) according to the following formula: 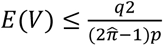 where,
8. 𝑉 is the number of variables wrongly selected by the procedure and 𝐸(𝑉) is the expected value of variables wrongly selected by the procedure
9. 𝑞 is the number of pre-anticipated selected variables
10. 𝜋^ is the selected probability
11. 𝑝 is the number of variables.

In our analysis, we pre-specified 𝐸(𝑉) = 1 (𝑃𝐹𝐸𝑅), 𝑞 = 3, and 𝑝 = 24.

## Code availability

All original code has been deposited at Zenodo at http://doi.org/10.5281/zenodo.17519084 and is publicly available as of the date of publication.

## RESULTS

### Higher bacterial burden is associated with elevated blood neutrophil and monocyte counts, and increased lung damage in PTB before treatment

We studied 295 HIV-negative people with culture-confirmed PTB. We also included 30 healthy controls, who did not have TB symptoms or recent TB exposure within the past two years, as a reference group. To identify the potential factors influencing pre-treatment *Mtb* burden, we examined associations with demographic, clinical, and immunological variables in PTB (**Table 1**).

No significant correlations were observed between *Mtb* burden and demographic variables (age, gender), prior TB history, or TB drug sensitivity (**Table 1**). However, *Mtb* burden showed a significant negative association with body mass index (BMI, *p*=0·034) and positive associations with prolonged symptom duration (*p*=0·041). Higher *Mtb* burden also correlated with increased peripheral blood neutrophil and monocyte counts (*p*<0·001 and *p*=0·021, respectively). Both cell counts were significantly higher compared to healthy controls (*p*<0·001).

As anticipated, higher *Mtb* burden was associated with chest X-ray cavities (*p*<0·001) and more extensive lung damage (*p*<0·001). These findings suggest that delayed TB diagnosis and treatment contributed to greater disease severity, with higher bacterial burden, elevated immune activation (particularly neutrophil and monocyte recruitment), and more extensive lung tissue damage. To further investigate the molecular basis of these clinical observations, we next used RNA-seq to dissect transcriptomics profiles.

### Higher bacterial burden is associated with increased innate immune response and decreased adaptive immune response

To investigate transcriptional profiles linked to *Mtb* burden before treatment, we performed whole blood RNA-seq from 295 individuals with PTB. Genes are working together in a transcriptional network; therefore, we used WGCNA to identify transcriptional modules. We randomly split the cohort into a training set (n=147) and a test set (n=148). A gene co-expression network was constructed in the training set (**Figures S1A–D**) and validated in the test set using preservation analysis (**Figures S2A-B**).

To assess the association between transcriptional responses and bacterial burden, together with other PTB severity phenotypes, constructed gene modules were tested with sputum *Mtb* burden (GeneXpert Ct value), lung cavities (chest X-ray), and lung damage (Raph score). Among 12 preserved modules, module-phenotype association analysis highlighted the **brown module**, consisting of 1,093 genes, as being robustly associated with all three severity phenotypes (**Figure 1A-B**). The **green** and **red modules** were also associated with severity, but exhibited opposite trends with weaker associations in the discovery cohort (*p*=0·04 and *p*=0·09, respectively, for *Mtb* burden, **Figure 1A-B)**.

These associations remained significant after adjusting for covariate symptom duration, a proxy for disease progression (**Figure S3**). This suggests that the observed associations reflect intrinsic immune mechanisms linked to bacterial load, rather than just the stage of the disease. Notably, among all preserved modules, the brown module showed the strongest correlation between genes’ module membership (*i.e.*, connectivity between genes within the module) and their association with *Mtb* burden (*r*=0·96, p<0·001; **Figure 1C, Table S1)**.

To identify immune regulatory pathways in the brown module, we used sub-network analysis and found multiple pathways were significantly enriched in both innate and adaptive immunity (**Figure S4A**). We calculated an activity score for each pathway in individuals using single-sample gene set enrichment analysis (ssGSEA). These pathways were then ranked by their correlation with *Mtb* burden. The top 20 pathways were grouped into three distinct sub-clusters using hierarchical clustering analysis: TNF with its related pathways, other receptor pathways, and T and B cell signaling pathways (**Figure S4B**). The five prominent pathways correlated with the overall activity of the brown module, represented by its first principal component (PC1) or eigengene, were illustrated in **Figure 2A**. Three **innate immune pathways**, including Nod-like receptor (NLR), Toll-like receptor (TLR), and Tumor necrosis factor (TNF) signaling, exhibited significant positive correlations with the brown module’s eigengene (Pearson correlation for NLR: *r*=0·69, *p*<0·001; TLR: *r*=0·67, *p*<0·001; TNF: *r*=0·71, *p*<0·001; **Figure 2A**). In contrast, two **adaptive immune pathways** (T-cell receptor (TCR) and B-cell receptor (BCR) signaling pathways) demonstrated strong negative correlations with the brown module’s eigengene (*r*=-0·92, *p*<0·001 for TCR and *r*=-0·53, p<0·001 for BCR; **Figure 2A**). These five pathways were also strongly associated with lung pathology and immune cell counts (**Figure 2B**).

**Figure 2.**
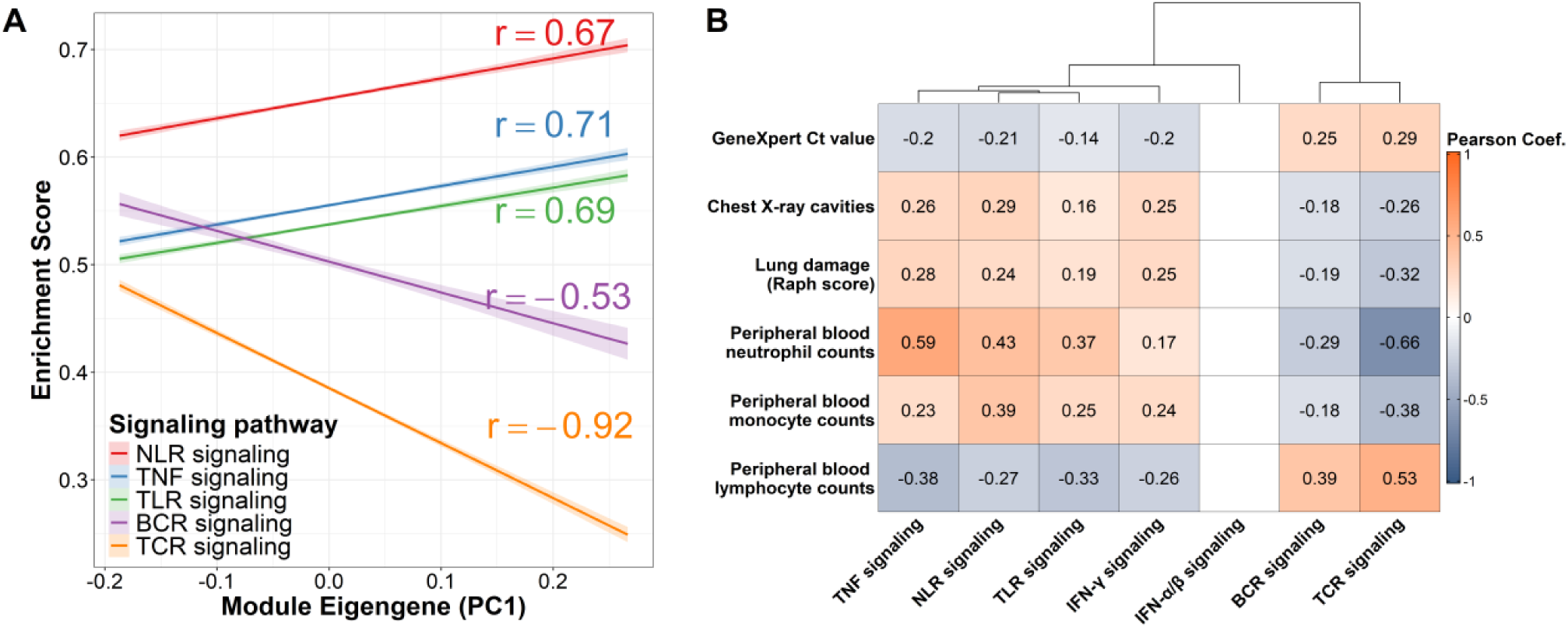
**Correlation between associated pathways and pre-treatment clinical characteristics** *(A)* Linear correlations between the brown module’s eigengene (PC1) and the single-sample GSEA enrichment scores for the five prominent Mtb burden-associated pathways. Each line represents the pathway’s enrichment score as a function of the brown module, with the Pearson correlation coefficient (r) indicated. Pathways, annotated using KEGG database, show innate Nod-like receptors (NLR) signaling, Toll-like receptors (TLR) signaling, and TNF signaling pathways; adaptive T cell receptor signaling (TCR) and B cell receptor signaling pathways. Shaded regions represent the 95% confidence intervals for the linear fits. *(B)* Heatmap showing the Pearson correlation coefficients between these five immune pathways, type I and II interferon signaling pathways (Reactome), and clinical characteristics, including Mtb burden (GeneXpert Ct), lung damage, and blood cell counts (lymphocytes, monocytes, neutrophils). Hierarchical clustering based on correlation with clinical characteristics highlights two major groups: innate immune pathways (TNF, NLR, TLR, IFN-γ signaling, IFN-α/β signaling) vs. adaptive immune pathways (TCR, BCR signaling). The color scale ranges from negative (blue) to positive (red) correlation; cells without significant correlations (p>0·05) are left blank.

Importantly, the IFN-γ (type II interferon) signaling pathway was one of the top 20 pathways associated with both *Mtb* burden and the brown module’s activity (**Figure S4A-B**). It was clustered with innate pathways (NLR and TLR signaling) (**Figure S4B**) and showed significant associations with lung cavities (*r*=0·25, *p<*0·001) and lung damage (*r*=0·25, *p<*0·001). Additionally, it also correlated with peripheral blood neutrophils (*r*=0·17, *p*=0·009), monocytes (*r*=0·24, *p<*0·001), and lymphocytes (*r*=-0·26, *p<*0·001) (**Figure 2B**).

The green and red modules, exhibiting weaker negative associations with bacterial burden, were involved in adaptive immune responses. Specifically, the green module was enriched for adaptive immune receptor recombination and T-cell differentiation pathway, while the red module was enriched for B-cell activation, proliferation, and differentiation pathway (**Figure 1A**).

### Interferon signaling pathway participates in modulating both innate and adaptive immunity in response to *Mtb* burden

To further dissect the immune landscape of PTB patients in relation to *Mtb* burden, we focused on IFN signaling, a hallmark of TB immunity.^18^ This included type I IFN (IFN-α/β) signaling, mediated by key genes STAT1, STAT2, which modulate positive feedback with innate immunity;^57^ and type II IFN signaling (IFN-γ), where STAT1 is known to negatively regulate TCR signaling pathway in TB.^58,59^ We also considered the activating role of AIM2-driven IL-1β^60,61^ in both type I and II-IFN signaling,^57^ and TNF signaling^62^ pathways. Using linear regression models, we investigated the correlations between IFN signaling (both IFN-α/β, IFN-γ) and the five identified pathways: NLR, TLR, TNF, TCR, and BCR signaling; as well as the corresponding key functional genes (**Figure 3A**).

**Figure 3.**
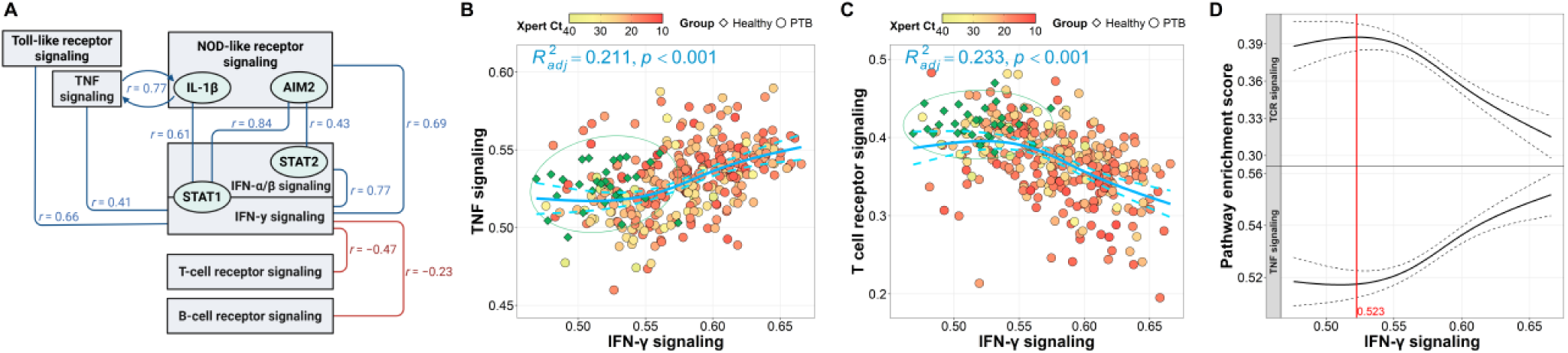
**Correlation between TNF, TCR and IFN-γ signaling pathways in PTB patients and healthy controls.** ***(A)*** Schematic diagram describing the significant Pearson’s correlation of IFN-γ signaling pathway with IFN −α/β signaling, five immune pathways, and between their key functional genes, including STAT1, STAT2, IL-1β and AIM2. **(B)** Scatterplot presenting the relationship of TNF signaling, **(C)** TCR signaling with IFN-γ signaling pathway in the study cohorts. Each point represents an individual profile, colored by Ct value for PTB patients and in green for healthy controls. Three PTB patients with missing Ct value were excluded from the plot. A smooth curve (blue line) was fitted using a generalized additive model (GAM) with a cubic regression spline, modeling the relationship of TNF, TCR and IFN-γ signaling pathways in PTB patients. The adjusted coefficient of determination 𝑅^2^ and p-value are shown for each model to indicate the percentage of each pathway activity explained by IFN-γ signaling variation and the significance of non-linear relationship. A 95% confidence ellipse, based on a bivariate t-distribution, was plotted for the healthy control in each panel. These ellipses represent the region where the data points defined by the two variables are most likely to occur. PTB patient profiles falling within this ellipse were considered comparable to those of the healthy controls. The median (IQR) Ct values for PTB patients within the ellipses were 20 (17·4, 23·9) in panel B and 21·5 (18·8, 26·3) in panel C. **(D)** Opposing trends of TNF and TCR signaling modeled by GAM as a function of increasing IFN-γ signaling. The red line indicates a critical threshold of IFN-γ signaling. See also **Supplementary Figure S5**.

Our analyses revealed a strong positive linear correlation between IFN-γ and IFN-α/β signaling (*r*=0·77, *p*<0·001). IFN-γ signaling also exhibited significant positive linear correlations with NLR, TLR (correlation coefficients: *r*=0·70, *p*<0·001, *r*=0·66, *p*<0·001, respectively; **Figure S5A-B**), and showed a notable non-linear increasing trend with TNF pathways (adjusted 𝑅²=0·21, *p*<0·001; **Figure 3B**). Across all pathways analyzed, correlations with IFN-γ signaling were consistently stronger than those with IFN-α/β signaling (**Figure S5D-H**). At the gene level, STAT1 showed a stronger correlation with AIM2, a key gene of the NLR pathway (*r*=0·84, *p*<0·001), than did STAT2 (*r*=0·43, *p*<0·001). Additionally, IL-1β expression correlated with both TNF pathway activity (*r*=0·77, *p*<0·001) and with STAT1 expression (*r*=0·61, *p*<0·001) (**Figure 3A**). Collectively, these findings reinforce the tight co-activation of innate immune pathways (NLR, TLR and TNF signaling) by IFN-γ, IFN-α/β signaling, with IL-1β and STAT1 emerging as central nodes linking these immune responses in PTB.

Conversely, IFN-γ signaling showed decreasing trends with both TCR and BCR signaling pathways. The trend with TCR signaling was a non-linear (adjusted 𝑅²=0·23, *p*<0·001, **Figure 3C**); however, for BCR signaling, the trend was linear (*r*=–0·20, *p*<0·001; **Figure S5C**). Notably, **Figure 3C** illustrates that PTB patients who exhibited TCR and IFN-γ signaling levels comparable to those of healthy controls (points within the green ellipse) had relatively lower bacterial burdens (Ct values with median (IQR) 21·5 (18·8, 26·3). However, marked suppression of TCR signaling was more likely observed in severe patients with medium and high bacterial loads (Ct values ≤ 23, red dots). This sigmoid-like pattern suggests a two-phase immune model (blue curve): (1) a **steady-state phase** under low inflammatory pressure, where TCR signaling is maintained, and (2) a **decline phase,** where TCR signaling is downregulated once IFN-γ signaling surpasses a critical threshold of 0·523 (**Figure 3D**). This threshold coincides with the onset of the upward trend in the non-linear relationship observed between IFN-γ and TNF signaling pathways (**Figure 3B, D**). Likewise, as IFN-γ signaling increases, the NLR and TLR pathways show linear increases, just like the TNF pathway, while both BCR and TCR pathways exhibit decreasing trends. These findings highlight a dynamic shift in the immune landscape of PTB, characterized by activation of innate immune signaling and suppression of TCR and BCR signaling with rising IFN-γ signaling, corresponding to *Mtb* burden, in PTB.

### PTB drives expansion of myeloid cell counts and antagonistic rewiring of innate and adaptive immune networks

Blood cell composition in TB patients could be affected by bacterial burden, which then alters transcriptional signatures within immune cell populations. To identify the relationship between pathway activities and *Mtb* burden in specific cell types, we performed a deconvolution analysis^49^ on our bulk RNA-seq data and estimated individual-level immune cell counts by leveraging unique gene expression profiles of distinct immune cell types. This analysis was applied to samples from healthy controls (n=30) and PTB patients (n=295) to compare their immune cell profiles (**Figure 4A, 4B**).

**Figure 4:**
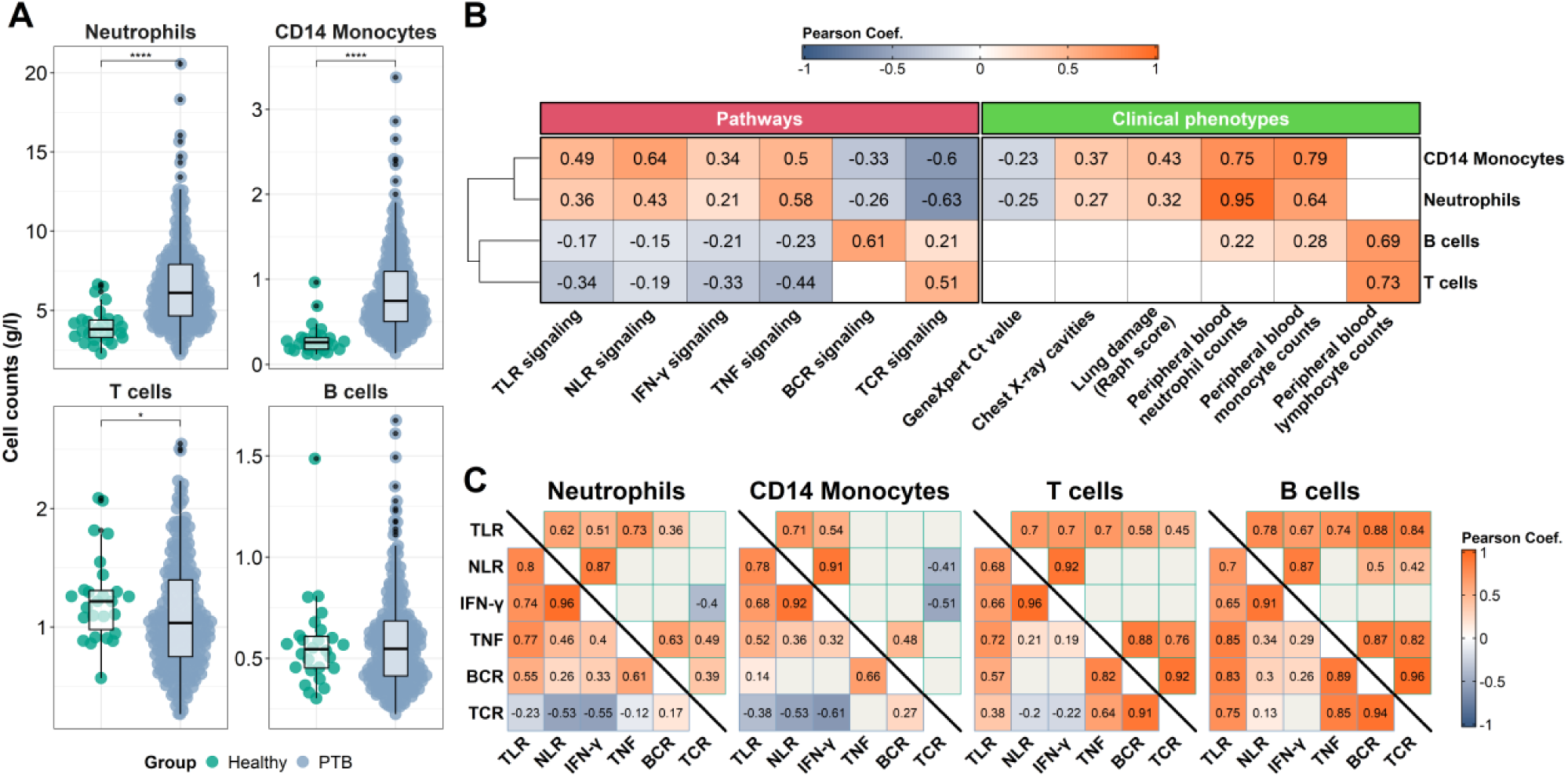
**Association of deconvoluted cell population with immune pathways and clinical phenotypes in PTB.** *(A)* Bee swarm boxplots showing the comparison of estimated cell counts (g/l) of neutrophil, CD14 monocyte, T cell and B cell between healthy control and PTB. Distributions were compared using Wilcoxon rank sum test, with significance level denoted as * (p≤0·05), ** (p≤0·01), *** (p≤0·001), **** (p≤0·0001), without correction for multiple testing. *(B)* Pearson’s correlation heatmap between estimated cell counts (g/l) of neutrophil, CD14 monocyte, T cell and B cell with immune pathways and clinical phenotypes in PTB patients; significant correlations (p<0·05) were visualised in the heatmap, while insignificant correlations were left blank. The order of cell-type was based on hierarchical clustering by computing the Pearson’s correlation coefficient between cell-types. *(C)* Pearson’s correlation between immune pathways within each cell type of PTB patients (lower triangle heatmap) and healthy controls (upper triangle heatmap). Significant correlations (p<0·05) were visualised in the heatmap while insignificant correlations were left blank with grey color.

Deconvolution using a reference single-cell RNA-seq data and applied on bulk RNA-seq to estimate the cell type composition of whole-blood samples in individuals. The method was well-calibrated, demonstrated by close alignment between measured and model-predicted values (Pearson’s correlation *r*=0·91, **Figure S6B**). Hierarchical clustering of immune cell populations in PTB revealed two major groupings: (1) myeloid subsets (neutrophils, monocytes, dendritic cells) and (2) lymphocyte subsets (T cells, B cells, natural killer cells) (**Figure S7A**). PTB patients exhibited a marked expansion of **neutrophil and CD14⁺ monocyte counts**, while healthy controls had higher proportions of T-cell subsets, particularly CD8 naïve and MAIT cells (**Figure 4A, Figure S7B, Table 1**). This expansion of myeloid cells was particularly pronounced in patients with a higher *Mtb* burden and more extensive lung damage, highlighting its link to disease severity (**Figure 4B**, **Table 1**). Furthermore, by correlating cell counts with pathway activity (**Figure 4B**), we observed that neutrophil and CD14⁺ monocyte counts were strongly positively associated with innate inflammatory signaling (TLR, NLR, TNF, and IFN-γ pathways), and inversely correlated with adaptive immune signaling (TCR and BCR pathways). Whereas lymphocyte (T and B cell) counts showed the opposite relationship: they were positively correlated with TCR and BCR pathway activity and inversely associated with innate immune pathways. Notably, lymphocyte counts showed no significant association with bacterial burden or lung pathology.

To dissect pathway interplay within each cell type, we examined network correlations among the associated immune signaling pathways (**Figure 4C**). In PTB, neutrophils and CD14⁺ monocytes exhibited very strong co-activation among TLR, NLR, and IFN-γ pathways (Pearson’s correlation *r*=0·7–0·96), and moderate correlations with TNF signaling. The TNF’s correlations were substantially weaker or absent in healthy controls, indicating that TB disease amplifies and coordinates inflammation responses within myeloid cells. In both PTB and healthy controls, T and B lymphocyte subsets showed robust TCR–BCR pathway correlations (*r*=0·91–0·94) and also extended to TNF signaling (*r*=0·64–0·89), indicating strong coordination of these pathways in lymphocytes. Notably, only in PTB, we observe significantly negative correlations between TCR signaling and the cluster of innate response pathways (NLR, TLR, and IFN-γ) in myeloid cells, a pattern not seen in healthy controls. This suggests an antagonistic rewiring between innate and adaptive immune networks during TB.

Together, these data demonstrate that PTB is marked by profound shifts in blood immune cell composition and a fundamental rewiring of immune signaling networks, characterised by elevated innate activation and antagonism between innate and adaptive responses.

### CNIH4, the strongest hub gene of brown module, is associated with bacterial burden and T-cell receptor signaling pathway

To determine specific genes associated with *Mtb* burden, we focused on hub genes, which are most strongly connected to other genes in the module, characterized by high module membership within the brown module and strong correlation with *Mtb* burden (**Figure 1C**). We identified 20 such hub genes, including key regulators of TCR signaling (ZAP70, PLC-γ1), NLR signaling (CARD6), and TNF signaling (TRAF1) (**Table S2**).

Among them, CNIH4 (cornichon family AMPA receptor auxiliary protein 4), known as the endoplasmic reticulum (ER)–Golgi cargo protein for the family of G-protein-coupled receptors (GPCR),^63^ emerged as the most prominent hub gene in the brown module. It showed the highest module membership and correlated with *Mtb* burden (**Table S2, Figure 1C**). In particular, CNIH4 accounted for 13% of the observed variation in bacterial load (non-linear association, 𝑅^2^ =0·13, *p*<0·001; **Figure 5A**). CHIH4 gene expression also demonstrated a strong negative correlation with TCR signaling activity (Pearson correlation *r*=–0·81, *p*<0·001; **Figure 5B**), suggesting elevated CNIH4 coincides with suppressed TCR pathway function.

**Figure 5:**
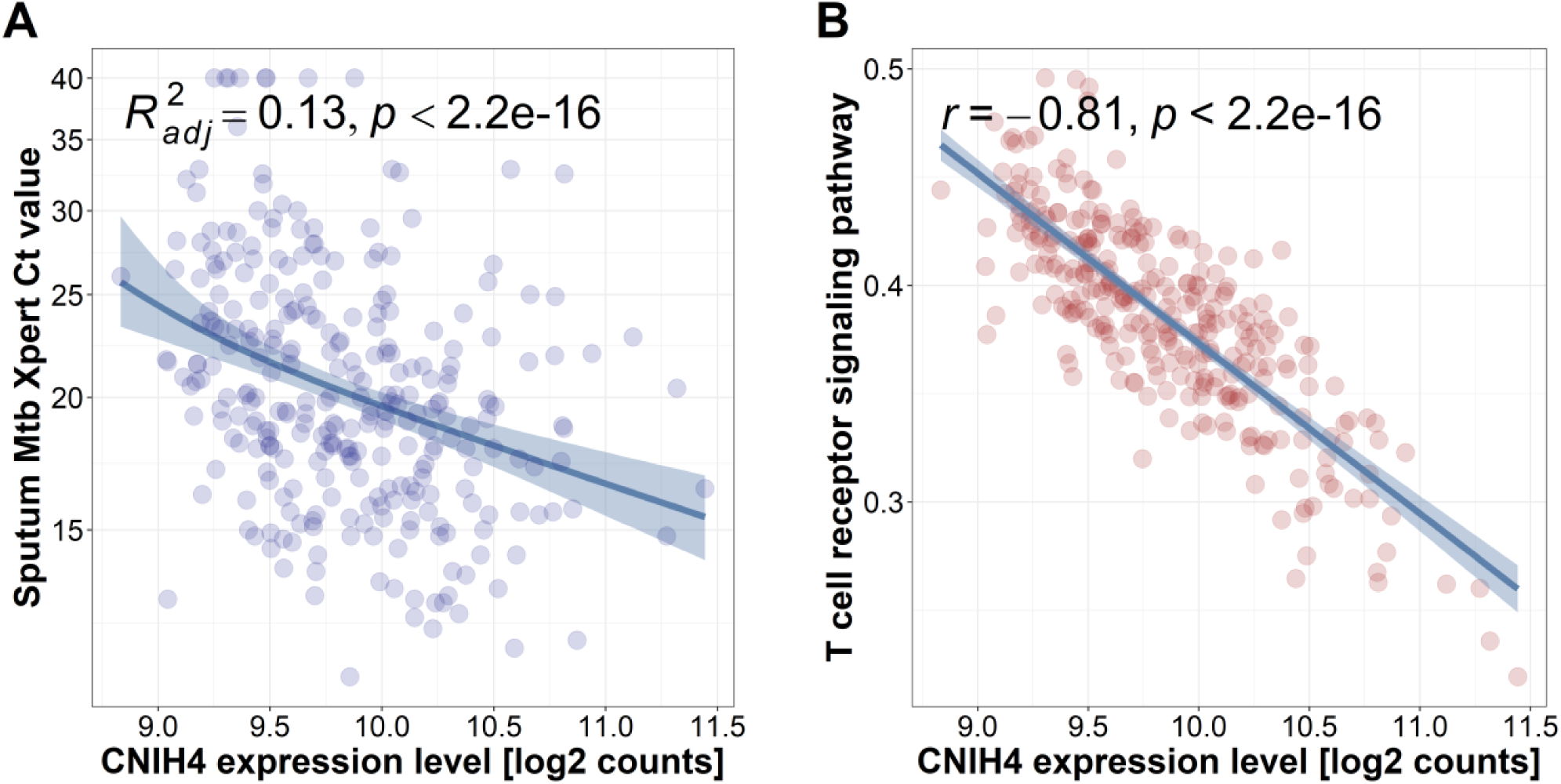
**CNIH4 gene expression was associated with Mtb burden and T-cell receptor signaling pathway.** *(A)* Scatterplot shows the association of CNIH4 expression levels with bacterial burden measured by sputum Mtb GeneXpert Ct value. Blue line was fitted line from the hurdle regression model in which Ct value was the outcome, CNIH4 expression was the covariate. The shaded band presents 95% confidence interval of the estimated trend. Adjusted coefficient of determination 𝑅^2^ and p-value are shown on the plot to indicate the percentage of Mtb variation explained by CNIH4 expression variation and significance of their non-linear relationship. *(B)* Scatterplot shows the association of CNIH4 expression level with the single-sample GSEA enrichment score for the TCR signaling pathway. Blue line is the fitted line from the linear regression model in which Ct value was the outcome, CNIH4 expression was the covariate. The shaded band presents 95% confidence interval of the estimated trend. Pearson correlation coefficient and p-value are displayed on the plot to highlight the strength and significance of the relationship.

To test whether CNIH4 was involved in suppressing TCR signaling in whole blood, we examined gene expression correlations between CNIH4 and known immune checkpoint ligands (PD-L1/2, CD80/CD86, Galectin-9, and MHC Class II), and blood immune cell counts (**Figures S8**). We found that CNIH4 strongest positively correlated with **CD274** (gene coding for PD-L1, Pearson correlation *r*=0·84, *p*<0·001). It also positively correlated with **neutrophil** (*r*=0·64, *p* < 0·001) and **monocyte** counts (*r*=0·43, *p*<0·001), but negatively correlated with **lymphocyte** counts (*r*=-0·34, *p*<0·001). Furthermore, CNIH4 also negatively correlated with ZAP70 and PLC-γ1, two other hub genes of the ***brown*** module (*r*=-0·68, *p*<0·001, *r*=-0·72, *p*<0·001, respectively).

### CNIH4 as a predictive biomarker for *Mtb* burden: validation in PTB, TBM and animal models

To identify predictive host response biomarkers for *Mtb* burden, we first performed variable selection analysis in a discovery PTB cohort. We then validated the biomarkers in independent human cohorts, including another PTB cohort (n=171) and TBM cohort (HIV-negative (n=207) and HIV-positive (n=74)^27^). Healthy controls were used as a reference group to compare the gene expression of biomarkers across different forms of TB disease. We also re-analyzed published animal data to compare the expression of these biomarkers in systemic and tissue levels.

In the discovery PTB cohort, variable selection analysis identified **CNIH4** and **DPRXP2** (a pseudogene) as top predictors of bacterial burden (**Figure 6A**). DPRXP2 (divergent-paired related homeobox pseudogene 2) gene expression was strongly correlated with CNIH4 (*r*=0·79, *p*<0·001) and significantly associated with *Mtb* burden in PTB (𝑅^2^ =0·119, *p*<0·001). In the independent PTB cohort, CNIH4 expression measured by RT-qPCR was also significantly associated with *Mtb* burden by Simes’s test 𝑝_𝑜𝑣𝑒𝑟𝑎𝑙𝑙_ = 0·014 (**Figure 6B**), indicating significance driven by CNIH4 effects on either the odds of undetectable burden (𝑝_ℎ𝑢_) or the magnitude of bacterial burden among *Mtb* detected cases (𝑝_𝑚𝑢_ ). A similar association was observed in cerebrospinal fluid (CSF) of TBM patients (𝑝_𝑜𝑣𝑒𝑟𝑎𝑙𝑙_ <0·001; **Figure 6C**). Across all cohorts, CNIH4 expression demonstrated a significant upward trend from healthy controls to PTB (HIV-negative) and TBM (HIV-negative and HIV-positive) patients (Jonckheere-Terpstra test, *p*<0·001, **Figure 6D**).

**Figure 6:**
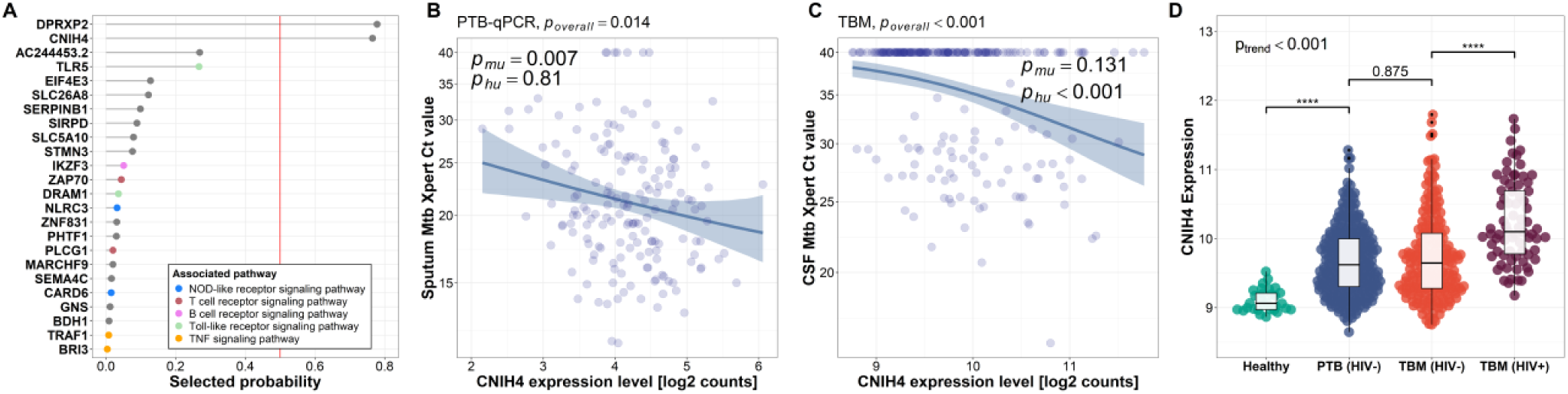
**CNIH4 as a predictive blood biomarker for Mtb burden in PTB and TBM** *(A)* Variable selection analysis identifies 24 genes most consistently predictive of Mtb burden in PTB. Each gene node is colored by their associated biological pathway. The x-axis shows the probability of selection for each gene. The red vertical line indicates the cut–off selection probability of 0·5. *(B)* Scatterplot shows a significant association of blood CNIH4 expression by RT-qPCR with Mtb burden in sputum samples from PTB patients and (C) in CSF samples from TBM patients. 51 TBM patients were removed from the plot due to missing Ct value. Blue line was fitted line from the hurdle regression model in which GeneXpert Ct value was the outcome, CNIH4 expression was the covariate. The shaded regions around the regression line indicate the 95% confidence interval for the predicted values. P-values from the non-hurdle component (𝑝_𝑚𝑢_ ) and the hurdle component ( 𝑝_ℎ𝑢_ ) are reported, along with a global p-value assessing whether at least one of the components is significant. The global test was performed using Simes’ procedure to control the family-wise error rate. ^37^ For PTB, the global p-value was 𝑝_𝑜𝑣𝑒𝑟𝑎𝑙𝑙_ = 0 · 014; for TBM 𝑝_𝑜𝑣𝑒𝑟𝑎𝑙𝑙_ < 0 · 001, to indicate the overall significance of the association. (D) Bee swarm boxplots show the distribution of CNIH4 expression in comparison among groups: healthy controls, PTB, TBM HIV- and TBM HIV+. Distributions were compared using Wilcoxon rank sum test, with significance level denoted as * (p ≤ 0·05), ** (p ≤ 0·01), *** (p ≤ 0·001), **** (p ≤ 0·0001). P-value in the top-left from Jonckheere-Terpstra trend test showing an increase of CNIH4 expression level among healthy and TB disease groups.

To investigate whether blood signatures reflect local responses, we analyzed published RNA-seq data from lung tissues of *Mtb*-infected mice (n=39) and macaques (n=16).^36^ These datasets included both infected and matched-uninfected animals. In both species, CNIH4 showed a positive but non-significant correlation with bacterial burden (mice: *r*=0·26, *p*=0·17; macaques: *r*=0·38, *p*=0·24). However, other innate immune genes, including NOD2, AIM2, TLR4, TLR6, IL-1β and STAT1, were significantly upregulated and positively correlated with bacterial load. These trends mirrored observations in PTB patients (**Figure 7A-C**). For hub genes in the TCR signaling pathway, ZAP70 showed a positive correlation with bacterial burden in both mice (*r*=0·41, *p*=0·028) and macaques (*r*=0·78, *p*=0·005). PLC-γ1 was inversely correlated in mice (*r*=–0·61, *p*<0·001) but did not exhibit a significant trend in macaques (*r*=0·47, *p*=0·14). These patterns partly diverged from those in PTB patients, where both ZAP70 and PLC-γ1 were negatively associated with bacterial burden (ZAP70: *r=-0·23, p*<0·001; PLC-γ1*: r*=-0·27, *p*<0·001) (**Figure 7C**).

**Figure 7:**
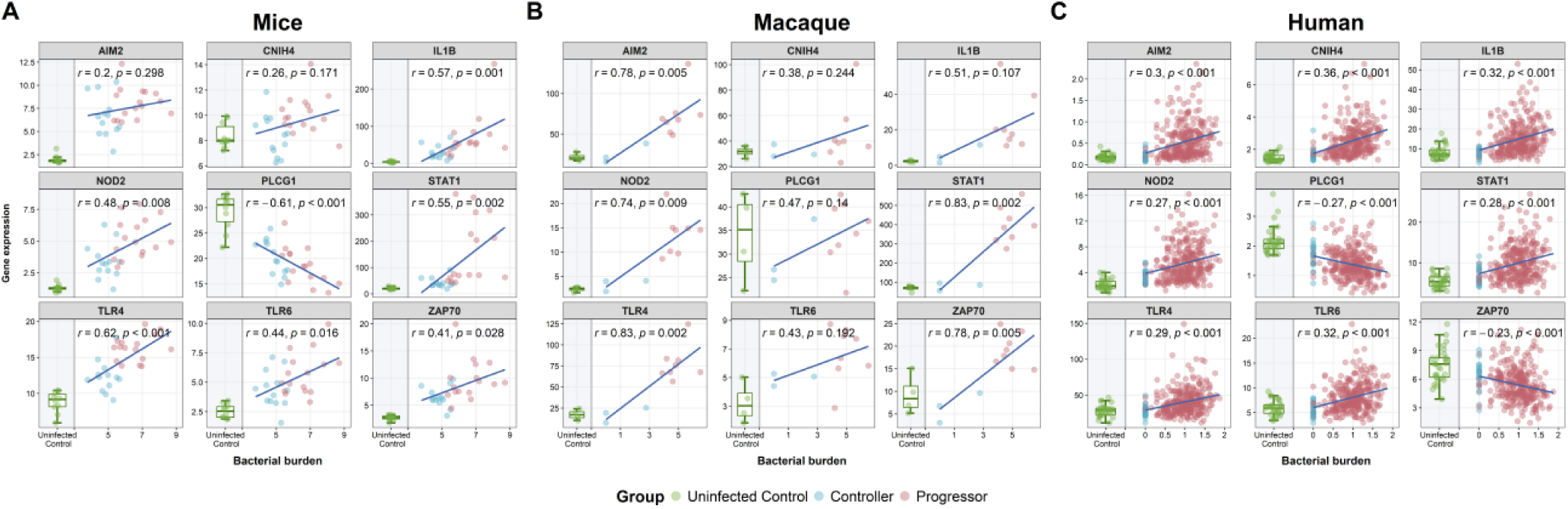
**Correlation of key immune gene expression with bacterial burden in lung tissue from Mtb-infected mice and macaques.** This figure illustrates the correlations between bacterial burden and the expression of immune-related genes (TLR4, TLR6, AIM2, NOD2, STAT1, IL-1β, PLC-γ1, ZAP70, and CNIH4) in lung tissue from Mtb-infected mice and macaques. Mice (10 naïve, 13 controllers, 16 progressors) were infected with ∼100 CFU of Mtb strain HN878, and macaques (4 naïve, 4 controllers, 8 progressors) were exposed to Mtb strain CDC1551. Lung tissue was collected and analyzed via RNA-seq. Each panel shows the gene expression levels (FPKM, y-axis) against bacterial burden (log_10_ CFU, x-axis) for individual animals, stratified by infection status (naïve, controller, progressor). Lines represent simple linear regression fits, with Pearson correlation coefficients (r) and p-values indicated. Data were re-analyzed from Ahmed et al., Science Translational Medicine, 2020. ^36^ Human samples include 30 healthy and 295 PTB individuals from this study as uninfected controls (green) and progressors (red), respectively. The controller group (blue) includes drug-susceptible TB cases with acid-fast bacilli (AFB) positive but GeneXpert negative. Bacterial burden in humans is calculated using log_2_(40/GeneXpert Ct value).

## DISCUSSION

Our study of 295 HIV-uninfected Vietnamese adults with PTB demonstrates blood transcriptional signatures associated with sputum bacterial burden. We found that higher pre-treatment sputum bacterial loads are accompanied by a coordinated shift in blood transcriptional response: innate pathways (TLR, NLR and TNF signaling) and IFN-γ signaling pathway increased, whereas adaptive receptor signaling pathways (TCR and BCR) decreased. This pattern indicates antagonistic rewiring between innate and adaptive immunity. We also observed a marked expansion of myeloid cell counts, particularly neutrophils and classical (CD14⁺) monocytes, in individuals with high bacterial burden and extensive lung damage. These myeloid populations, in interaction with TCR, are primarily drivers of this antagonistic immune rewiring. Notably, the ER–Golgi cargo receptor coding **CNIH4** emerged as the strongest hub gene linking high bacterial burden and TCR suppression, a relationship that confirmed across both PTB and TBM cohorts, and animal lung transcriptional profiles, suggesting its promise as a blood biomarker for treatment monitoring and host-directed therapy target.

Mechanistically, our data reinforce the established roles for innate sensors in *Mtb* containment and inflammatory pathology. TLR and NLR family members detect pathogen- and damage-associated patterns to trigger TNF-α and IL-1β production (Box 1). While this response is critical for initial bacterial control, excessive innate activation results in lung tissue damage. In high-burden patients, we observed the tight co-expression of innate immune pathways (TLR, NLR and TNF) coupled with inverse correlations with TCR/BCR signaling. The IFN–γ signaling pathway is involved in modulating these two opposing responses, causing a negative feedback loop. Our cell type composition further confirmed that the expansion of neutrophils and classical monocytes (CD14⁺) underlies this antagonistic rewiring and suppresses adaptive immunity, consistent with persistent lymphopenia and impaired antigen-receptor signaling observed in severe TB.^64^ Collectively, these findings suggest that excessive myeloid-driven innate activation contributes to adaptive immune suppression and may underlie the unfavorable treatment outcomes observed in patients with high *Mtb* burden.^3^ Our findings highlight the need for combination therapies that incorporate novel host-directed strategies, in addition to targeting bacterial eradication and inflammatory modulation.^65^ Adjunctive therapies should aim to prevent downregulation of adaptive immune responses, specifically TCR and BCR signaling, in order to sustain or restore effective adaptive immunity, thereby optimizing bacterial control while minimizing host tissue injury.

A key element in the TB immune crosstalk is IFN signaling, which is crucial for macrophage activation and T-cell effector function.^16,18^ Our data show that IFN-γ signaling pathway co-expresses with innate inflammatory pathways and increases with disease severity. However, in patients with high *Mtb* burden, excessive IFN-γ signaling paradoxically suppresses T-cell (possibly B-cell) receptor signaling via negative regulatory feedback, potentially mediated through immune checkpoints such as programmed cell death protein 1 and its ligand (PD-1/PD-L1). This aligns with evidence from tumor immunology IFN signaling pathway regulates the PD-L1 expression via the JAK–STAT–IRF-1 pathway38,39 and TB study showing that PD-1 expression on *Mtb*-specific CD4 T cells in smear-positive patients correlates with bacterial load.^68^ Furthermore, mouse studies demonstrate that PD-1 limits lethal over-production of IFN-γ,^69^ and *in vitro* blockade of PD-1 rescues *Mtb*-specific IFN-γ-producing T cells from apoptosis.^59^ From these findings, we proposed a two-phase IFN-γ-dependent immune model: **phase 1** involves steady-state TCR function at low bacterial loads with minimal IFN-γ signaling, and **phase 2** involves declining TCR signaling via PD-1/PD-L1 axis under intense inflammation as IFN-γ signaling response surpasses a critical threshold. This model aligns with a recent single-cell RNA-seq study of PTB lung tissues,^70^ which demonstrated that immunosuppressive myeloid subsets and Tregs upregulate genes linked to disease severity. Together, our work underscores the delicate balance between protective innate inflammation and the suppression of adaptive immunity in TB pathophysiology.

Within this network, CNIH4 gene stands out as both a mechanistic node for TCR signaling pathway and a promising blood-based biomarker for *Mtb* burden. It shows strong negative correlations with ZAP70 and PLC-γ1 hub genes. These genes are downstream targets of the receptor PD-1 in the TCR signaling pathway.^71,72^ These findings suggest that CNIH4 may contribute to TCR signaling downregulation via the PD-1/PD-L1 axis. As a COPII cargo receptor, CNIH4 regulates ER-to-Golgi trafficking of GPCRs.^63^ We speculate that its expression may contribute to the export of PD-L1, thereby amplifying PD-1 mediated suppression of TCR signaling. Across PTB, TBM, and animal models, CNIH4 shows robust associations with increased bacterial burden. Moreover, in smear-positive patients, PD-1 expression on *Mtb*-specific CD4⁺ T cells declines following anti-TB treatment.^68^ Together, findings from correlation analysis and biological connection support that CNIH4, a gene involved in TB pathophysiology, could be used as a blood biomarker for monitoring bacterial burden and as a potential host-directed therapeutic target to restore T-cell responses in PTB.

A strength of our study is the large, well-characterized clinical cohorts and the integrated whole-blood transcriptomic approach and advanced statistical model analyses. Crucially, we leverage the broad spectrum of pre-treatment *Mtb* burden, an inherent feature of large clinical cohorts but typically absent in controlled experimental models. This allows us to uncover complex, non-linear interactions between innate and adaptive immune pathways, cell types, and the potential biomarker CNIH4 in relation to pre-treatment *Mtb* burden and lung pathology. However, the cross-sectional design limits causal inferences and understanding immune dynamics after initiating the anti-TB treatment. Likewise, the systemic transcriptional profile could not fully capture the lung transcriptional profiles of these PTB patients.

**In conclusion**, our findings demonstrate that high pre-treatment *Mtb* burden was associated with a dysregulated immune response, driven by neutrophils and monocytes, characterized by excessive innate and IFN-γ signaling, and impaired adaptive immunity. The CNIH4–TCR axis is involved in this shift, which provides insights into TB pathophysiology. CNIH4 is a promising predictive biomarker for treatment monitoring and a candidate host-directed therapeutic node for restoring adaptive responses in severe disease.

### APPENDIX

**Box 1:**
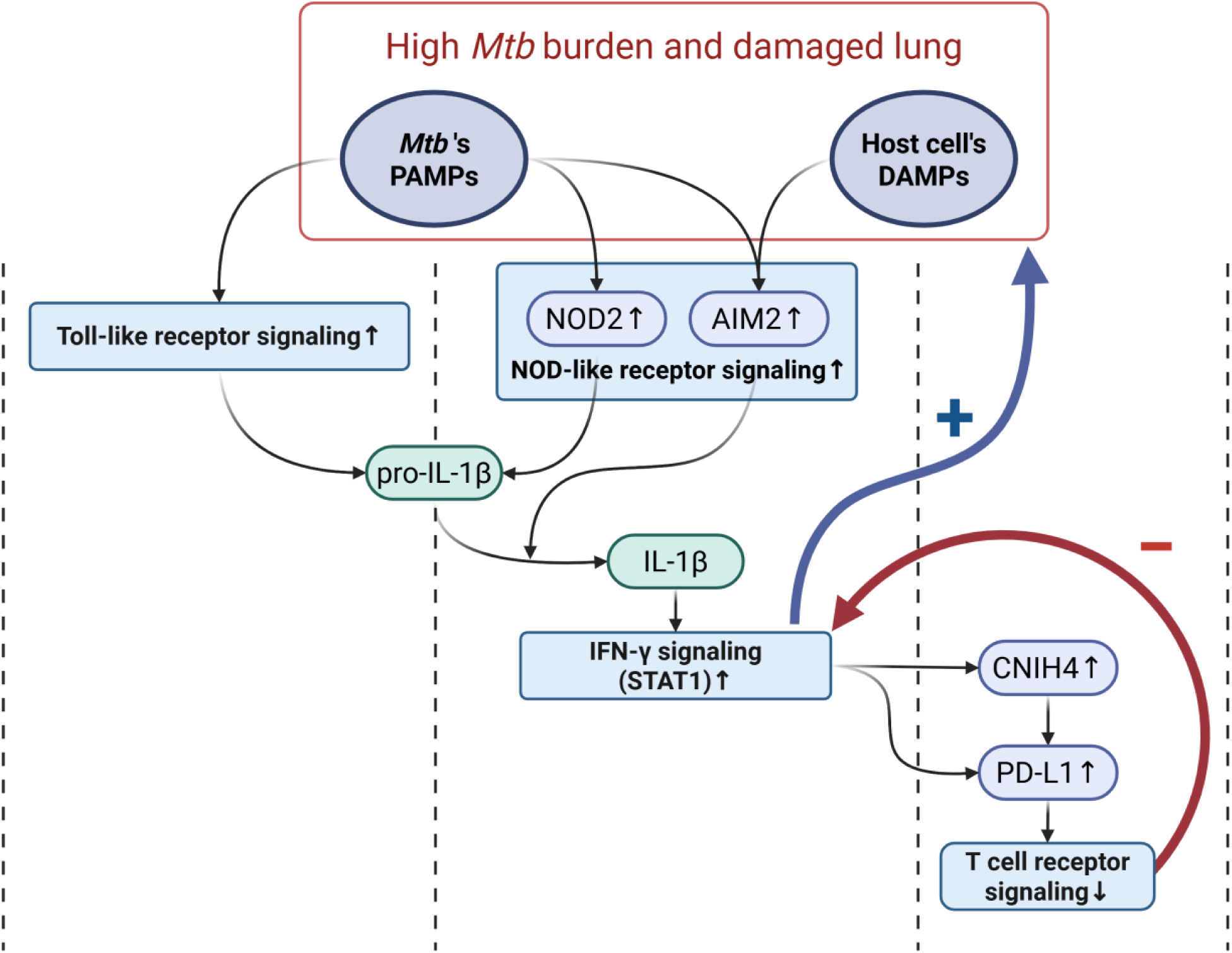
Toll-like receptors and Nod-like receptors signaling pathways in TB immunity. Toll-like receptors (TLRs), particularly TLR2, TLR4, and TLR6, located on cell membranes, recognize Mtb PAMPs and activate MyD88-dependent pathways, leading to the activation of transcription factors NF-κB and AP-1. These pathways upregulate pro-inflammatory cytokines (IL-1β, TNF-α, IL-6), promoting the recruitment of immune cells to the infection site and aiding in controlling Mtb^5^. In addition, TLR activation via MyD88-independent mechanisms induces type I interferon responses, including IFN-α and IFN-inducible genes, further upregulating the immune response against Mtb.^4,5^ NOD-like receptors (NLRs), a family of cytoplasmic sensors, such as NOD2 and the absent in melanoma 2 (AIM2), complement TLR-mediated responses by detecting intracellular signals. NOD2, for example, recognizes muramyl dipeptide (MDP) from the Mtb cell wall, also activating NF-κB and contributing to the production of pro-IL-1β^73–75^ Additionally, the AIM2 inflammasome is triggered by sensing Mtb-derived double-stranded DNA (dsDNA) within phagocytosed Mtb or damaged host cells. This activation cleaves pro-caspase-1 into its active form, caspase-1, which processes pro-IL-1β into mature IL-1β, further amplifying IL-1β production ^76–78^. Notably, TNF-α and IL-1β engage in a positive feedback loop: IL-1β upregulates TNFR1 and TNF-α secretion, and TNF-α, particularly TNF signaling pathway in turn promotes IL-1β production, immune cell activation, oxidative stress (iNOS), and programmed cell death—all aimed at containing Mtb.^79^ TNF-α and IL-1β-mediated inflammation is essential for controlling bacterial growth, excessive production can lead to tissue damage.^8,9^

**Interferons signaling pathways in TB immunity:** Type I IFNs (IFN-α/β), induced by virulent *Mtb* strains via ESX-1, suppress Th1 and IFN-γ responses, enhance IL-10 production, and promote apoptosis, undermining bacterial control.^57,58^ Type II IFN (IFN-γ), produced by T cells and innate lymphoid cells, activates macrophages, drives granuloma formation, and enhances reactive species production, essential for resistance to *Mtb*.^58^ In vitro PD-1 blocking rescued *Mtb*–specific IFN-γ–producing T cells from undergoing apoptosis. The number of PD-1–expressing T cells decreased significantly during therapy and inversely correlated with IFN-γ–dominant T-cell response against *Mtb*. ^59^

## ACKNOWLEDGEMENTS

The authors thank the study participants, study coordinators, doctors, nurses and research support staff at the Oxford University Clinical Research Unit, Pham Ngoc Thach Hospital and Hospital for Tropical Diseases. We also thank Professor Thomas R. Hawn for giving insightful review and suggestions on the manuscript.

## AUTHOR CONTRIBUTIONS

Conceptualization, L.T.H.N., L.H.V., G.T. and N.T.T.T.; Data Generation, H.T.H., L.H.V., L.N.H.T., T.T.B.T., D.D.A.T., D.T.M.H., H.D.T.N.; Methodology, L.T.H.N., T.M.T., H.T.H., N.T.T.T.; Data Analysis: L.T.H.N., T.M.T H.T.H..; Investigation: L.T.H.N., T.M.T., H.T.H and N.T.T.T; Writing – Original Draft, L.T.H.N., T.M.T, H.T.H, G.T and N.T.T.T.; Writing – Review & Editing, all authors. Corresponding authors had unrestricted access to all data. All authors agreed to submit the manuscript, read and approved the final draft and take full responsibility of its content, including the accuracy of the data.

## FUNDING

This work was supported by the Wellcome Trust (206724/Z/17/Z to N.T.T.T and 110179/Z/15/Z to G.T), and National Institutes of Health (U19AI162583). The funders had no role in study design, data collection and analysis, decision to publish, or preparation of the manuscript.

## SUPPLEMENTAL INFORMATION

**Figure S1.**
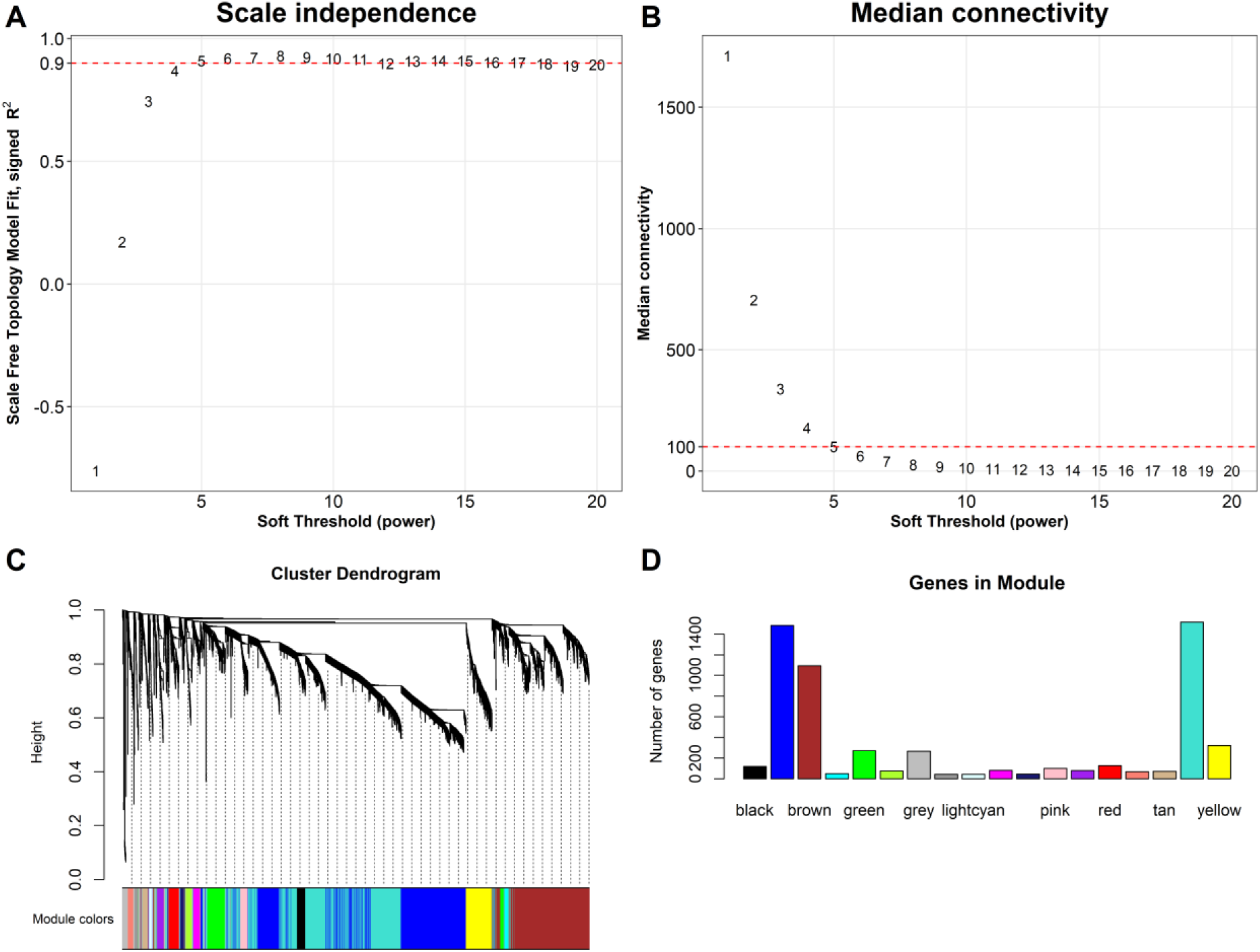
**Construction of WGCNA in train set** Analysis of network topology for various soft-thresholding powers of the top 5,850 most variant genes based on the scale-free network model. A soft-thresholding power was optimally determined to balance between enforcing scale-free topology and preserving meaningful gene-gene interactions. **(A)** The scale-free fit index (y-axis) as a function of the soft-thresholding power (x-axis). The red horizontal dash line corresponds to R^2^ = 0 · 9 and soft-thresholding power β=5, which was chosen for the construction gene-expression network. **(B)** The median connectivity on the y-axis as a function of the soft-thresholding power on the x-axis. The adjacency matrix of the scale-free network between genes was determined as A = (a_ij_ ), where a_ij_ = |cor(gene_i_, gene_j_)|^β^. Red line indicates the optimal median connectivity using soft-thresholding power β=5. **(C)** Dendrogram resulted from the hierarchical clustering analysis using topological overlap of the adjacency matrix A served as a dissimilarity metric. Each cluster was referred as a module and assigned with a color. In total, we identified 17 transcriptional modules in the training set, excluding the grey module as it consisted of non-connected genes. **(D)** Bar-plots indicate the number of genes contained in each module.

**Figure S2.**
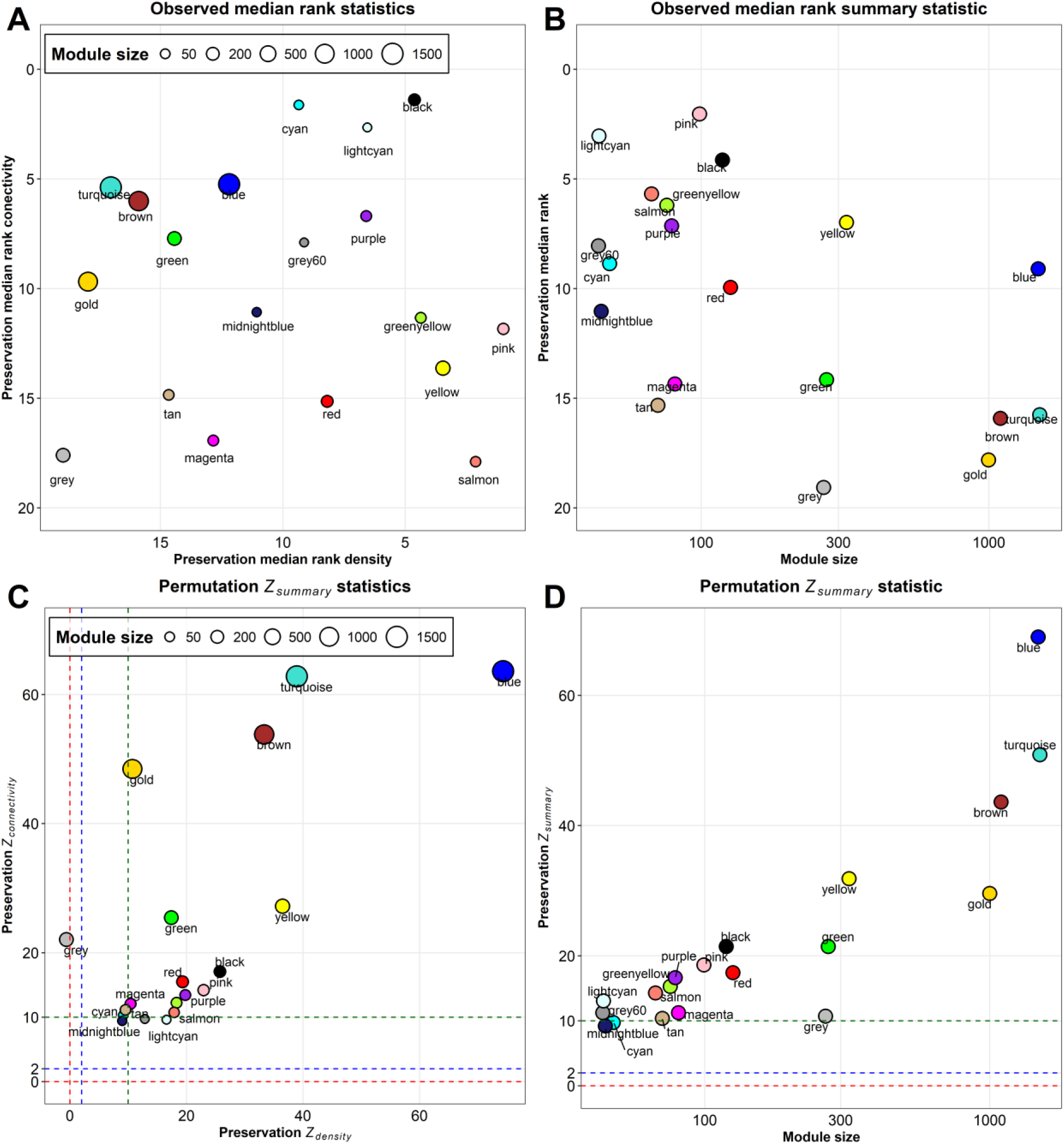
Preservation of discovery modules in validation set. **(A)** The median rank preservation statistics of the modules. Each module was represented by a point, labeled with the corresponding color and name. The X and Y-axis represent the median rank density and connectivity of observed preservation statistics per module, respectively. A low preservation median rank value indicates a high level of preservation. The gold module was an artificial module comprised of 500 randomly selected genes. The grey module consisted of non-connected genes identified in the WGCNA analysis of the discovery cohort. **(B)** The median rank summary statistic by module size showing gold and grey modules exhibited the highest median rank, indicating low preservation, which ensured our preservation analysis controlling well the background noise signal. **(C)** The Z_density_ and Z_connectivity_ preservation statistic of the modules. Each module was represented by a point, labeled with the corresponding color and name. The X-axis and Y-axis represents the Z_density_ and Z_connectivity_ statistic of each module based on 1,000 permutations of module labels, respectively. **(D)** The Z_summary_ preservation statistic by module size. In both figures, the blue and green dash lines correspond to the threshold of Z statistics of 2 and 10, where ≥10: High preservation; 2<Z<10: Moderate preservation; <2: Low preservation. Overall, using the threshold of Z≥10 across density, connectivity and summary statistics, 12 out of 17 identified modules in the training set were validated in the validation set.

**Figure S3.**
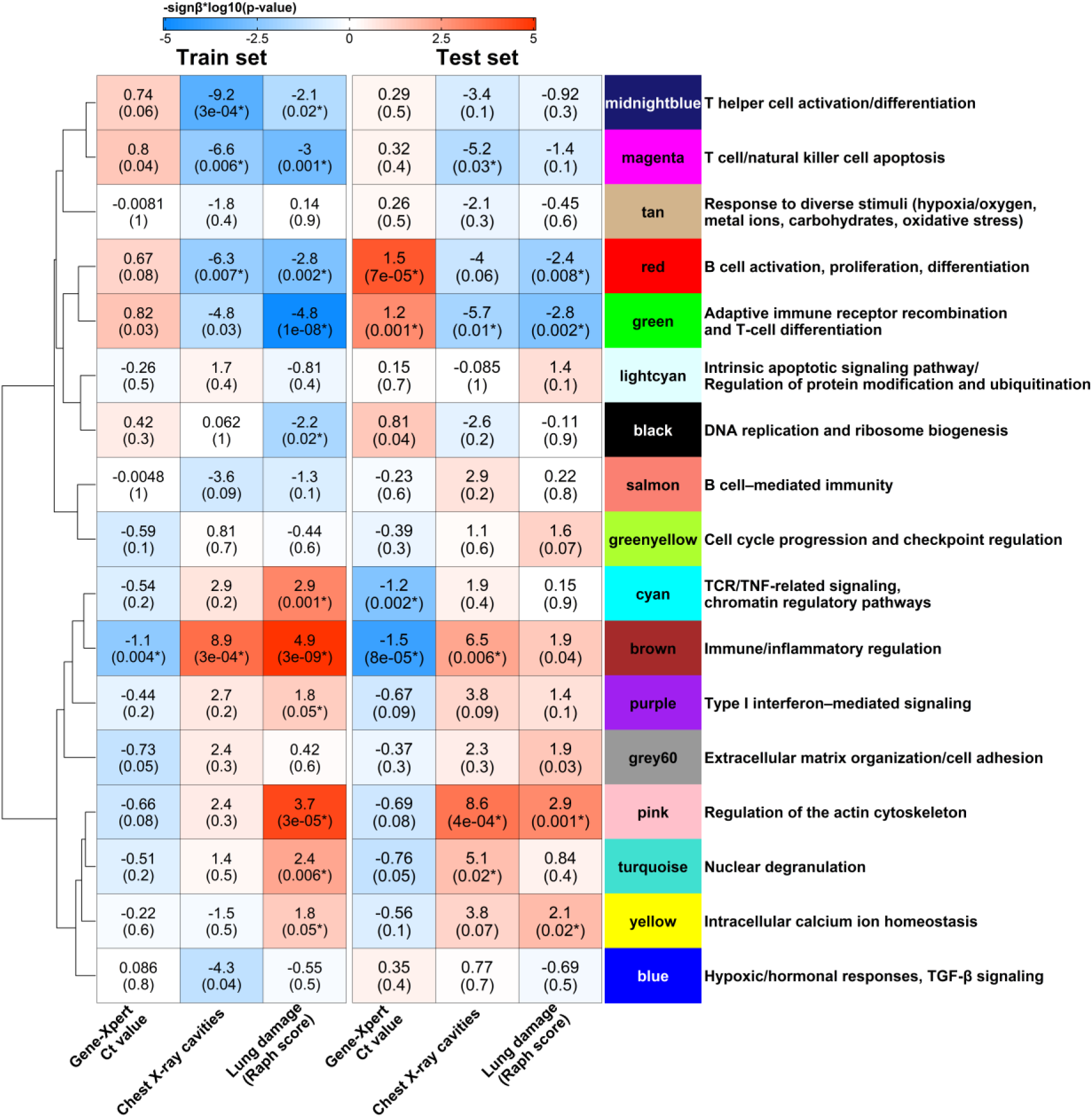
**Whole blood transcriptional modules remain associated with Mtb burden and lung pathology after adjusted for symptom duration.** Heatmap showing the associations between WGCNA modules and Mtb burden (GeneXpert Ct values), chest X-ray cavities, and lung damage (Raph score) in the training and test sets. Each cell displays the slope estimate (with unadjusted p-value, * indicates FDR < 0·1) derived from linear regression (for bacterial burden), logistic regression (for chest X-ray cavity), or beta regression (for lung proportion damage), with module’s eigengene (PC1) and log_2_(symptom duration+1) as the covariates. Blue shading indicates negative slopes, while red shading indicates positive slopes. Modules are organized based on hierarchical clustering of module eigengene correlations, as shown by the dendrogram on the left. Each module was annotated with the most highly correlated biological pathway (from GO-BP, KEGG, or REACTOME) with its eigengene (PC1).

**Figure S4.**
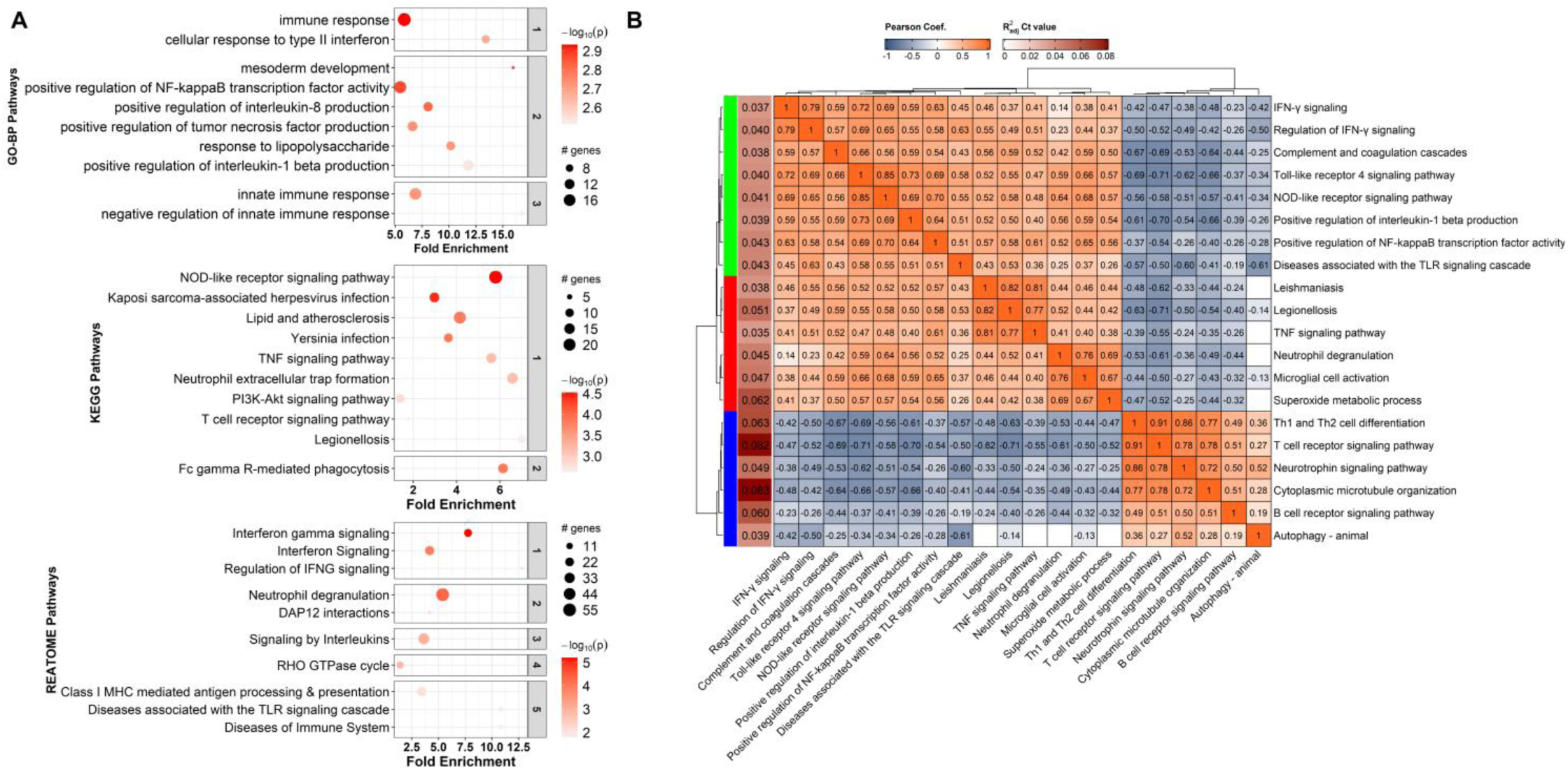
Pathway analysis of the module brown and their correlations. **(A)** Bubble chart presenting the pathway enrichment analysis results for brown module. The results are plotted using bubble charts, where the y-axis lists the names of top-10 lowest p-value pathways separately for Gene Ontology - Biological Process (GO-BP), Kyoto Encyclopedia of Genes and Genomes (KEGG) and REACTOME databases. The x-axis indicates the fold enrichment values, which measure the degree of overrepresentation of genes in each pathway. Higher fold enrichment values suggest stronger associations with the input gene set. Each pathway is depicted as a bubble, with size and color representing, respectively, the number of associated genes and the statistical significance of fold enrichment by −log_10_(p-value). Enriched pathways from each database are grouped into sub-panels using hierarchical clustering (kappa statistics and average linkage method). **(B)** Pearson correlation heatmap of top 20 enriched pathways having the highest association with Mtb burden. Hierarchical clustering analysis highlights the grouping of pathways into three sub-groups, including TNF and its related pathways (red), innate immune response (green), and adaptive immune response (blue), based on their correlation patterns. The left vertical brown bar with numbers represent the adjusted R squared values from GAM model of the association between pathway activity and bacterial burden (log_2_Ct value). Cells without significant correlations (p>0·05) are left blank.

**Figure S5.**
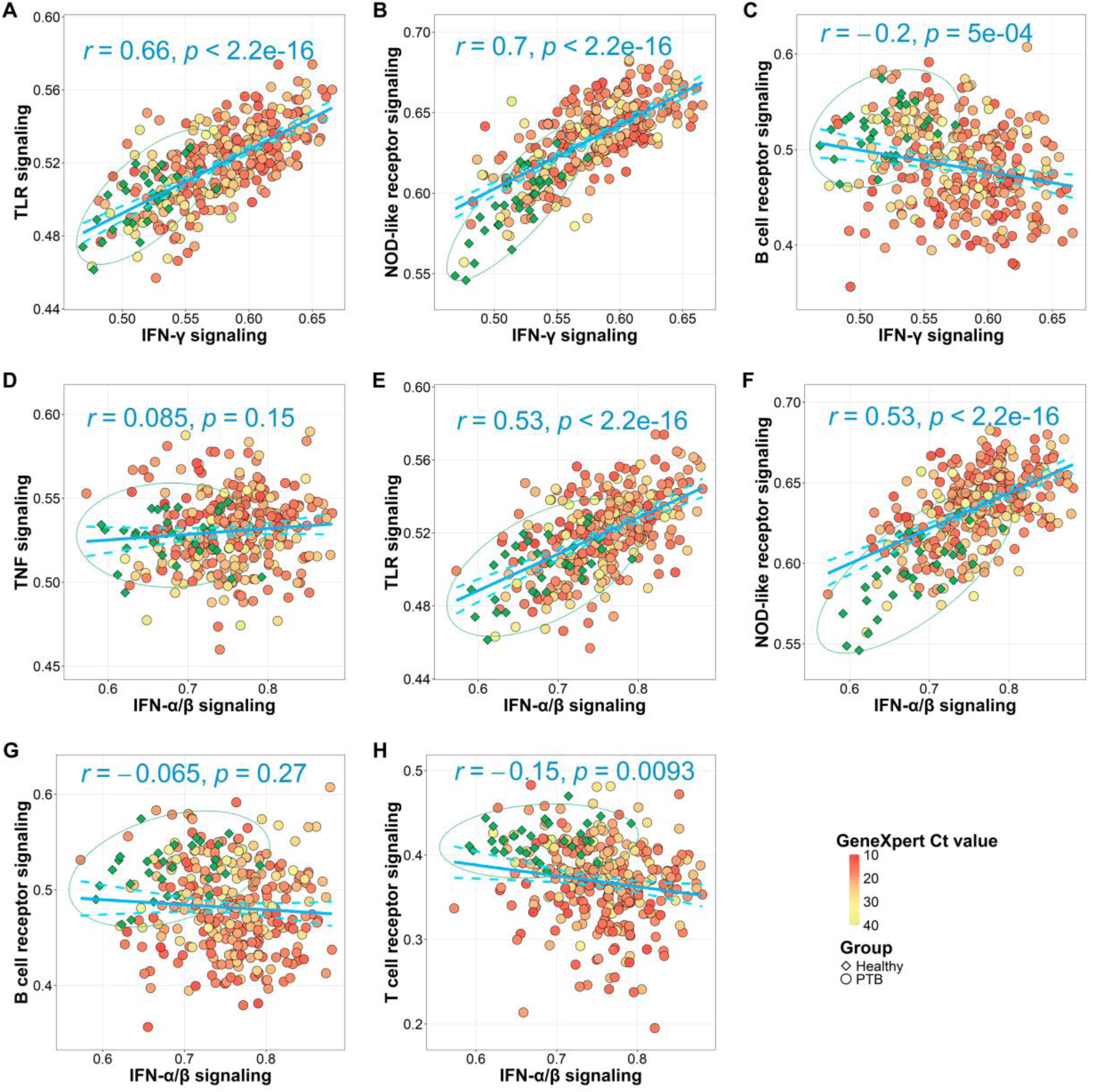
**Associations of immune pathways with Type I and II IFN signaling in PTB**. Scatterplots presenting the relationship between TNF, TLR, NLR, BCR, TCR signaling with IFN-γ and IFN-α/β signaling pathways in PTB patients and healthy controls. Each point represents an individual profile, colored by Ct value for PTB or green for healthy control. A 95% confidence ellipse, based on a bivariate t-distribution, was plotted for the healthy control in each panel. Blue line was the fitted line from the linear regression modeling the relationship of immune pathways with IFN-γ and IFN-α/β signaling pathways in PTB patients (n=292), with Pearson correlation and p-value are indicated on the plot. The two blue dashed lines present 95% confident interval of the estimated trend.

**Figure S6:**
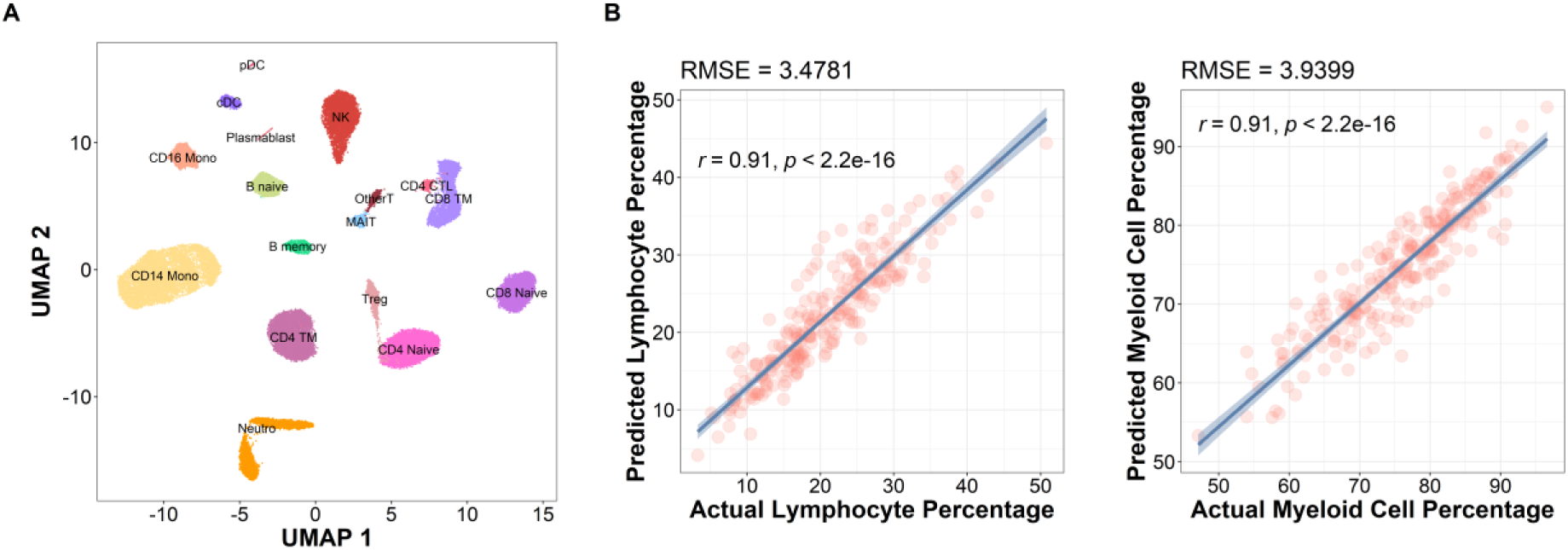
Cell type deconvolution analysis results. **(A)** UMAP visualization of 17 cell subtypes identified in the reference dataset. Each point represents a single cell, with cells grouped into clusters based on their transcriptional profiles and were annotated by subtypes. **(B)** The scatterplots show the predicted cell proportion (y-axis) of Lymphocyte and Myeloid cells compared to the true proportion (x-axis) for PTB patients (n=295). Root mean square error (RMSE) measures the total deviation between the predicted and true proportion for each cell type, with lower value indicating better estimation. Blue line represents the linear regression analysis with shaded region indicating 95% confidence interval. The Pearson correlation coefficient (r) and p-value are also indicated on the plot, showing the significance of the fitted line.

**Figure S7.**
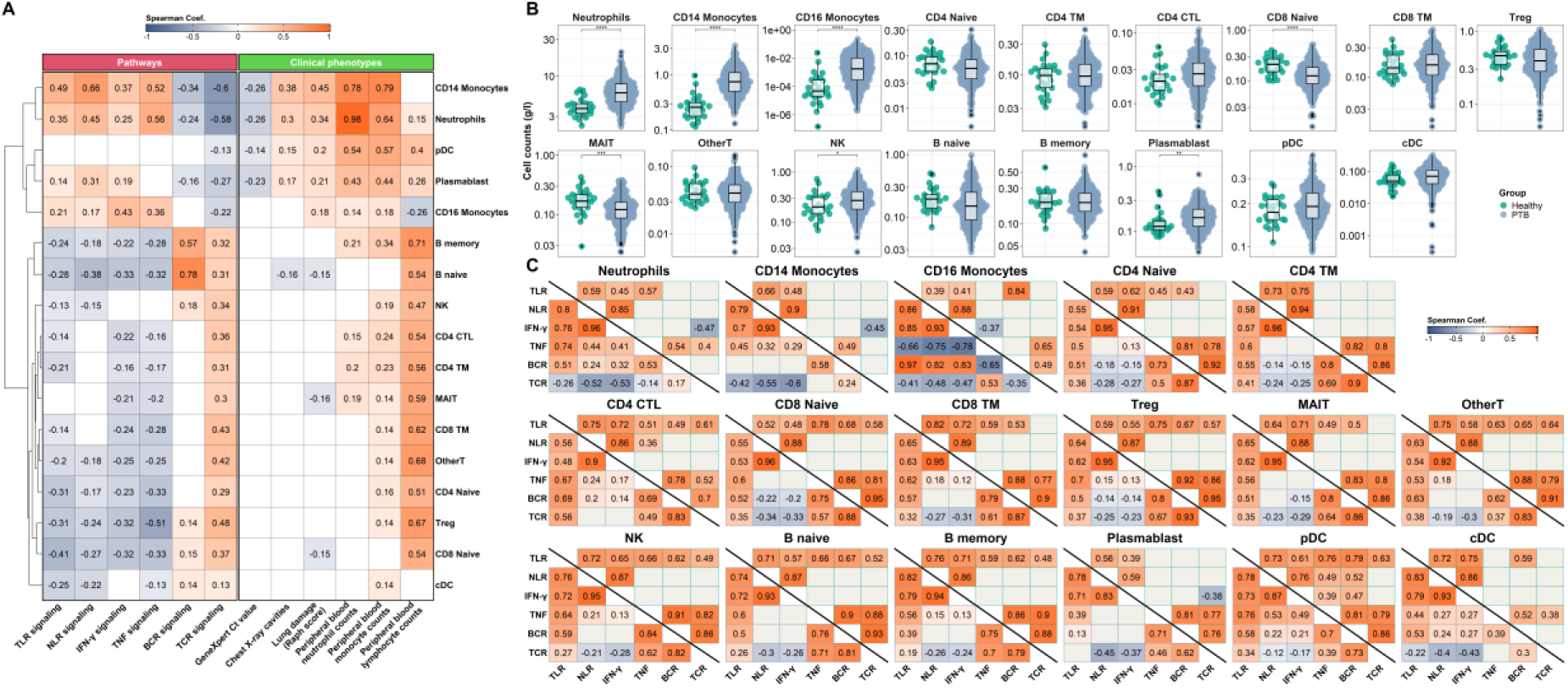
**Association of deconvoluted cell population with immune pathways and clinical phenotypes in PTB**. **(A)** Correlation between estimated counts of 17 cell subtypes with immune pathways and clinical characteristics in PTB patients, including Mtb burden (GeneXpert Ct), lung damage, and blood cell counts (lymphocytes, monocytes, neutrophils). Significant Spearman’s correlation (p < 0·05) were visualised in the heatmap while insignificant correlations were left blank. The order of cell-type was based on hierarchical clustering from correlation coefficient between cell-types. **(B)** Bee swarm boxplots showing comparison of estimated cell counts (g/l) of 17 cell subtypes between healthy control and PTB. Distributions were compared based on Wilcoxon rank sum test, with significance level denoted as * (p≤0·05), ** (p≤0·01), *** (p≤0·001), **** (p≤0·0001), without correction for multiple testing. **(C)** Correlation between immune pathways (ssGSEA) within each cell type of PTB patients (lower triangle heatmap) and healthy controls (upper triangle heatmap). Significant Spearman’s correlations (p < 0·05) were visualised in the heatmap while insignificant correlations were left blank with grey color.

**Figure S8.**
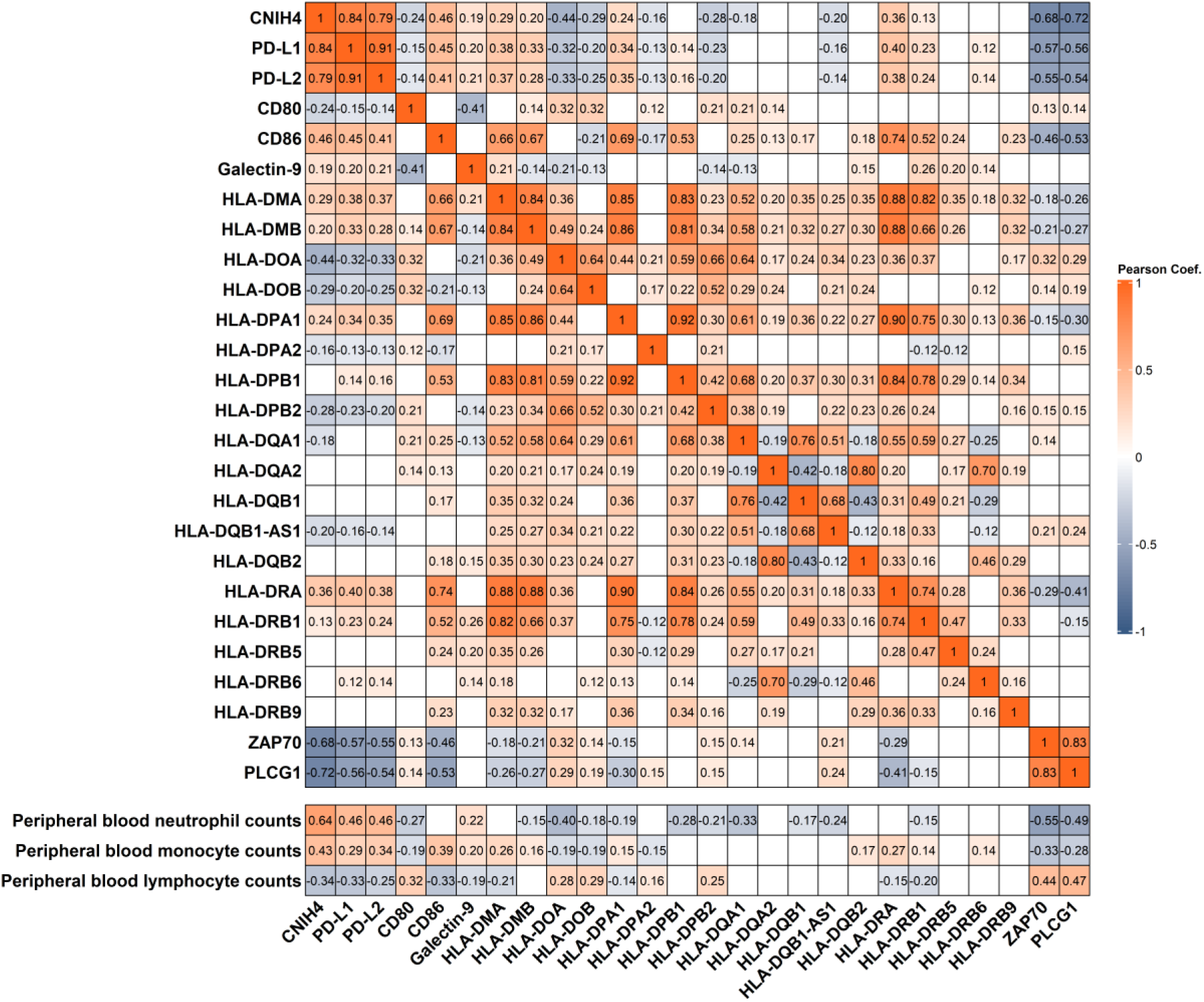
**Correlation between CNIH4, immune check points and immune cell counts** Heatmap showing pairwise Pearson correlation between of CNIH4 with immune checkpoint ligands, including PD-L1/2, CD80/CD86, Galectin-9, and MHC Class II, two key genes of TCR signaling pathway, ZAP70 and PLC-γ1, and blood immune cell counts. Significant Pearson’s correlations (p < 0·05) were visualised in the heatmap while insignificant correlations were left blank.

**Figure S9.**
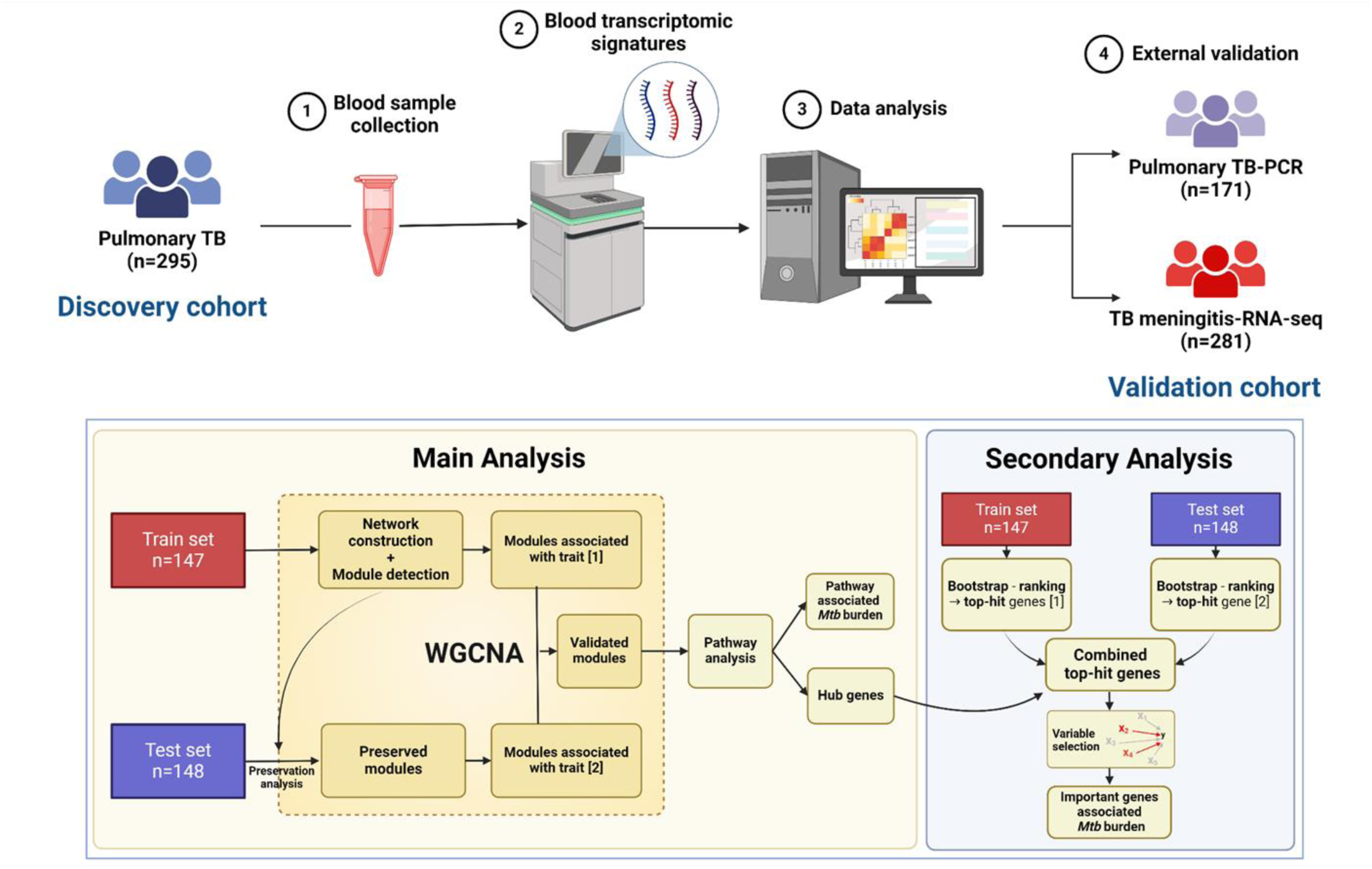
**Schematic diagram of analysis workflow.**

**Figure S10:**
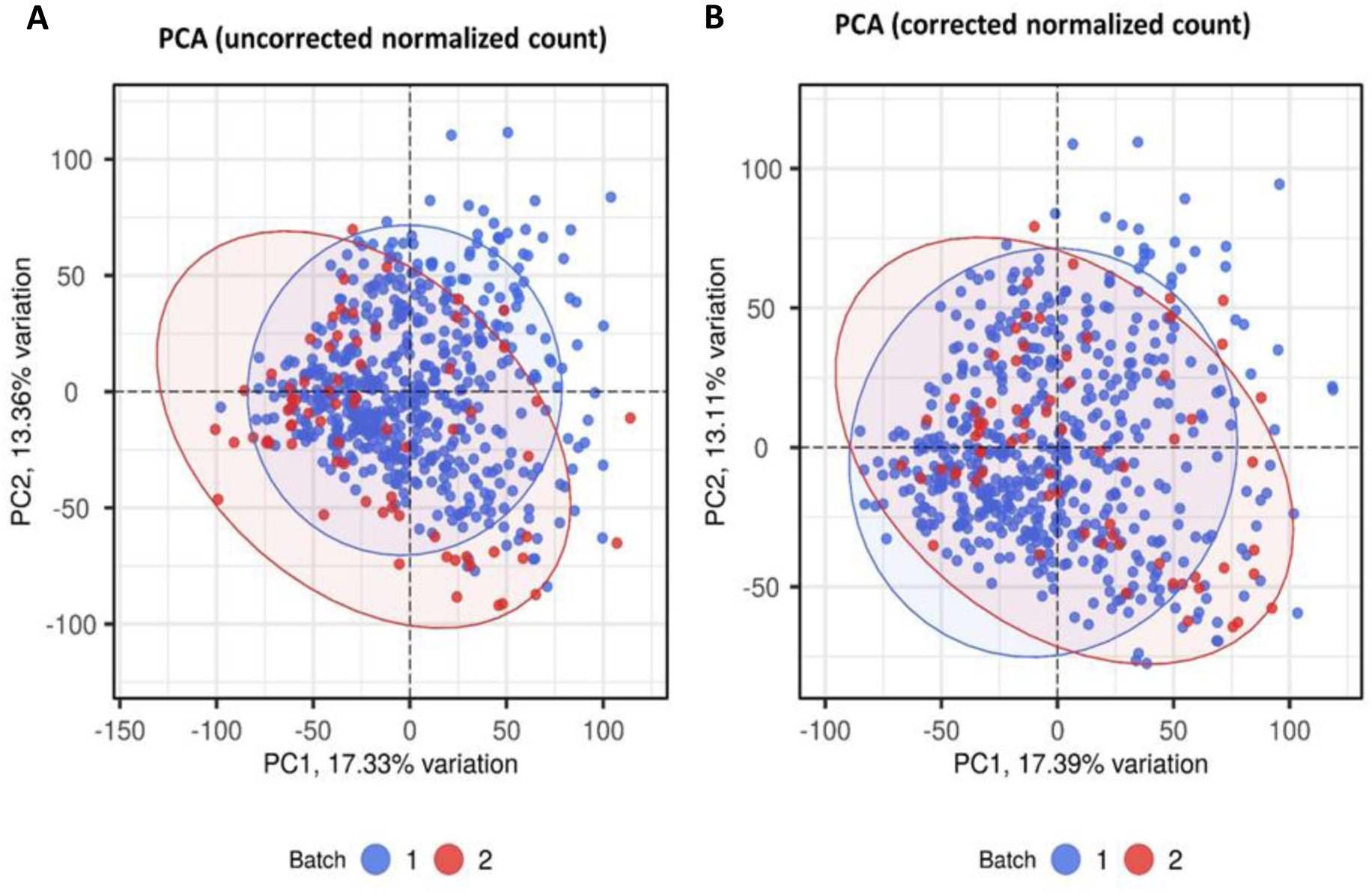
**Batch correction for RNA-seq datasets from healthy, PTB, and TBM cohorts** Principal component analysis (PCA) of RNA-seq datasets from 3 cohorts—295 PTB, 281 HIV-negative or HIV-positive TBM, and 30 healthy controls—before **(A)** and after **(B)** batch correction using the ComBat function from the SVA R package. Each point represents an individual sample, with color indicating the sequencing batch. The x-axis and y-axis represent principal components 1 and 2, respectively. Before batch correction, samples were clustered according to their sequencing batch. After correction, samples from batches 1 and 2 showed substantial overlap, indicating successful removal of batch effects.

**Table S1.**
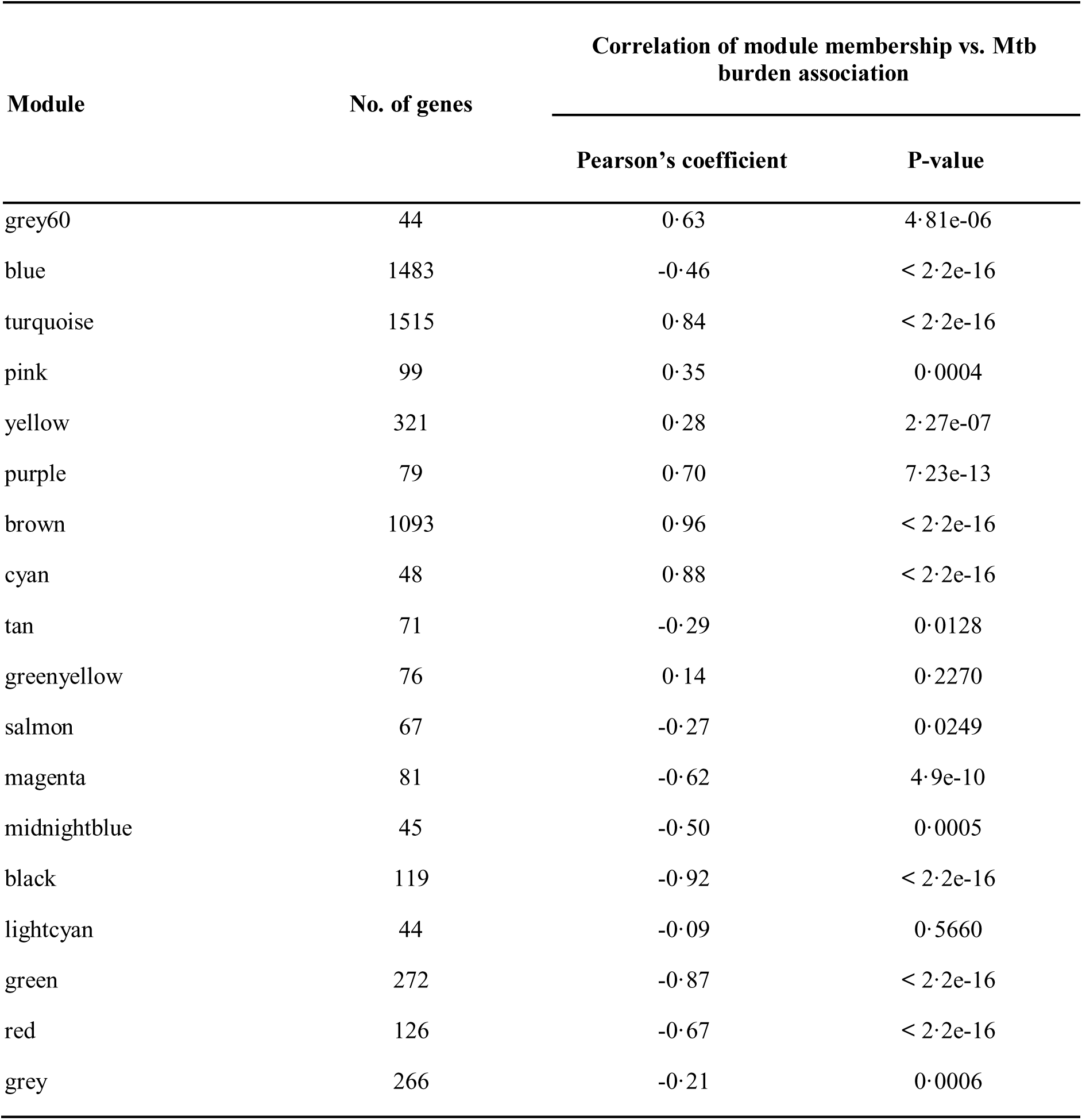
Correlation of gene module membership vs. Mtb burden association.

**Table S2.**
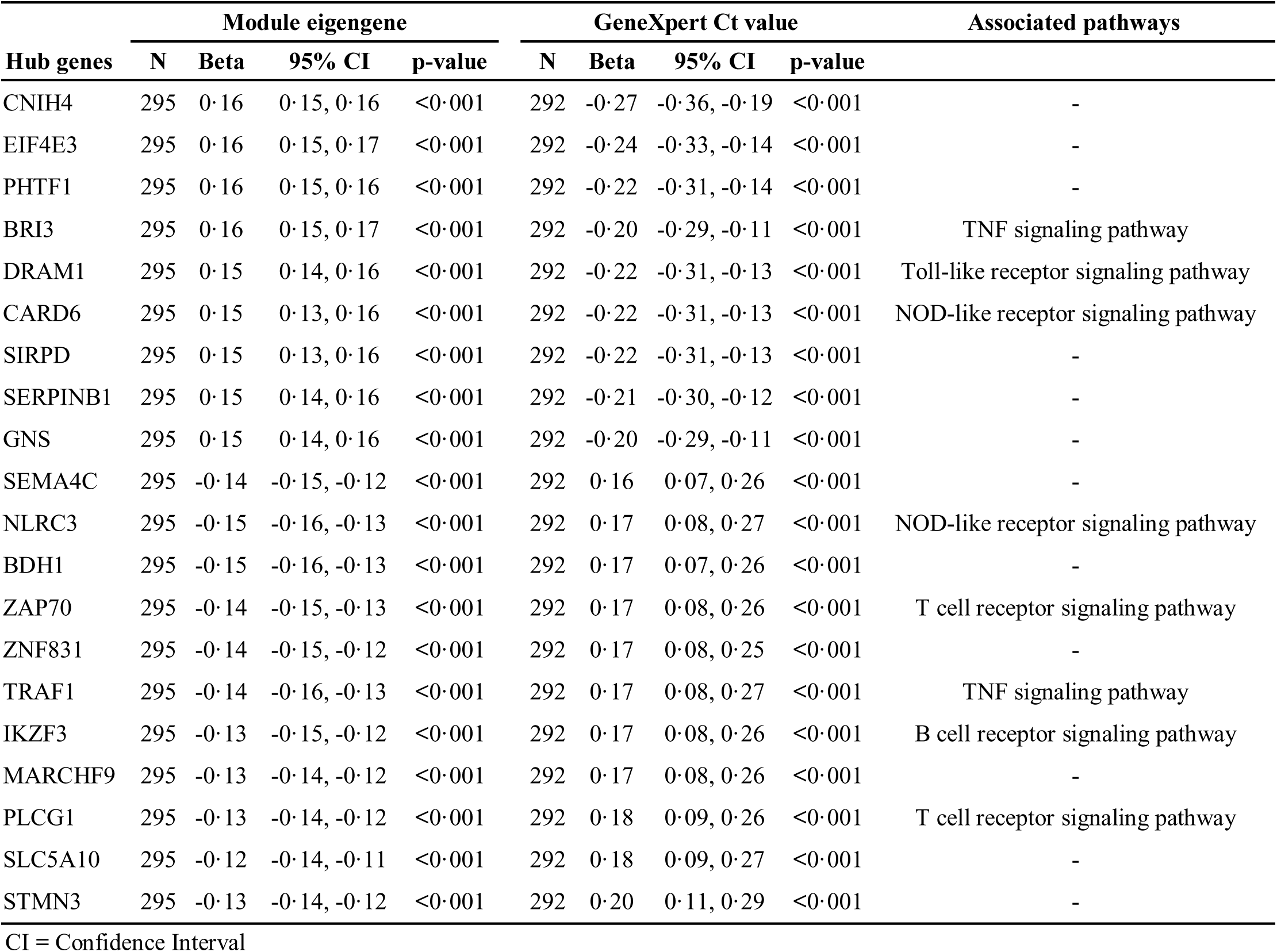
Top 20 hub genes in module brown associated with bacterial burden.

## Notes

### Competing Interest Statement

The authors have declared no competing interest.

## REFERENCE

1. World Health Organitation. Report 2023. Vol. t/malaria/, January. 2024. 2 p.

2. Hai HT, Vinh DN, Thu DDA, Hanh NT, Phu NH, Srinivasan V, et al. Comparison of the Mycobacterium tuberculosis molecular bacterial load assay, microscopy and GeneXpert versus liquid culture for viable bacterial load quantification before and after starting pulmonary tuberculosis treatment. Tuberculosis. 2019;119:101864.

3. Chang VK, Imperial MZ, Phillips PPJ, Velásquez GE, Nahid P, Vernon A, et al. Risk-stratified treatment for drug-susceptible pulmonary tuberculosis. Nat Commun. 2024;15(1):9400.

4. Mortaz E, Adcock IM, Tabarsi P, Masjedi MR, Mansouri D, Velayati AA, et al. Interaction of Pattern Recognition Receptors with Mycobacterium Tuberculosis. J Clin Immunol. 2015;35(1):1–10.

5. Wei L, Liu L, Meng Z, Qi K, Gao X, Feng J, et al. Recognition of Mycobacterium tuberculosis by macrophage Toll-like receptor and its role in autophagy. Inflammation Research. 2024;73(5):753–70.

6. Rastogi S, Briken V. Interaction of Mycobacteria With Host Cell Inflammasomes. Front Immunol. 2022;13(February):1–14.

7. Chandra P, Grigsby SJ, Philips JA. Immune evasion and provocation by Mycobacterium tuberculosis. Nat Rev Microbiol. 2022;20(12):750–66.

8. Silvério D, Gonçalves R, Appelberg R, Saraiva M. Advances on the Role and Applications of Interleukin-1 in Tuberculosis. Vol. 12, mBio. American Society for Microbiology; 2021.

9. Tsao TC, Hong J h, Li LF, Hsieh MJ, Liao SK, Chang KS. Imbalances between tumor necrosis factor-alpha and its soluble receptor forms, and interleukin-1beta and interleukin-1 receptor antagonist in BAL fluid of cavitary pulmonary tuberculosis. Chest. 2000 Jan;117(1):103–9.

10. Amaral EP, Vinhaes CL, Oliveira-De-Souza D, Nogueira B, Akrami KM, Andrade BB. The Interplay between Systemic Inflammation, Oxidative Stress, and Tissue Remodeling in Tuberculosis. Antioxid Redox Signal. 2021;34(6):471–85.

11. Green AM, DiFazio R, Flynn JL. IFN-γ from CD4 T Cells Is Essential for Host Survival and Enhances CD8 T Cell Function during Mycobacterium tuberculosis Infection . The Journal of Immunology. 2013;190(1):270–7.

12. Fabri M, Stenger S, Shin DM, Yuk JM, Liu PT, Realegeno S, et al. Vitamin D is required for IFN-gamma-mediated antimicrobial activity of human macrophages. Sci Transl Med. 2011 Oct;3(104):104ra102.

13. Flynn JL, Goldstein MM, Chan J, Triebold KJ, Pfeffer K, Lowenstein CJ, et al. Tumor necrosis factor-alpha is required in the protective immune response against Mycobacterium tuberculosis in mice. Immunity. 1995 Jun;2(6):561–72.

14. Flynn JL, Chan J, Lin PL. Macrophages and control of granulomatous inflammation in tuberculosis. Mucosal Immunol. 2011;4(3):271–8.

15. Kolloli A, Singh P, Subbian S. Granulomatous response to Mycobacterium tuberculosis infection. Understanding the Host Immune Response Against Mycobacterium tuberculosis Infection. 2018. 41–66 p.

16. Travar M, Petkovic M, Verhaz A. Type I, II, and III Interferons: Regulating Immunity to Mycobacterium tuberculosis Infection. Arch Immunol Ther Exp (Warsz). 2016;64(1):19–31.

17. Ong CWM, Elkington PT, Friedland JS. Tuberculosis, pulmonary cavitation, and matrix metalloproteinases. Am J Respir Crit Care Med. 2014;190(1):9–18.

18. Berry MPR, Graham CM, McNab FW, Xu Z, Bloch SAA, Oni T, et al. An interferon-inducible neutrophil-driven blood transcriptional signature in human tuberculosis. Nature. 2010 Aug;466(7309):973–7.

19. Gideon HP, Phuah J, Junecko BA, Mattila JT. Neutrophils express pro- and anti-inflammatory cytokines in granulomas from Mycobacterium tuberculosis-infected cynomolgus macaques. Mucosal Immunol. 2019;12(6):1370–81.

20. Ravimohan S, Kornfeld H, Weissman D, Bisson GP. Tuberculosis and lung damage: From epidemiology to pathophysiology. European Respiratory Review. 2018;27(147).

21. Tiwari D, Martineau AR. Inflammation-mediated tissue damage in pulmonary tuberculosis and host-directed therapeutic strategies. Semin Immunol. 2023;65(November 2022):101672.

22. Boehme CC, Nabeta P, Hillemann D, Nicol MP, Shenai S, Krapp F, et al. Rapid molecular detection of tuberculosis and rifampin resistance. N Engl J Med. 2010 Sep;363(11):1005–15.

23. Donovan J, Thu DDA, Phu NH, Dung VTM, Quang TP, Nghia HDT, et al. Xpert MTB/RIF Ultra versus Xpert MTB/RIF for the diagnosis of tuberculous meningitis: a prospective, randomised, diagnostic accuracy study. Lancet Infect Dis. 2020 Mar;20(3):299–307.

24. Nhu NTQ, Heemskerk D, Thu DDA, Chau TTH, Mai NTH, Nghia HDT, et al. Evaluation of GeneXpert MTB/RIF for diagnosis of tuberculous meningitis. J Clin Microbiol. 2014 Jan;52(1):226–33.

25. Fradejas I, Ontañón B, Muñoz-Gallego I, Ramírez-Vela MJ, López-Roa P. The value of xpert MTB/RIF-generated CT values for predicting the smear status of patients with pulmonary tuberculosis. J Clin Tuberc Other Mycobact Dis. 2018 Dec;13:9–12.

26. Thuong NTT, Vinh DN, Hai HT, Thu DDA, Nhat LTH, Heemskerk D, et al. Pretreatment cerebrospinal fluid bacterial load correlates with inflammatory response and predicts neurological events during tuberculous meningitis treatment. Journal of Infectious Diseases. 2019 Feb 23;219(6):986–95.

27. Hai HT, Thanh Hoang Nhat L, Tram TTB, Vinh DD, Nath AP, Donovan J, et al. Whole blood transcriptional profiles and the pathogenesis of tuberculous meningitis. Elife. 2024 Oct 30;13:2023.10.06.561265.

28. Conesa A, Madrigal P, Tarazona S, Gomez-Cabrero D, Cervera A, McPherson A, et al. A survey of best practices for RNA-seq data analysis. Genome Biol. 2016 Jan;17:13.

29. Yalamanchili HK, Wan YW, Liu Z. Data Analysis Pipeline for RNA-seq Experiments: From Differential Expression to Cryptic Splicing. Curr Protoc Bioinformatics. 2017 Sep;59:11.15.1–11.15.21.

30. Andrews S. FastQC: a quality control tool for high throughput sequence data. 2010;

31. Bolger AM, Lohse M, Usadel B. Trimmomatic: a flexible trimmer for Illumina sequence data. Bioinformatics. 2014 Aug;30(15):2114–20.

32. Dobin A, Davis CA, Schlesinger F, Drenkow J, Zaleski C, Jha S, et al. STAR: ultrafast universal RNA-seq aligner. Bioinformatics. 2013 Jan;29(1):15–21.

33. Liao Y, Smyth GK, Shi W. featureCounts: an efficient general purpose program for assigning sequence reads to genomic features. Bioinformatics. 2014 Apr;30(7):923–30.

34. Love MI, Huber W, Anders S. Moderated estimation of fold change and dispersion for RNA-seq data with DESeq2. Genome Biol. 2014;15(12):550.

35. Leek JT, Johnson WE, Parker HS, Jaffe AE, Storey JD. The sva package for removing batch effects and other unwanted variation in high-throughput experiments. Bioinformatics. 2012 Mar;28(6):882–3.

36. Ahmed M, Thirunavukkarasu S, Rosa BA, Thomas KA, Das S, Rangel-Moreno J, et al. Immune correlates of tuberculosis disease and risk translate across species. Sci Transl Med. 2020;12(528):1–17.

37. Simes RJ. An improved Bonferroni procedure for multiple tests of significance.pdf. Vol. 73, Biometrika. 1986. p. 751–4.

38. Langfelder P, Zhang B, Horvath S. Defining clusters from a hierarchical cluster tree: the Dynamic Tree Cut package for R. Bioinformatics. 2007;24(5):719–20.

39. Langfelder P, Luo R, Oldham MC, Horvath S. Is My Network Module Preserved and Reproducible? PLoS Comput Biol. 2011 Jan 20;7(1):e1001057.

40. Benjamini Y, Hochberg Y. Controlling the False Discovery Rate : A Practical and Powerful Approach to Multiple Testing. Journal of the Royal Statistical Society. 1995;57(1):289–300.

41. Horvath S, Dong J. Geometric interpretation of gene coexpression network analysis. PLoS Comput Biol. 2008;4(8):24–6.

42. Aleksander SA, Balhoff J, Carbon S, Cherry JM, Drabkin HJ, Ebert D, et al. The Gene Ontology knowledgebase in 2023. Genetics. 2023;224(1):1–14.

43. Kanehisa M, Furumichi M, Sato Y, Kawashima M, Ishiguro-Watanabe M. KEGG for taxonomy-based analysis of pathways and genomes. Nucleic Acids Res. 2023;51(D1):D587–92.

44. Milacic M, Beavers D, Conley P, Gong C, Gillespie M, Griss J, et al. The Reactome Pathway Knowledgebase 2024. Nucleic Acids Res. 2024 Jan;52(D1):D672–8.

45. Ulgen E, Ozisik O, Sezerman OU. PathfindR: An R package for comprehensive identification of enriched pathways in omics data through active subnetworks. Front Genet. 2019;10(SEP):1–33.

46. Chatr-Aryamontri A, Oughtred R, Boucher L, Rust J, Chang C, Kolas NK, et al. The BioGRID interaction database: 2017 update. Nucleic Acids Res. 2017;45(D1):D369–79.

47. Ideker T, Ozier O, Schwikowski B, Siegel AF. Discovering regulatory and signalling circuits in molecular interaction networks. Bioinformatics. 2002 Jul 1;18(suppl_1):S233–40.

48. Benjamini Y, Yekutieli D. THE CONTROL OF THE FALSE DISCOVERY RATE IN MULTIPLE TESTING UNDER DEPENDENCY. The annals of Statistics. 2001;29(4):1165–88.

49. Chu T, Wang Z, Pe’er D, Danko CG. Cell type and gene expression deconvolution with BayesPrism enables Bayesian integrative analysis across bulk and single-cell RNA sequencing in oncology. Nat Cancer. 2022;3(4):505–17.

50. Hu M, Chikina M. InstaPrism: an R package for fast implementation of BayesPrism. Bioinformatics. 2024;40(7):btae440.

51. Hao Y, Hao S, Andersen-Nissen E, Mauck WM 3rd, Zheng S, Butler A, et al. Integrated analysis of multimodal single-cell data. Cell. 2021 Jun;184(13):3573–3587.e29.

52. Wigerblad G, Cao Q, Brooks S, Naz F, Gadkari M, Jiang K, et al. Single-Cell Analysis Reveals the Range of Transcriptional States of Circulating Human Neutrophils. J Immunol. 2022 Aug;209(4):772–82.

53. Erdmann-Pham DD, Fischer J, Hong J, Song YS. Likelihood-based deconvolution of bulk gene expression data using single-cell references. Genome Res. 2021 Oct;31(10):1794–806.

54. Meinshausen N, Bühlmann P. Stability selection. J R Stat Soc Series B Stat Methodol. 2010;72(4):417–73.

55. Najjingo I, Muttamba W, Kirenga BJ, Nalunjogi J, Bakesiima R, Olweny F, et al. Comparison of GeneXpert cycle threshold values with smear microscopy and culture as a measure of mycobacterial burden in five regional referral hospitals of Uganda- A cross-sectional study. PLoS One. 2019;14(5):e0216901.

56. Ralph AP, Ardian M, Wiguna A, Maguire GP, Becker NG, Drogumuller G, et al. A simple, valid, numerical score for grading chest x-ray severity in adult smear-positive pulmonary tuberculosis. Thorax. 2010 Oct 1;65(10):863–9.

57. Donovan ML, Schultz TE, Duke TJ, Blumenthal A. Type I interferons in the pathogenesis of tuberculosis: Molecular drivers and immunological consequences. Front Immunol. 2017;8(NOV).

58. Shanmuganathan G, Orujyan D, Narinyan W, Poladian N, Dhama S, Parthasarathy A, et al. Role of Interferons in Mycobacterium tuberculosis Infection. Clin Pract. 2022;12(5):788–96.

59. Singh A, Mohan A, Dey AB, Mitra DK. Inhibiting the programmed death 1 pathway rescues Mycobacterium tuberculosis-specific interferon γ-producing T cells from apoptosis in patients with pulmonary tuberculosis. Journal of Infectious Diseases. 2013;208(4):603–15.

60. Han C, Godfrey V, Liu Z, Han Y, Liu L, Peng H, et al. The AIM2 and NLRP3 inflammasomes trigger IL-1-mediated antitumor effects during radiation. Sci Immunol. 2021;6(59):1–9.

61. Hupa KJ, Stein K, Schneider R, Lysson M, Schneiker B, Hornung V, et al. AIM2 inflammasome-derived IL-1β induces postoperative ileus in mice. Sci Rep. 2019;9(1):1–13.

62. Di Paolo NC, Shafiani S, Day T, Papayannopoulou T, Russell DW, Iwakura Y, et al. Interdependence between Interleukin-1 and Tumor Necrosis Factor Regulates TNF-Dependent Control of Mycobacterium tuberculosis Infection. Immunity [Internet]. 2015 Dec 15;43(6):1125–36. Available from: 10.1016/j.immuni.2015.11.016

63. Sauvageau E, Rochdi MD, Oueslati M, Hamdan FF, Percherancier Y, Simpson JC, et al. CNIH4 interacts with newly synthesized GPCR and controls their export from the endoplasmic reticulum. Traffic. 2014;15(4):383–400.

64. Li F, Chen D, Zeng Q, Du Y. Possible Mechanisms of Lymphopenia in Severe Tuberculosis. Microorganisms. 2023;11(11).

65. Cubillos-Angulo JM, Nogueira BMF, Arriaga MB, Barreto-Duarte B, Araújo-Pereira M, Fernandes CD, et al. Host-directed therapies in pulmonary tuberculosis: Updates on anti-inflammatory drugs. Front Med (Lausanne). 2022;9(September):1–15.

66. Garcia-Diaz A, Shin DS, Moreno BH, Saco J, Escuin-Ordinas H, Rodriguez GA, et al. Interferon Receptor Signaling Pathways Regulating PD-L1 and PD-L2 Expression. Cell Rep. 2017;19(6):1189–201.

67. Moon JW, Kong SK, Kim BS, Kim HJ, Lim H, Noh K, et al. IFNγ induces PD-L1 overexpression by JAK2/STAT1/IRF-1 signaling in EBV-positive gastric carcinoma. Sci Rep. 2017;7(1):1–13.

68. Day CL, Abrahams DA, Bunjun R, Stone L, de Kock M, Walzl G, et al. PD-1 Expression on Mycobacterium tuberculosis-Specific CD4 T Cells Is Associated With Bacterial Load in Human Tuberculosis. Front Immunol. 2018;9(August):1–18.

69. Sakai S, Kauffman KD, Sallin MA, Sharpe AH, Young HA, Ganusov V V, et al. CD4 T Cell-Derived IFN-γ Plays a Minimal Role in Control of Pulmonary Mycobacterium tuberculosis Infection and Must Be Actively Repressed by PD-1 to Prevent Lethal Disease. PLoS Pathog. 2016 May;12(5):e1005667.

70. Wang L, Ma H, Wen Z, Niu L, Chen X, Liu H, et al. Single-cell RNA-sequencing reveals heterogeneity and intercellular crosstalk in human tuberculosis lung. Journal of Infection. 2023;87(5):373–84.

71. Belmont J, Gu T, Mudd A, Salomon AR. A PLC-γ1 Feedback Pathway Regulates Lck Substrate Phosphorylation at the T-Cell Receptor and SLP-76 Complex. J Proteome Res. 2017 Aug 4;16(8):2729–42.

72. Sheppard KA, Fitz LJ, Lee JM, Benander C, George JA, Wooters J, et al. PD-1 inhibits T-cell receptor induced phosphorylation of the ZAP70/CD3ζ signalosome and downstream signaling to PKCθ. FEBS Lett. 2004;574(1–3):37–41.

73. Juárez E, Carranza C, Hernández–Sánchez F, León–Contreras JC, Hernández–Pando R, Escobedo D, et al. NOD2 enhances the innate response of alveolar macrophages to Mycobacterium tuberculosis in humans. Eur J Immunol. 2012 Apr 24;42(4):880–9.

74. Negroni A, Pierdomenico M, Cucchiara S, Stronati L. NOD2 and inflammation: current insights. J Inflamm Res. 2018 Feb;Volume 11:49–60.

75. Pandey AK, Yang Y, Jiang Z, Fortune SM, Coulombe F, Behr MA, et al. NOD2, RIP2 and IRF5 Play a Critical Role in the Type I Interferon Response to Mycobacterium tuberculosis. PLoS Pathog. 2009 Jul 3;5(7):e1000500.

76. Hornung V, Ablasser A, Charrel-Dennis M, Bauernfeind F, Horvath G, Caffrey DanielR, et al. AIM2 recognizes cytosolic dsDNA and forms a caspase-1-activating inflammasome with ASC. Nature [Internet]. 2009;458(7237):514–8. Available from: 10.1038/nature07725

77. Saiga H, Kitada S, Shimada Y, Kamiyama N, Okuyama M, Makino M, et al. Critical role of AIM2 in Mycobacterium tuberculosis infection. Int Immunol. 2012 Oct 1;24(10):637–44.

78. Sharma BR, Karki R, Kanneganti T. Role of AIM2 inflammasome in inflammatory diseases, cancer and infection. Eur J Immunol. 2019 Nov 14;49(11):1998–2011.

79. Yuk K, Lim J, Moon HY. Comparing the effects of resistance and aerobic exercise on mood-related symptoms and EEG activity in young healthy adults: A non-randomized pilot study. Ment Health Phys Act. 2024 Oct;27:100626.

