## Supplemental Information for "Whole blood transcriptional responses associated with bacterial burden in pulmonary tuberculosis"

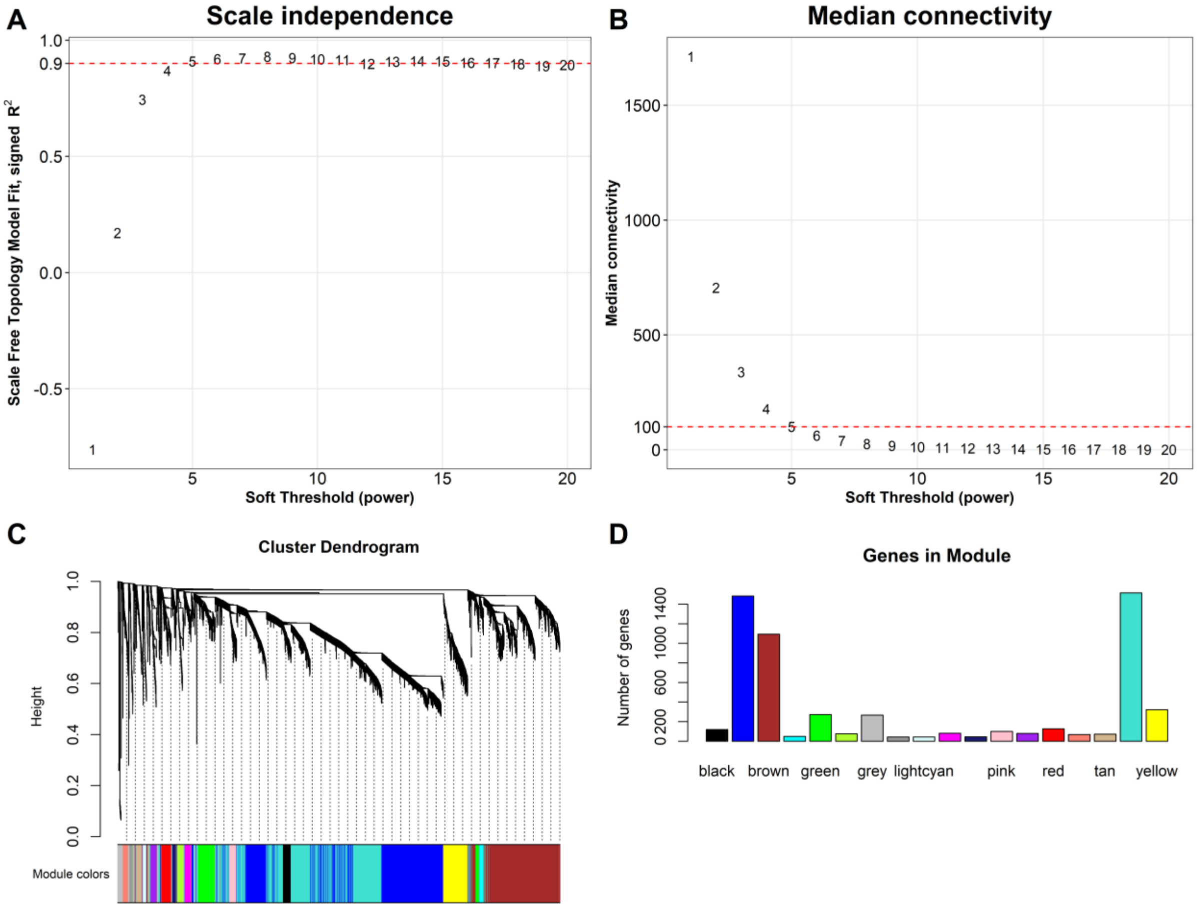


**Figure S1. Construction of WGCNA in train set**

Analysis of network topology for various soft-thresholding powers of the top 5,850 most variant genes based on the scale-free network model. A soft-thresholding power was optimally determined to balance between enforcing scale-free topology and preserving meaningful gene-gene interactions. **(A)** The scale-free fit index (y-axis) as a function of the soft-thresholding power (x-axis). The red horizontal dash line corresponds to $R^2 = 0 \cdot 9$ and soft-thresholding power β=5, which was chosen for the construction gene-expression network. **(B)** The median connectivity on the y-axis as a function of the soft-thresholding power on the x-axis. The adjacency matrix of the scale-free network between genes was determined as $A = (a_{ij})$, where $a_{ij} = |cor(gene_i, gene_j)|^{\beta}$. Red line indicates the optimal median connectivity using soft-thresholding power β=5. **(C)** Dendrogram resulted from the hierarchical clustering analysis using topological overlap of the adjacency matrix A served as a dissimilarity metric. Each cluster was referred as a module and assigned with a color. In total, we identified 17 transcriptional modules in the training set, excluding the grey module as it consisted of non-connected genes. **(D)** Bar-plots indicate the number of genes contained in each module.

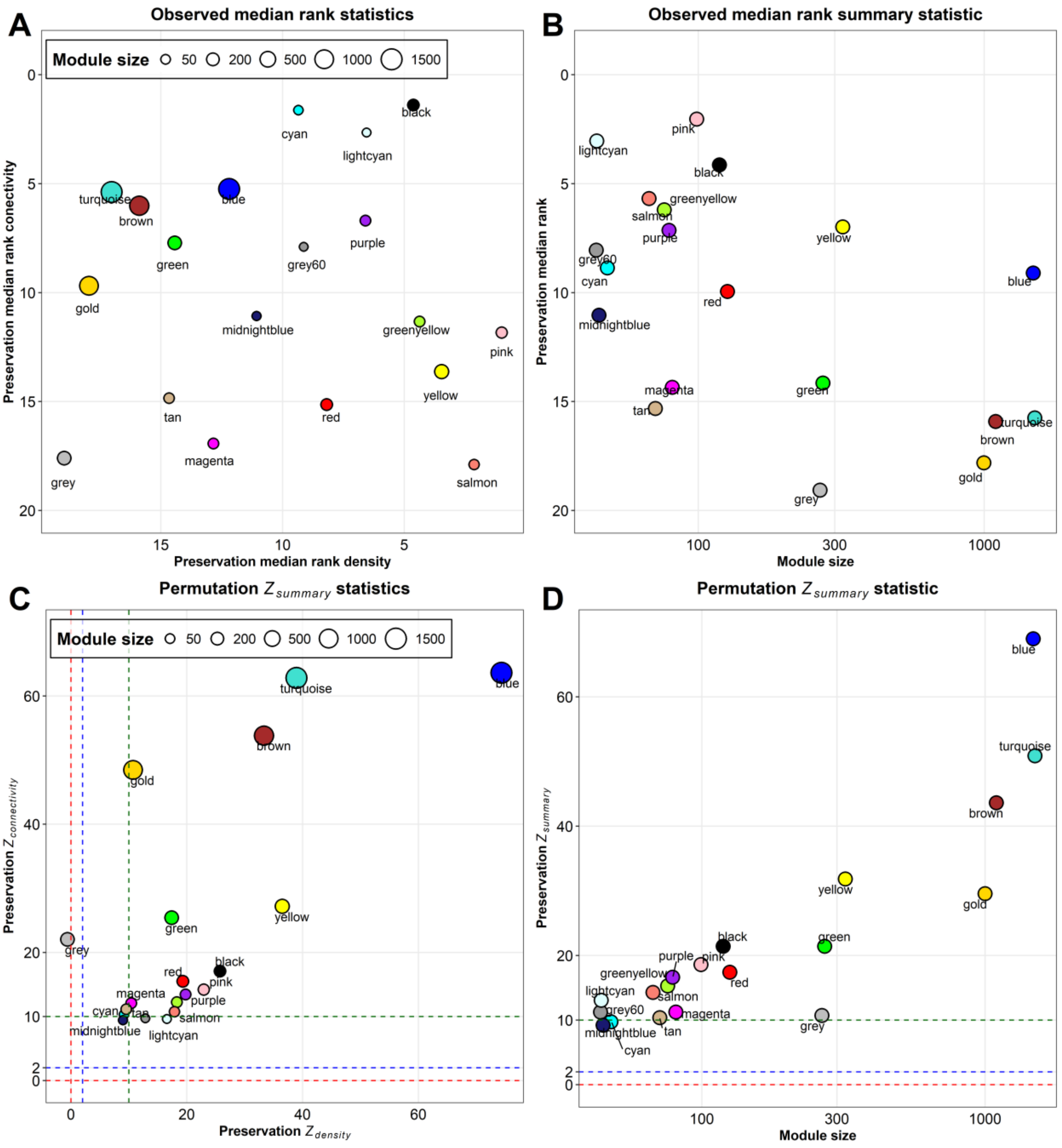


**Figure S2. Preservation of discovery modules in validation set**

**(A)** The median rank preservation statistics of the modules. Each module was represented by a point, labeled with the corresponding color and name. The X and Y-axis represent the median rank density and connectivity of observed preservation statistics per module, respectively. A low preservation median rank value indicates a high level of preservation. The gold module was an artificial module comprised of 500 randomly selected genes. The grey module consisted of non-connected genes identified in the WGCNA analysis of the discovery cohort. **(B)** The median rank summary statistic by module size showing gold and grey modules exhibited the highest median rank, indicating low preservation, which ensured our preservation analysis controlling well the background noise signal. **(C)** The $Z_{density}$ and $Z_{connectivity}$ preservation statistic of the modules. Each module was represented by a point, labeled with the corresponding color and name. The X-axis and Y-axis represents the $Z_{density}$ and $Z_{connectivity}$ statistic of each module based on 1,000 permutations of module labels, respectively. **(D)** The $Z_{summary}$ preservation statistic by module size. In both figures, the blue and green dash lines correspond to the threshold of Z statistics of 2 and 10, where ≥10: High preservation; 2<Z<10: Moderate preservation; <2: Low preservation. Overall, using the threshold of Z≥10 across density, connectivity and summary statistics, 12 out of 17 identified modules in the training set were validated in the validation set.

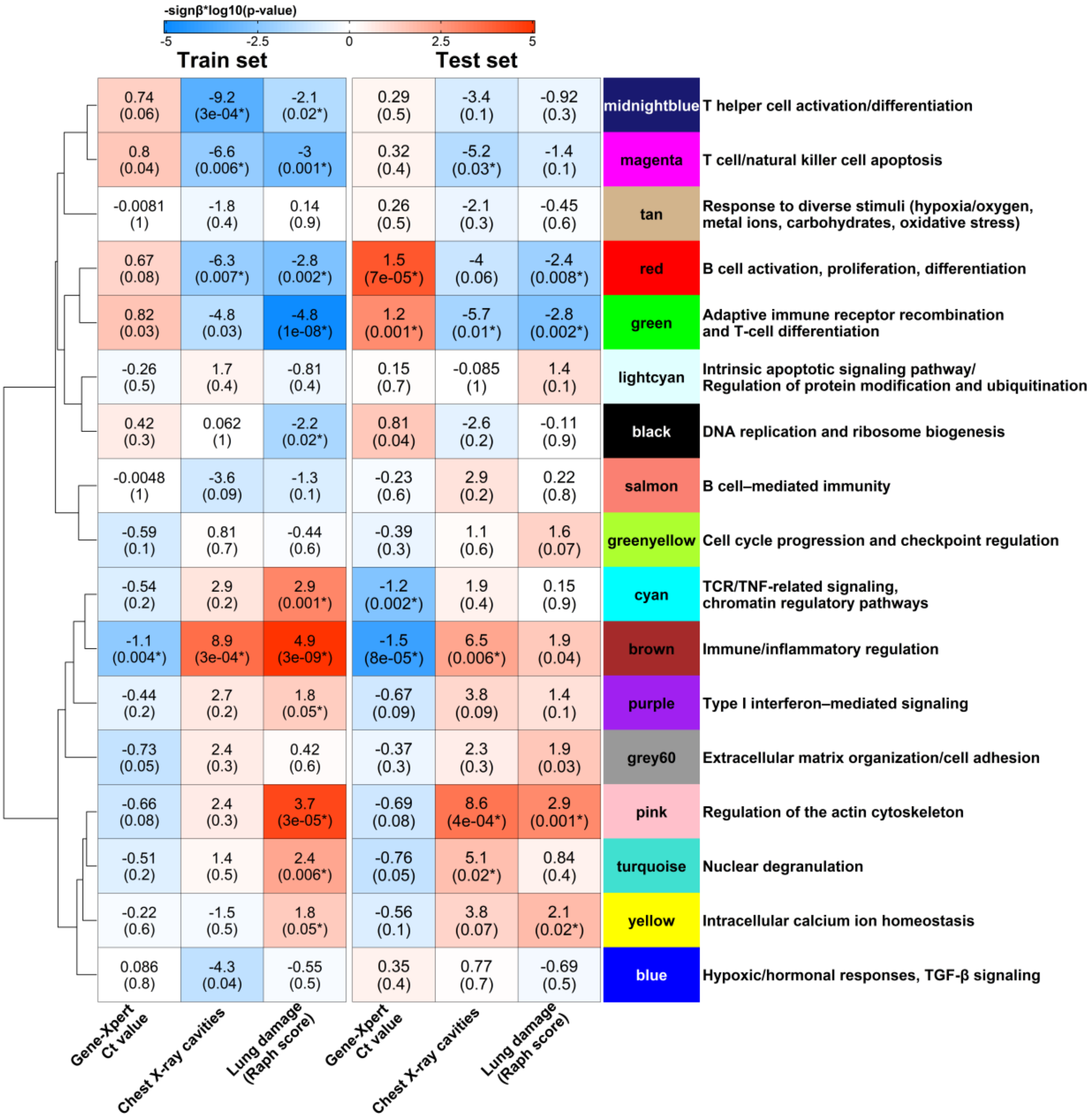


**Figure S3. Whole blood transcriptional modules remain associated with Mtb burden and lung pathology after adjusted for symptom duration.**

Heatmap showing the associations between WGCNA modules and Mtb burden (GeneXpert Ct values), chest X-ray cavities, and lung damage (Raph score) in the training and test sets. Each cell displays the slope estimate (with unadjusted p-value, * indicates FDR < 0·1) derived from linear regression (for bacterial burden), logistic regression (for chest X-ray cavity), or beta regression (for lung proportion damage), with module's eigengene (PC1) and $\log_2$(symptom duration+1) as the covariates. Blue shading indicates negative slopes, while red shading indicates positive slopes. Modules are organized based on hierarchical clustering of module eigengene correlations, as shown by the dendrogram on the left. Each module was annotated with the most highly correlated biological pathway (from GO-BP, KEGG, or REACTOME) with its eigengene (PC1).

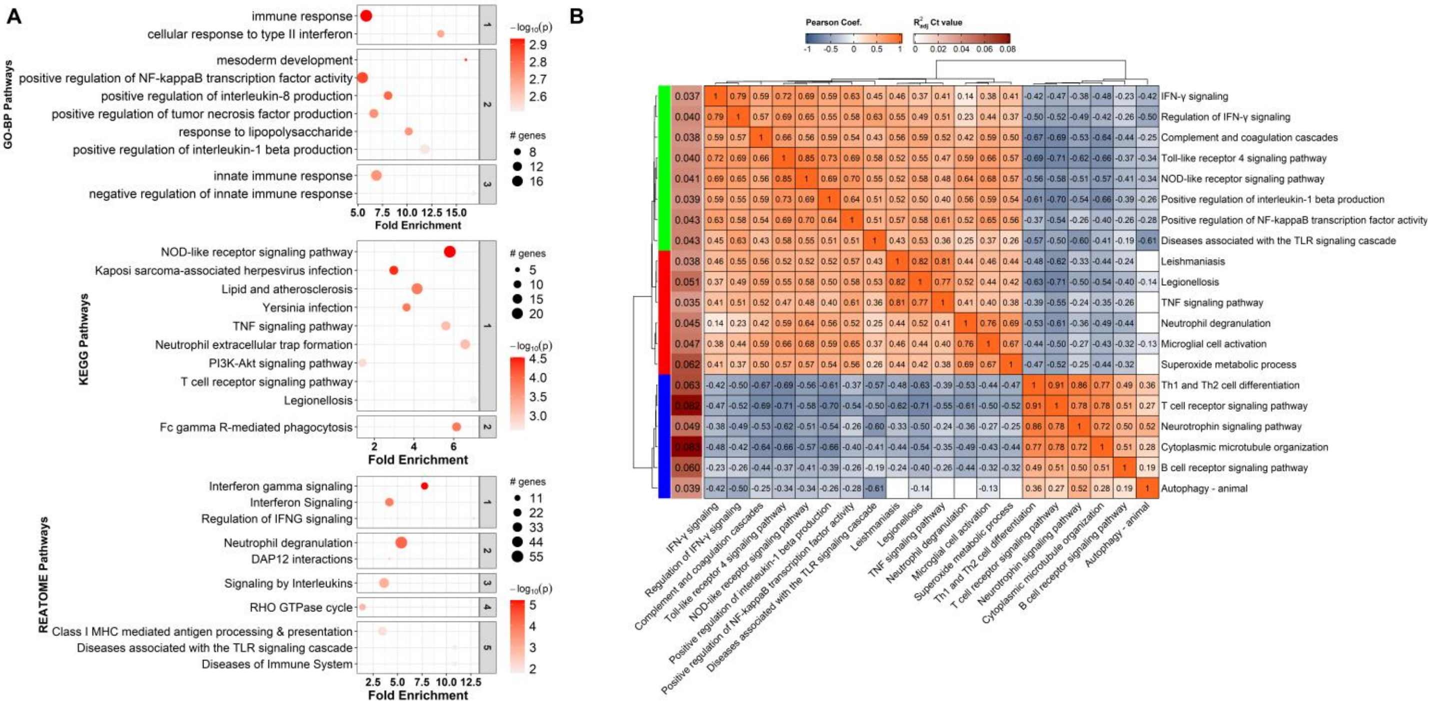


**Figure S4. Pathway analysis of the module brown and their correlations**

**(A)** Bubble chart presenting the pathway enrichment analysis results for brown module. The results are plotted using bubble charts, where the y-axis lists the names of top-10 lowest p-value pathways separately for Gene Ontology - Biological Process (GO-BP), Kyoto Encyclopedia of Genes and Genomes (KEGG) and REACTOME databases. The x-axis indicates the fold enrichment values, which measure the degree of overrepresentation of genes in each pathway. Higher fold enrichment values suggest stronger associations with the input gene set. Each pathway is depicted as a bubble, with size and color representing, respectively, the number of associated genes and the statistical significance of fold enrichment by -$\log_{10}$(p-value). Enriched pathways from each database are grouped into sub-panels using hierarchical clustering (kappa statistics and average linkage method). **(B)** Pearson correlation heatmap of top 20 enriched pathways having the highest association with Mtb burden. Hierarchical clustering analysis highlights the grouping of pathways into three sub-groups, including TNF and its related pathways (red), innate immune response (green), and adaptive immune response (blue), based on their correlation patterns. The left vertical brown bar with numbers represent the adjusted R squared values from GAM model of the association between pathway activity and bacterial burden ($\log_2$Ct value). Cells without significant correlations (p>0·05) are left blank.

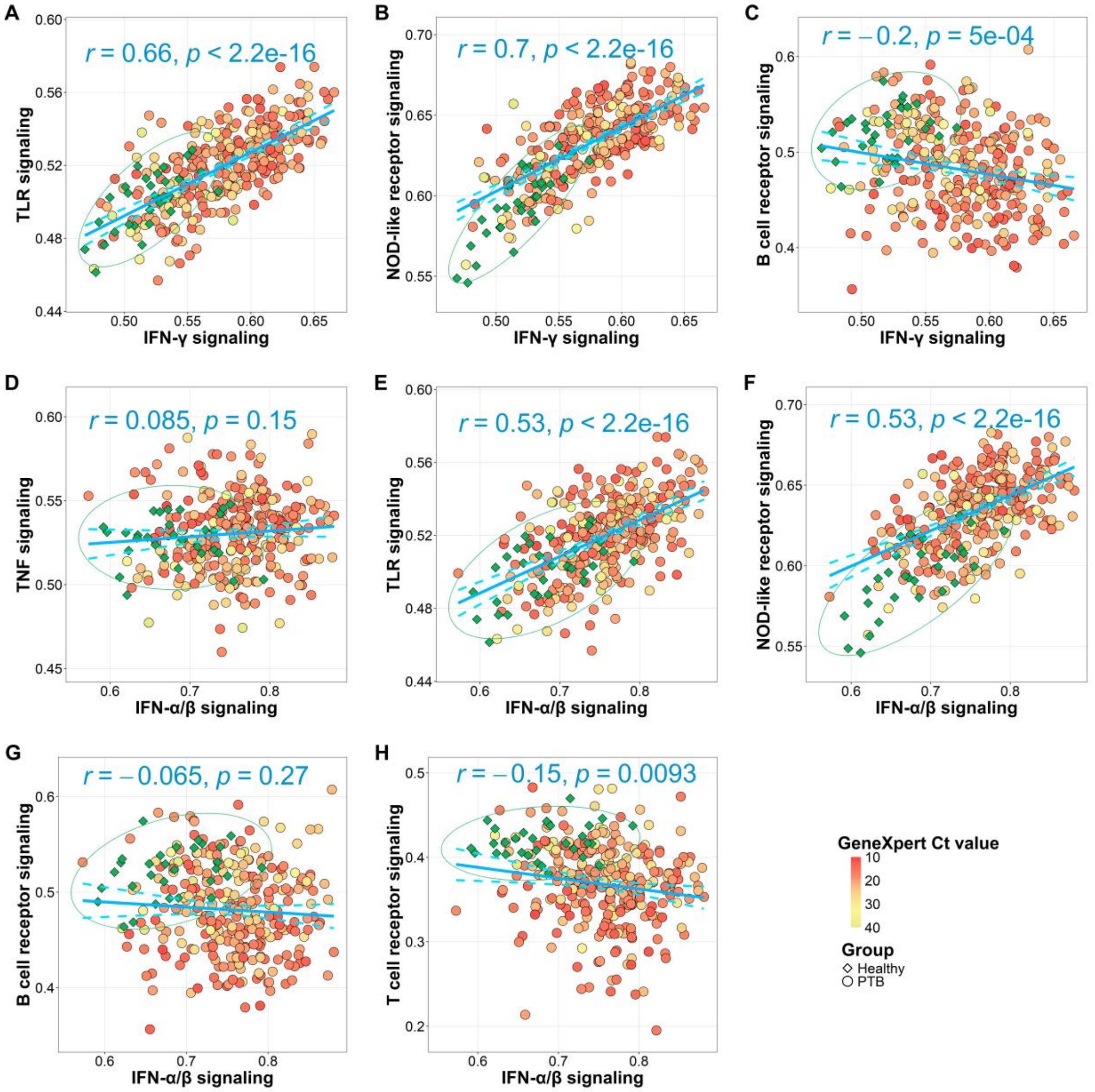


**Figure S5. Associations of immune pathways with Type I and II IFN signaling in PTB**.

Scatterplots presenting the relationship between TNF, TLR, NLR, BCR, TCR signaling with IFN-γ and IFN-α/β signaling pathways in PTB patients and healthy controls. Each point represents an individual profile, colored by Ct value for PTB or green for healthy control. A 95% confidence ellipse, based on a bivariate t-distribution, was plotted for the healthy control in each panel. Blue line was the fitted line from the linear regression modeling the relationship of immune pathways with IFN-γ and IFN-α/β signaling pathways in PTB patients (n=292), with Pearson correlation and p-value are indicated on the plot. The two blue dashed lines present 95% confident interval of the estimated trend.

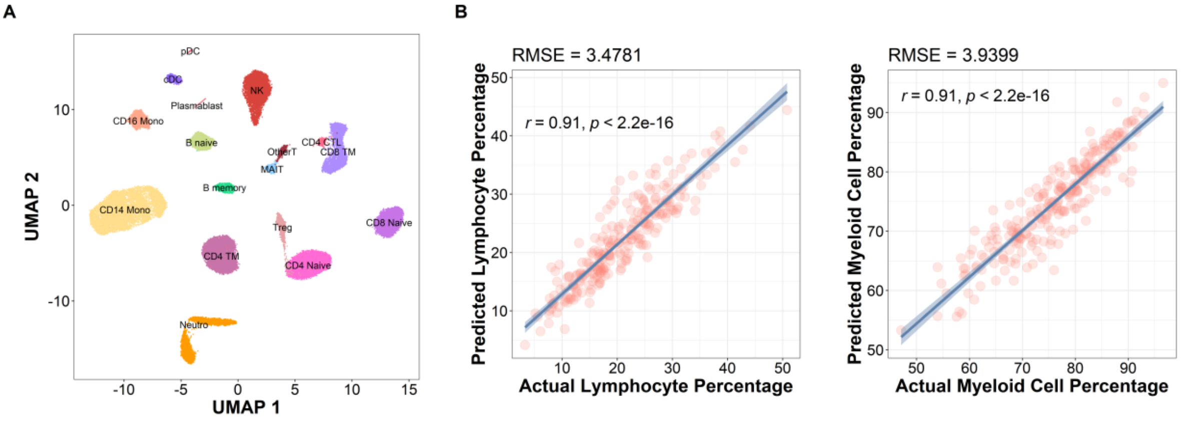


**Figure S6: Cell type deconvolution analysis results.**

**(A)** UMAP visualization of 17 cell subtypes identified in the reference dataset. Each point represents a single cell, with cells grouped into clusters based on their transcriptional profiles and were annotated by subtypes. **(B)** The scatterplots show the predicted cell proportion (y-axis) of Lymphocyte and Myeloid cells compared to the true proportion (x-axis) for PTB patients (n=295). Root mean square error (RMSE) measures the total deviation between the predicted and true proportion for each cell type, with lower value indicating better estimation. Blue line represents the linear regression analysis with shaded region indicating 95% confidence interval. The Pearson correlation coefficient (r) and p-value are also indicated on the plot, showing the significance of the fitted line.

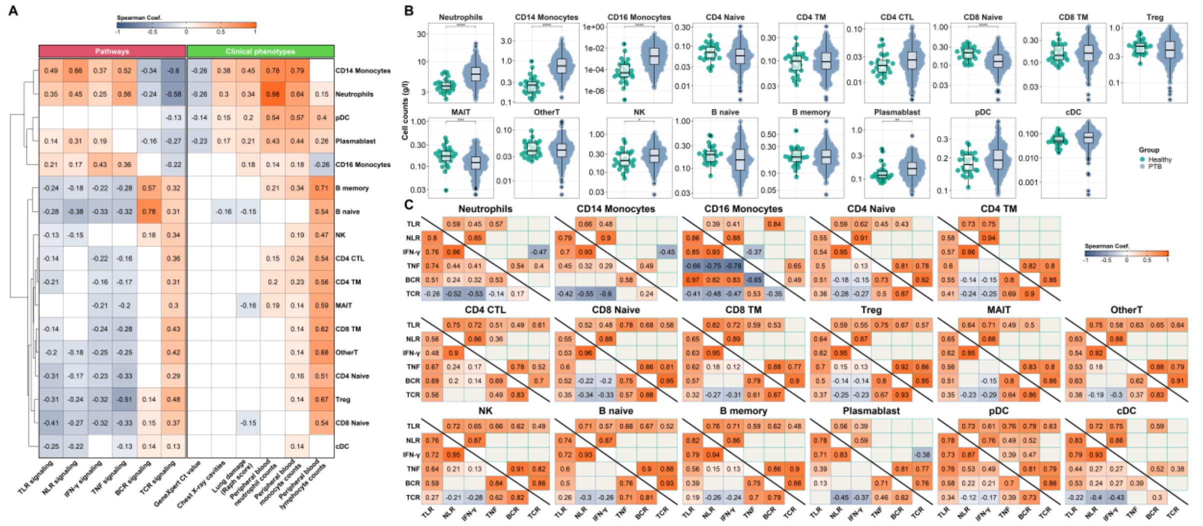

**Figure S7. Association of deconvoluted cell population with immune pathways and clinical phenotypes in PTB**.

**(A)** Correlation between estimated counts of 17 cell subtypes with immune pathways and clinical characteristics in PTB patients, including Mtb burden (GeneXpert Ct), lung damage, and blood cell counts (lymphocytes, monocytes, neutrophils). Significant Spearman's correlation ($p < 0{\cdot}05$) were visualised in the heatmap while insignificant correlations were left blank. The order of cell-type was based on hierarchical clustering from correlation coefficient between cell-types. **(B)** Bee swarm boxplots showing comparison of estimated cell counts (g/l) of 17 cell subtypes between healthy control and PTB. Distributions were compared based on Wilcoxon rank sum test, with significance level denoted as * ($p \le 0{\cdot}05$), ** ($p \le 0{\cdot}01$), *** ($p \le 0{\cdot}001$), **** ($p \le 0{\cdot}0001$), without correction for multiple testing. **(C)** Correlation between immune pathways (ssGSEA) within each cell type of PTB patients (lower triangle heatmap) and healthy controls (upper triangle heatmap). Significant Spearman's correlations ($p < 0{\cdot}05$) were visualised in the heatmap while insignificant correlations were left blank with grey color.

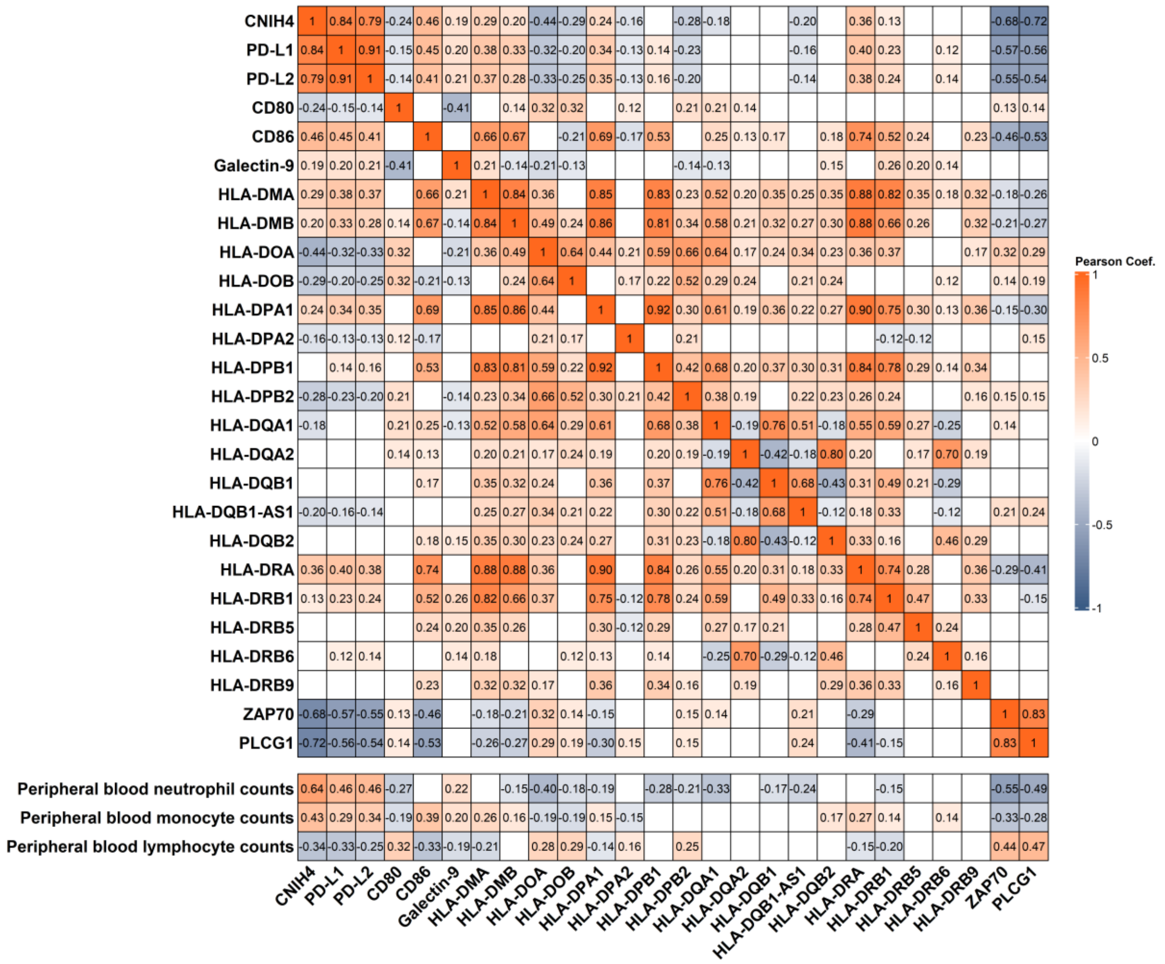

**Figure S8. Correlation between CNIH4, immune check points and immune cell counts**

Heatmap showing pairwise Pearson correlation between of CNIH4 with immune checkpoint ligands, including PD-L1/2, CD80/CD86, Galectin-9, and MHC Class II, two key genes of TCR signaling pathway, ZAP70 and PLC-γ1, and blood immune cell counts. Significant Pearson's correlations ($p < 0{\cdot}05$) were visualised in the heatmap while insignificant correlations were left blank.

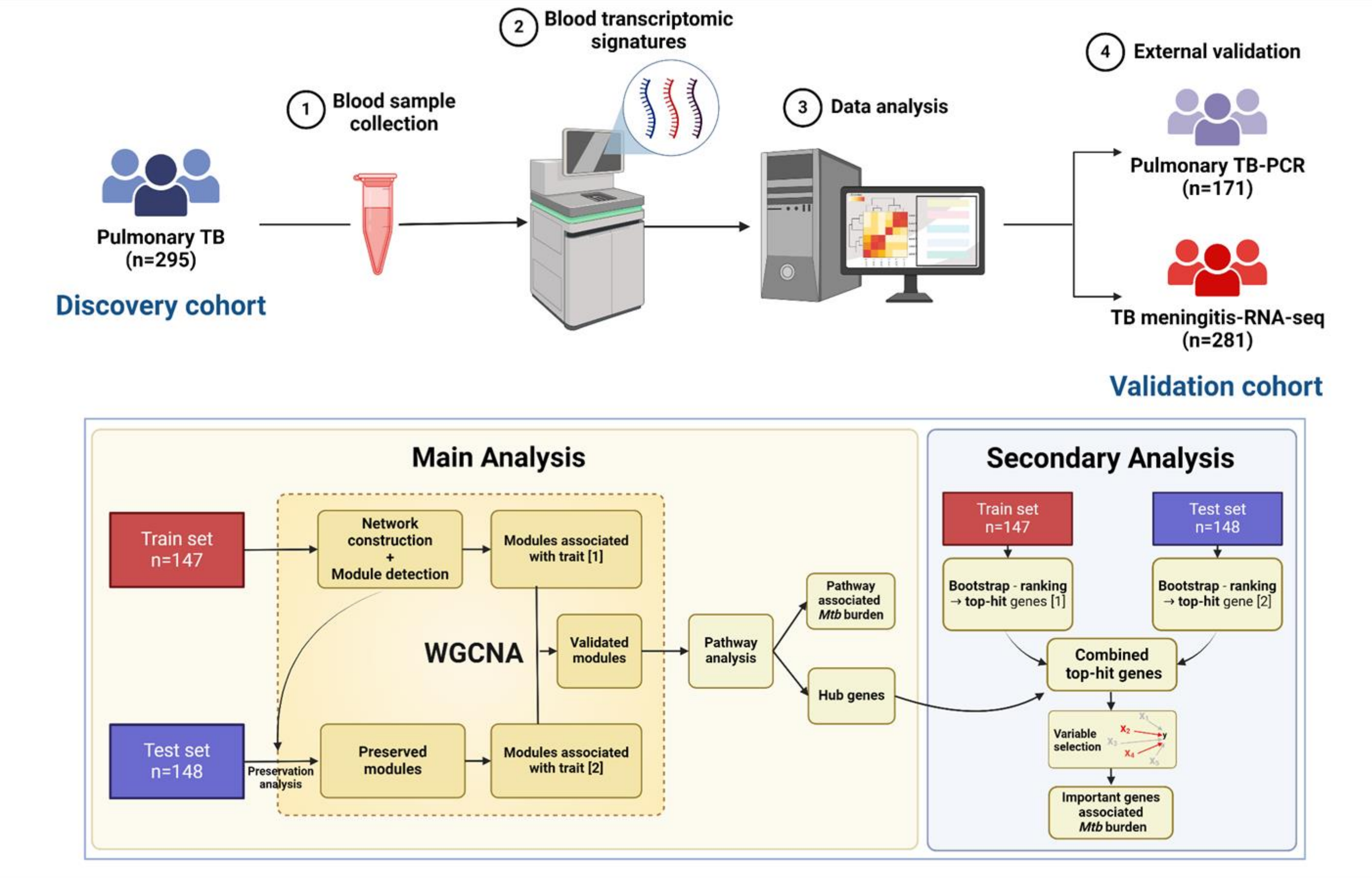


**Figure S9. Schematic diagram of analysis workflow.**

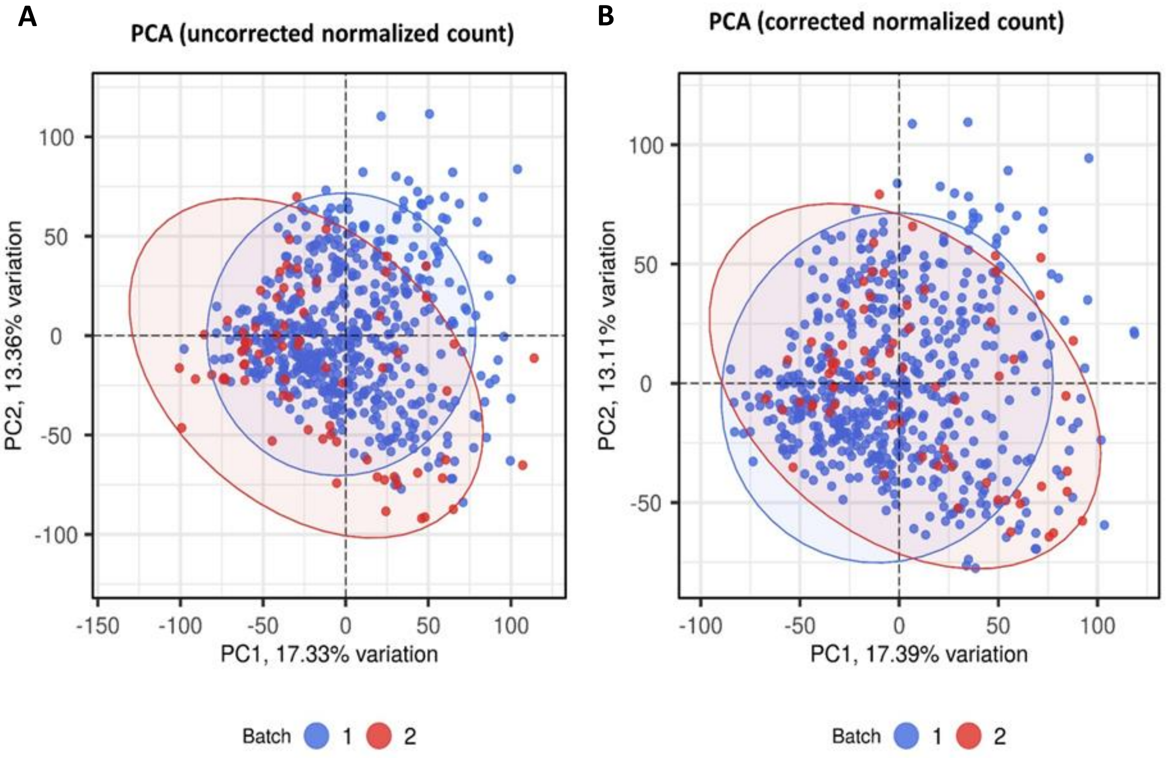


**Figure S10: Batch correction for RNA-seq datasets from healthy, PTB, and TBM cohorts**

Principal component analysis (PCA) of RNA-seq datasets from 3 cohorts—295 PTB, 281 HIV-negative or HIV-positive TBM, and 30 healthy controls—before **(A)** and after **(B)** batch correction using the ComBat function from the SVA R package. Each point represents an individual sample, with color indicating the sequencing batch. The x-axis and y-axis represent principal components 1 and 2, respectively. Before batch correction, samples were clustered according to their sequencing batch. After correction, samples from batches 1 and 2 showed substantial overlap, indicating successful removal of batch effects.

**Table S1. Correlation of gene module membership vs. Mtb burden association**

| Module | No. of genes | Correlation of module membership vs. Mtb burden association | |
|---|---|---|---|
| | | Pearson's coefficient | P-value |
| grey60 | 44 | 0·63 | 4·81e-06 |
| blue | 1483 | -0·46 | < 2·2e-16 |
| turquoise | 1515 | 0·84 | < 2·2e-16 |
| pink | 99 | 0·35 | 0·0004 |
| yellow | 321 | 0·28 | 2·27e-07 |
| purple | 79 | 0·70 | 7·23e-13 |
| brown | 1093 | 0·96 | < 2·2e-16 |
| cyan | 48 | 0·88 | < 2·2e-16 |
| tan | 71 | -0·29 | 0·0128 |
| greenyellow | 76 | 0·14 | 0·2270 |
| salmon | 67 | -0·27 | 0·0249 |
| magenta | 81 | -0·62 | 4·9e-10 |
| midnightblue | 45 | -0·50 | 0·0005 |
| black | 119 | -0·92 | < 2·2e-16 |
| lightcyan | 44 | -0·09 | 0·5660 |
| green | 272 | -0·87 | < 2·2e-16 |
| red | 126 | -0·67 | < 2·2e-16 |
| grey | 266 | -0·21 | 0·0006 |

**Table S2. Top 20 hub genes in module brown associated with bacterial burden**

| | Module eigengene | | | | GeneXpert Ct value | | | | Associated pathways |
|---|---|---|---|---|---|---|---|---|---|
| **Hub genes** | **N** | **Beta** | **95% CI** | **p-value** | **N** | **Beta** | **95% CI** | **p-value** | |
| CNIH4 | 295 | 0·16 | 0·15, 0·16 | <0·001 | 292 | -0·27 | -0·36, -0·19 | <0·001 | - |
| EIF4E3 | 295 | 0·16 | 0·15, 0·17 | <0·001 | 292 | -0·24 | -0·33, -0·14 | <0·001 | - |
| PHTF1 | 295 | 0·16 | 0·15, 0·16 | <0·001 | 292 | -0·22 | -0·31, -0·14 | <0·001 | - |
| BRI3 | 295 | 0·16 | 0·15, 0·17 | <0·001 | 292 | -0·20 | -0·29, -0·11 | <0·001 | TNF signaling pathway |
| DRAM1 | 295 | 0·15 | 0·14, 0·16 | <0·001 | 292 | -0·22 | -0·31, -0·13 | <0·001 | Toll-like receptor signaling pathway |
| CARD6 | 295 | 0·15 | 0·13, 0·16 | <0·001 | 292 | -0·22 | -0·31, -0·13 | <0·001 | NOD-like receptor signaling pathway |
| SIRPD | 295 | 0·15 | 0·13, 0·16 | <0·001 | 292 | -0·22 | -0·31, -0·13 | <0·001 | - |
| SERPINB1 | 295 | 0·15 | 0·14, 0·16 | <0·001 | 292 | -0·21 | -0·30, -0·12 | <0·001 | - |
| GNS | 295 | 0·15 | 0·14, 0·16 | <0·001 | 292 | -0·20 | -0·29, -0·11 | <0·001 | - |
| SEMA4C | 295 | -0·14 | -0·15, -0·12 | <0·001 | 292 | 0·16 | 0·07, 0·26 | <0·001 | - |
| NLRC3 | 295 | -0·15 | -0·16, -0·13 | <0·001 | 292 | 0·17 | 0·08, 0·27 | <0·001 | NOD-like receptor signaling pathway |
| BDH1 | 295 | -0·15 | -0·16, -0·13 | <0·001 | 292 | 0·17 | 0·07, 0·26 | <0·001 | - |
| ZAP70 | 295 | -0·14 | -0·15, -0·13 | <0·001 | 292 | 0·17 | 0·08, 0·26 | <0·001 | T cell receptor signaling pathway |
| ZNF831 | 295 | -0·14 | -0·15, -0·12 | <0·001 | 292 | 0·17 | 0·08, 0·25 | <0·001 | - |
| TRAF1 | 295 | -0·14 | -0·16, -0·13 | <0·001 | 292 | 0·17 | 0·08, 0·27 | <0·001 | TNF signaling pathway |
| IKZF3 | 295 | -0·13 | -0·15, -0·12 | <0·001 | 292 | 0·17 | 0·08, 0·26 | <0·001 | B cell receptor signaling pathway |
| MARCHF9 | 295 | -0·13 | -0·14, -0·12 | <0·001 | 292 | 0·17 | 0·08, 0·26 | <0·001 | - |
| PLCG1 | 295 | -0·13 | -0·14, -0·12 | <0·001 | 292 | 0·18 | 0·09, 0·26 | <0·001 | T cell receptor signaling pathway |
| SLC5A10 | 295 | -0·12 | -0·14, -0·11 | <0·001 | 292 | 0·18 | 0·09, 0·27 | <0·001 | - |
| STMN3 | 295 | -0·13 | -0·14, -0·12 | <0·001 | 292 | 0·20 | 0·11, 0·29 | <0·001 | - |

CI = Confidence Interval
